# Cone-driven retinal responses are shaped by rod but not cone HCN1

**DOI:** 10.1101/2021.11.03.467151

**Authors:** Colten K. Lankford, Yumiko Umino, Deepak Poria, Vladimir Kefalov, Eduardo Solessio, Sheila A. Baker

## Abstract

Signal integration of converging neural circuits is poorly understood. One example is in the retina where the integration of rod and cone signaling is responsible for the large dynamic range of vision. The relative contribution of rods versus cones is dictated by a complex function involving background light intensity and stimulus temporal frequency. One understudied mechanism involved in coordinating rod and cone signaling onto the shared retinal circuit is the hyperpolarization activated current (*I*_h_) mediated by HCN1 channels. *I*_h_ opposes membrane hyperpolarization driven by activation of the phototransduction cascade and modulates the strength and kinetics of the photoreceptor voltage response. We examined conditional knockout of HCN1 from rods using electroretinography. In the absence of HCN1, rod responses are prolonged in dim light which altered the response to slow modulation of light intensity both at the level of retinal signaling and behavior. Under brighter intensities, cone-driven signaling was suppressed. To our surprise, conditional knockout of HCN1 from cones had no effect on cone-mediated signaling. We propose that *I*_h_ is dispensable in cones due to the high level of temporal control of cone phototransduction. Thus, HCN1 is required for cone-driven retinal signaling only indirectly by modulating the voltage response of rods to limit their output.

**GRAPHICAL ABSTRACT:** 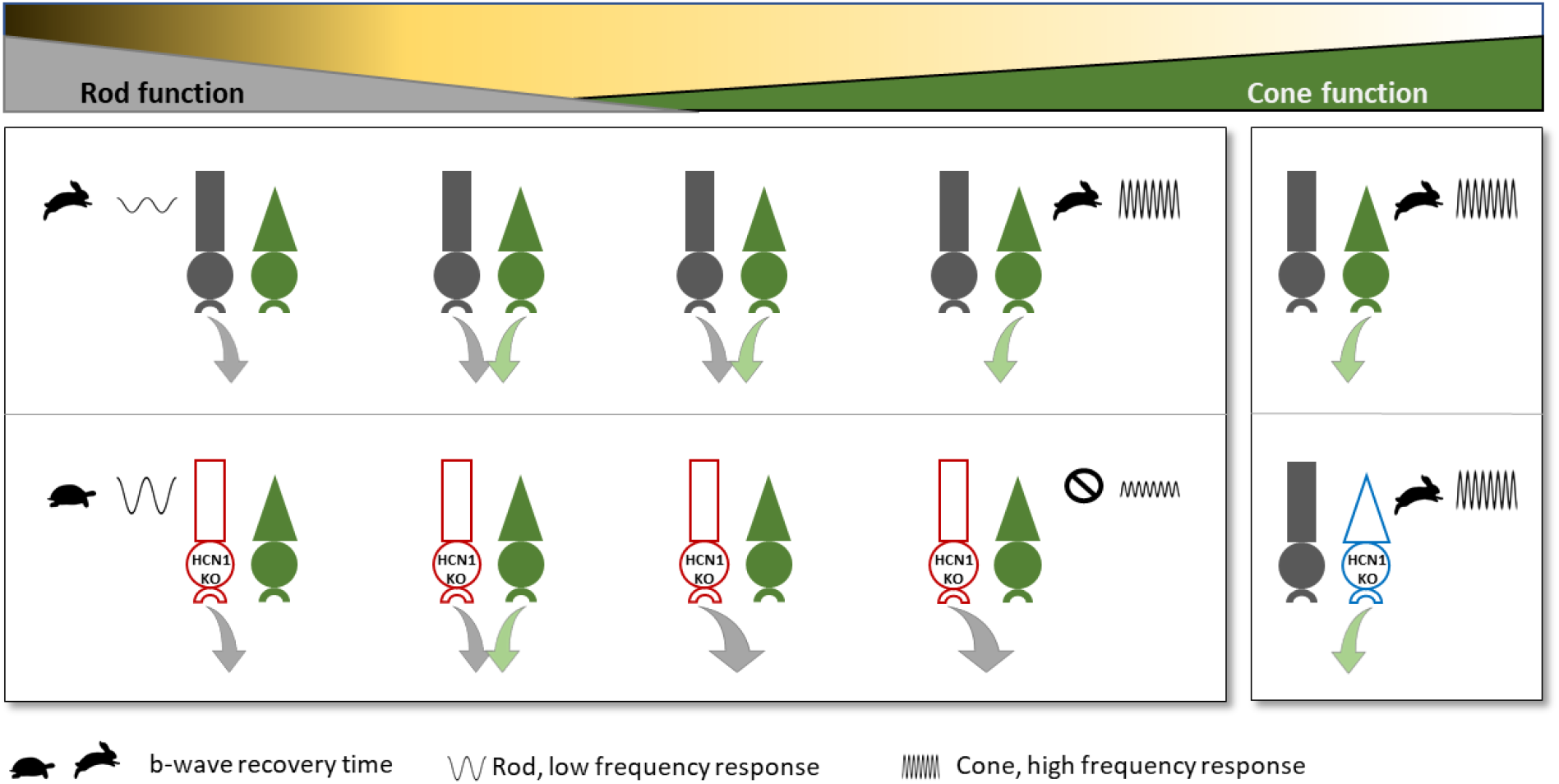

## INTRODUCTION

The exceptionally large dynamic range of vision depends on the coordinated activity of rod and cone photoreceptors (Lamb, 2016). Visual function flawlessly switches from purely rod driven (scotopic) to combined rod/cone driven (mesopic) and on to purely cone driven (photopic) as light intensity increases (Stockman and Sharpe, 2006; Zele and Cao, 2014). Rods and cones have highly interconnected circuits where rods, which evolved after cones, appear to have piggybacked onto the existing cone circuitry at multiple points (Demb and Singer, 2015; Fain and Sampath, 2018; Grimes et al., 2018; Masland, 2012; Völgyi et al., 2004). This raises the fundamental question as to how rod and cone output is controlled as vision transitions between these two systems given the downstream retinal circuitry can only carry a finite amount of information.

Two independent processes that control rod responses are light adaptation of the photoresponse and modulation of the voltage response. Light adaptation consists of multiple mechanisms operating at the level of the rod phototransduction cascade to reduce sensitivity as background light increases or persists (Arshavsky and Burns, 2012). This results in rod-driven responses with reduced magnitude but faster temporal properties and effectively limits the output from rods until rods reach saturation and no longer contribute to vision (Naarendorp et al., 2010). While the mechanisms and consequences of classical adaptation are well described, the role that voltage modulation plays in controlling rod output is largely speculative.

The voltage response of photoreceptors is modulated primarily by two currents originating from photoreceptor inner segments: *I*_h_ and *I*_kx_ carried by hyperpolarization activated cyclic nucleotide gated 1 (HCN1) and heteromeric Kv2.1/Kv8.2 channels respectively (Beech and Barnes, 1989; Czirjak et al., 2007; Fain et al., 1978; Fain and Sampath, 2021; Knop et al., 2008). These currents have opposing activity with *I*_kx_ being an outward current active under depolarized potentials and *I*_h_ being an inward current active under hyperpolarized potentials. *I*_kx_ acts to establish the resting membrane potential, accounting for ~70% of the outward component of the dark current (Fortenbach et al., 2021). In response to a brief flash of light *I*_kx_ inactivation and *I*_h_ activation quickly drive the membrane potential back toward the depolarized dark potential (Barrow and Wu, 2009; Beech and Barnes, 1989). What remains unclear is how HCN1 activation in response to prolonged or temporally dynamic light stimuli affects rod and cone output to the shared downstream retinal circuitry.

The expectation is that by accelerating the kinetics of the rod voltage response, HCN1 activity facilitates rod responses to rapidly modulated stimuli. This is supported by electroretinography (ERG) studies using HCN1 knockout (KO) mice which concluded that HCN1 amplifies the response to high frequency stimulation under bright but not dim conditions (Knop et al., 2008; Seeliger et al., 2011; Sothilingam et al., 2016). However, this contrasts with direct rod recordings which concluded that HCN1 acts to attenuate the magnitude of the voltage response to low frequency stimulation but has no apparent impact on the voltage response to high frequency stimulation (Barrow and Wu, 2009; Della Santina et al., 2012). It is not clear how to reconcile these observations.

Under photopic conditions, where cones drive the ERG response, HCN1 KO mice were reported to have responses that were reduced in amplitude but prolonged. That report contrasts with the expectation derived from voltage recordings of salamander cones where pharmacological block of *I*_h_ did not prolong the cone voltage response but enhanced the magnitude of light induced hyperpolarization (Barrow and Wu, 2009). It is difficult to directly compare these two studies because they used different species and techniques, but the differing conclusions highlight the state of uncertainty regarding the role HCN1 plays in cones. Cone function under mesopic conditions was reported to require rod expressed HCN1 to prevent rods from saturating the retinal circuit (Seeliger et al., 2011). It is not known if HCN1 serves a similar function under photopic conditions.

In all prior studies of HCN1 in the retina, interpretation is confounded by the fact that HCN1 is expressed in multiple cell types (Knop et al., 2008; Müller et al., 2003). To directly assess the role of HCN1 in photoreceptors under different lighting conditions, we generated rod and cone specific HCN1 KO lines. The animals were examined with a battery of ERG tests. Contrary to expectations, we found that selective ablation of HCN1 in rods did not limit responses to high frequency mesopic flickering light but instead significantly shaped the responses to low frequency flicker. We confirmed this at the behavioral level using optomotor response (OMR) assays where loss of HCN1 in rods resulted in reduced contrast sensitivity at low temporal frequencies. There was a major impact on cone-driven ERG responses in the rod-specific HCN1 KO under high mesopic conditions, as predicted by Seeliger et al., and under photopic conditions where rods are not thought to contribute to vision. Most surprising to us, ablation of HCN1 in cones did not alter retina function.

## RESULTS

### Generation of HCN1-Rod KO mice

To ablate HCN1 from rods, a previously characterized conditional HCN1 KO line was crossed to the Rho-iCre (also known as iCre-75) line to generate littermate control HCN1^fl/fl^:Rho-iCre^−^ (Rod-Control) and experimental animals, HCN1^fl/fl^:Rho-iCre^+^ (Rod-HCN1 KO). Some colonies of Rho-iCre are contaminated by a transgene resulting in overexpression of R9AP (Sundermeier et al., 2014). However, we obtained our mice from the Jackson Lab which do not have this contaminating transgene and we independently verified that using PCR genotyping (data not shown). Cre expression from transgenes can exhibit variable mosaicism. Using immunohistochemistry, Cre expression was absent in the Rod-Control (Fig. 1A) but was present in the majority of rod nuclei in the ONL of the Rod-HCN1 KO (Fig. 1B). Co-labeling with HCN1 and cone arrestin revealed HCN1 signal throughout the inner segment and the membranes of the outer nuclear layer in the Rod-Control (Fig. 1C) while HCN1 signal was restricted to Cone Arrestin positive cones in the Rod-HCN1 KO (Fig. 1D). This validation was conducted on two-month-old mice and all subsequent experiments used mice of this age or older.

**Figure 1:**
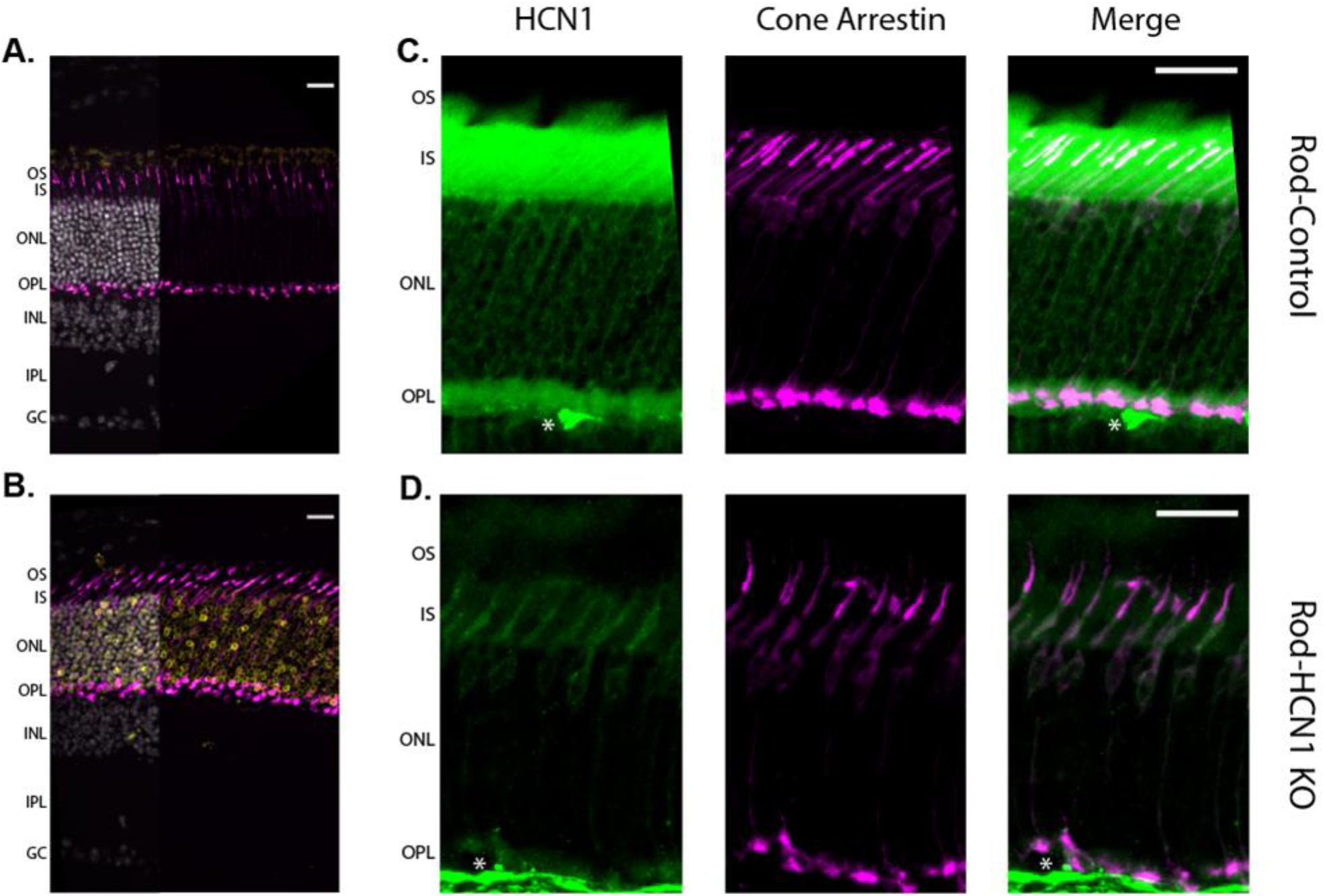
Validation of Rod-HCN1 KO line. Low magnification view of retina from **A)** Rod-Control or **B)** Rod-HCN1 KO immunolabeled for Cre (yellow) and Cone Arrestin (magenta). Overlay with Hoechst labeled nuclei (grey) only partially shown to increase visibility of Cre labeling which is detected only in the Rod-HCN1 KO. Higher magnification view of the photoreceptor layer from **C)** Rod-Control or **D)** Rod-HCN1 KO immunolabeled for HCN1 (green) and Cone Arrestin (magenta). HCN1 staining is cone-specific in the Rod-HCN1 KO. Both scale bars are 20 μm. Abbreviations are OS, outer segment; IS, inner segment; ONL, outer nuclear layer; OPL, outer plexiform layer; INL, inner nuclear layer; IPL, inner plexiform layer; GC, ganglion cell layer. Asterisks (*) are blood vessels non-specifically stained with secondary antibody.

### Rod-HCN1 KO mice have a prolonged rod-driven b-wave

A characteristic rod-driven phenotype of HCN1 KO mice is the prolonged scotopic b-wave. To validate that the HCN1-Rod KO mice exhibit this characteristic response, we recorded ERGs from dark-adapted Rod-HCN1 KO and Rod-Control mice stimulated with brief flashes of light from 0.003 cd.s/m^2^ to 100 cd.s/m^2^ (Fig. 2A). The ERG response under these conditions has an initial negative inflection termed the a-wave followed by a larger, positive inflection termed the b-wave. The a-wave is generated by activation of the phototransduction cascade and reflects the initial phase of the photoreceptor voltage response. As such, the a-wave is unlikely to be affected by loss of HCN1. We observed an increase in the HCN1-Rod KO a-wave amplitude (Fig. 2B – dashed lines; ANOVA, p = 0.0046), with no change in the time to peak (Fig. 2C – dashed lines; ANOVA, p = 0.0773). The difference in a-wave amplitude between HCN1-KO and controls was very small and multiple comparisons test revealed differences only at 0.01, 0.1, and 3 cd.s/m^2^ (ANOVA, p = 0.0019, 0.0212, and 0.0250). This slight alteration in the a-wave did not translate to an altered b-wave as the amplitude (Fig. 2B – solid lines) and time to peak (Fig. 2C – solid lines) for the b-wave, derived in large part from synaptic transmission and subsequent activation of bipolar cells, was unaltered in the Rod-HCN1 KO (ANOVA p = 0.6904 for amplitude and 0.1231 for time to peak).

**Figure 2:**
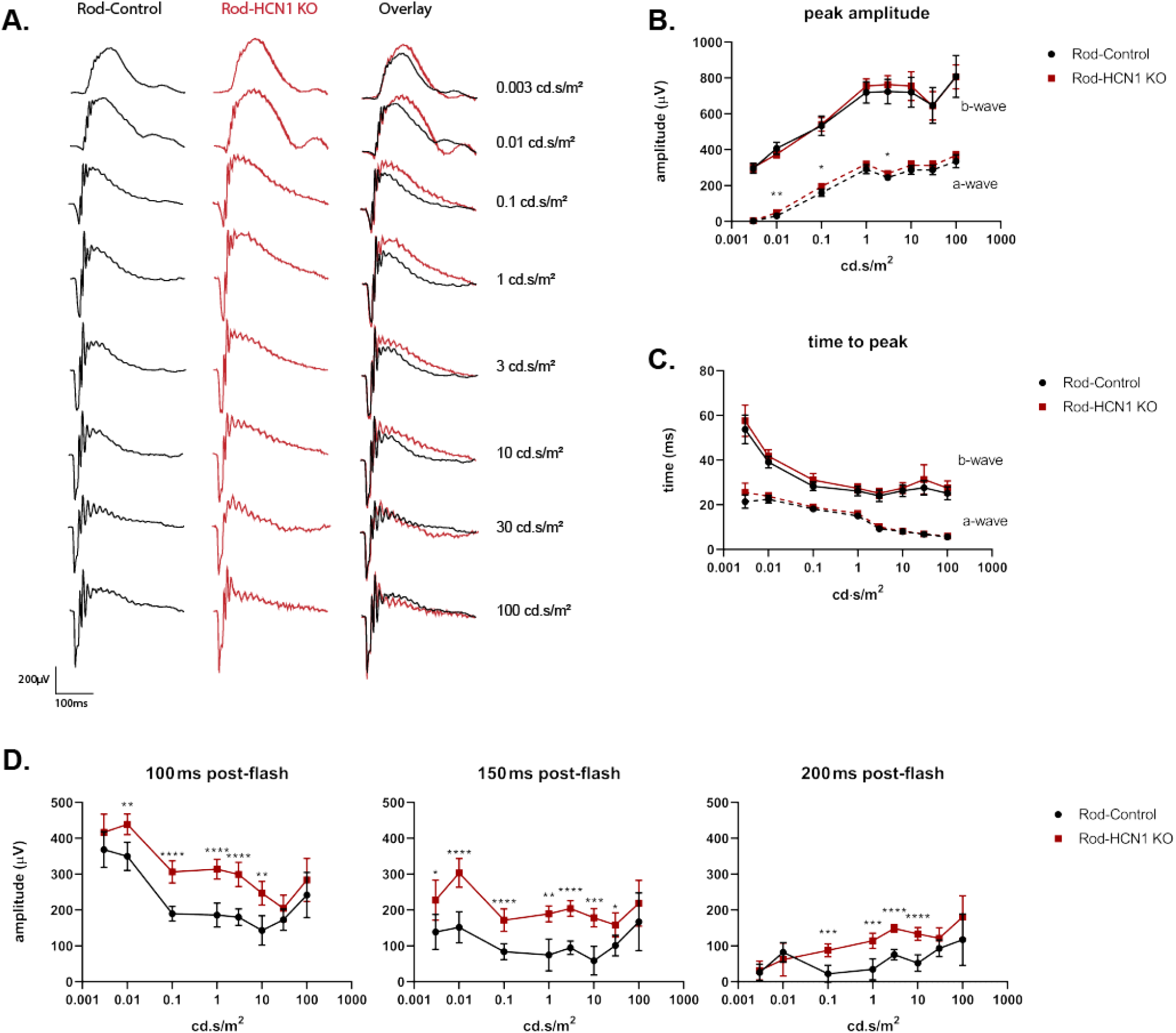
Dark adapted ERG: Rod-HCN1 KO. **A)** Representative family of ERG traces from dark adapted Rod-Control (black) and Rod-HCN1 KO (red) mice following a flash at the given intensity. **B)** Amplitude of a-wave (dashed line) and b-wave (solid line) plotted against stimulus intensity. **C)** Time to peak of a-wave (dashed line) and b-wave (solid line) plotted against stimulus intensity. **D)** Amplitude of the b-wave relative to baseline at 100, 150, or 200 ms after the flash; p <0.0001 for difference between genotypes at all three timepoints. Data is presented as mean ± SD. For sample size and detailed statistics see Supplemental Table 2.1-2.7.

While the initial phase of the b-wave was identical between Rod-Control and Rod-HCN1 KO, recovery of the b-wave was delayed in the Rod-HCN1 KO. To examine this delay, we quantified the amplitude of the descending b-wave from baseline at 100, 150, and 200 ms after the flash. The b-wave amplitude was consistently larger in the Rod-HCN1 KO at all three timepoints (Fig. 2D; ANOVA p < 0.0001 for all three timepoints). At the dimmest flash, 0.003 cd.s/m^2^, the b-wave elevation was only statistically significant at 150 ms (adj p = 0.0357). Likely because at this intensity, the b-wave is slower and shorter in duration than at brighter intensities. Note that we measure the b-wave amplitude as the second oscillatory potential peak on the rising b-wave. The prolonged b-wave is not apparent at the highest flash intensities where the contribution from cone-driven signaling is increased. Together, these results are consistent with observations in the global HCN1 KO mouse and confirm that HCN1 in rods is required for rapid rod voltage recovery (Knop et al., 2008).

### Rod-HCN1 KO mice can respond to a high frequency sinusoidal flicker under scotopic and low mesopic conditions

To examine the temporal properties of HCN1-dependent signaling we used a flicker ERG protocol with a sinusoidal stimulus ranging from 0.5-30 Hz. For scotopic conditions, a mean background illumination of 0.05 cd/m^2^ (equating to 40 R*/rod/s) was used. Both Rod-Control and Rod-HCN1 KO mice generated responses with similarly shaped waveforms (Fig. 3A). To quantify the response to the sinusoidal flicker we measured the amplitude of the response and plotted this as a function of stimulus frequency. This revealed a band-pass pattern for both Rod-Control and Rod-HCN1 KO responses (Fig. 3B). The Rod-HCN1 KO mice trended towards having a larger amplitude (ANOVA p = 0.0040), which by multiple comparisons testing was only significant at 0.5 Hz (adj p = 0.0074). Traditionally, the sinusoidal flicker response is analyzed using the fast Fourier transform to isolate the fundamental (F0), or higher harmonic, components. When we applied this analysis, we observed a similar bandpass pattern with a trend towards increased response magnitude in the Rod-HCN1 KO (Fig. S1A).

**Figure 3:**
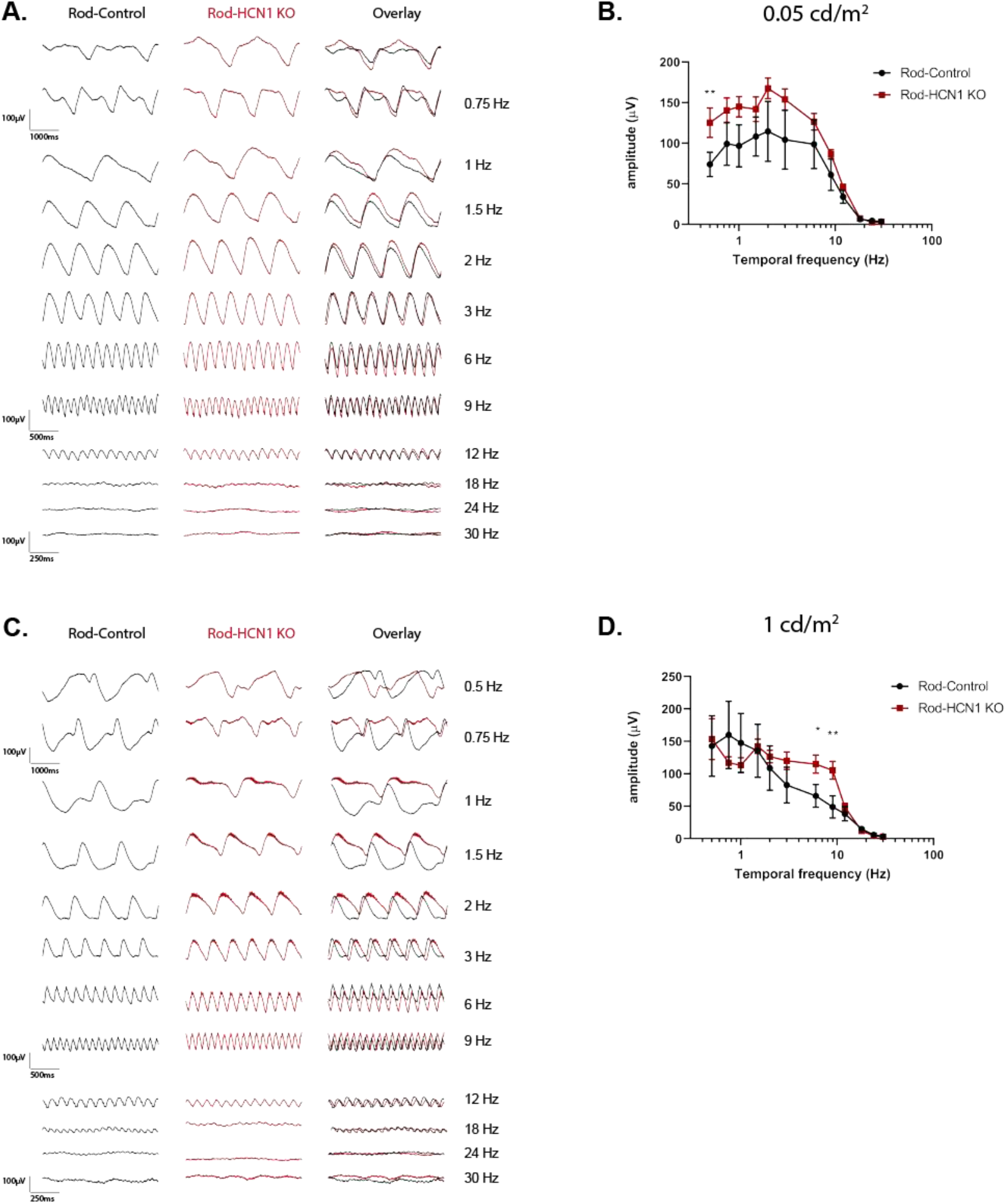
Low light sinusoidal flicker ERG: Rod-HCN1 KO. Representative family of ERG traces from Rod-Control (black) and Rod-HCN1 KO (red) mice using a sinusoidal flicker at the given frequency with a mean illumination of 0.05 cd/m^2^ **(A)** or 1 cd/m^2^ **(D)**. **B)** Amplitude of the responses from the data shown in (A) plotted against stimulus frequency; difference between genotypes is p = 0.0040. **D)** Data presented in (B) normalized to mean amplitude of Rod-Control. **E)** Amplitude of the responses from the data shown (D) plotted against stimulus frequency; difference between genotypes p = 0.3874. **F)** Data presented in (E) normalized to mean amplitude of Rod-Control. Data is presented as mean ± SD. For sample size and detailed statistics see Supplemental Tables 3.1-3.2.

When the mean illumination was increased to 1 cd/m^2^ (equating to 800 R*/rod/s) to test mesopic conditions, the shape of the response waveform generated by the Rod-HCN1 KO mice differed from Rod-Control at frequencies less than 2 Hz but became similar at higher frequencies (Fig. 3D). When we measured the response amplitude as a function of frequency there was no difference between genotypes (ANOVA, p = 0.3874). Because of the difference in waveform shape at low versus high frequency we examined the multiple comparisons test. This indicated that the Rod-HCN1 KO responses were significantly elevated but only from 6-9 Hz (adj p = 0.0135 and 0.0047, respectively). Calculation of F_0_ generated a similar result as obtained analyzing the untransformed amplitude (Fig. S1B).

The waveform below 2 Hz was not sinusoidal suggesting the possibility of non-linear responses at low frequencies. To take that into account we previously characterized the sinusoidal flicker response to low frequency waveforms as being composed of “b-wave-like” and “c-wave-like” components with a large recovery phase between the two (Inamdar et al., 2021). Applying that characterization here, the altered response waveform in the HCN1-Rod KO mice appeared consistent with these two components merging, as would be expected from a prolonged b-wave. We measured the magnitude and rate of the recovery following the b-wave-like component as noted in Fig. S1C. The magnitude of the voltage recovery following the b-wave-like peak was greatly reduced (ANOVA p < 0.0001) and exhibited a slowed decay rate (ANOVA p < 0.0001) in the Rod-HCN1 KO mice (Fig. S1D, E). Thus, we conclude that HCN1 in rods shapes the ERG response to low frequency stimulation in a mean illumination dependent manner but does not extend the temporal resolution of the retina as Rod-HCN1 KO mice were able to response to high frequency flicker under scotopic and low mesopic conditions.

#### Rod-HCN1 KO mice have suppressed sinusoidal flicker responses under high mesopic and photopic conditions

It was previously reported that HCN1 KO impaired cone-driven signaling to a high frequency mesopic flicker. We repeated the sinusoidal flicker assay but increased the mean illumination to high mesopic (10 cd/m^2^, estimated 8,000 R*/rod/s) and photopic conditions (30 cd/m^2^, estimated 24,000 R*/rod/s) (Fig. 4). The Rod-HCN1 KO mice were able to respond to the stimulus but in both cases, there was a trend for the Rod-HCN1 KO response amplitudes being reduced compared to Rod-Controls at all frequencies higher than 0.5 Hz (ANOVA, p = 0.0229 for the high mesopic or p = 0.0097 for the photopic test; Fig. 4B and D). Calculation of F_0_ generated a similar result as obtained analyzing the untransformed amplitude (Fig. S2).

**Figure 4:**
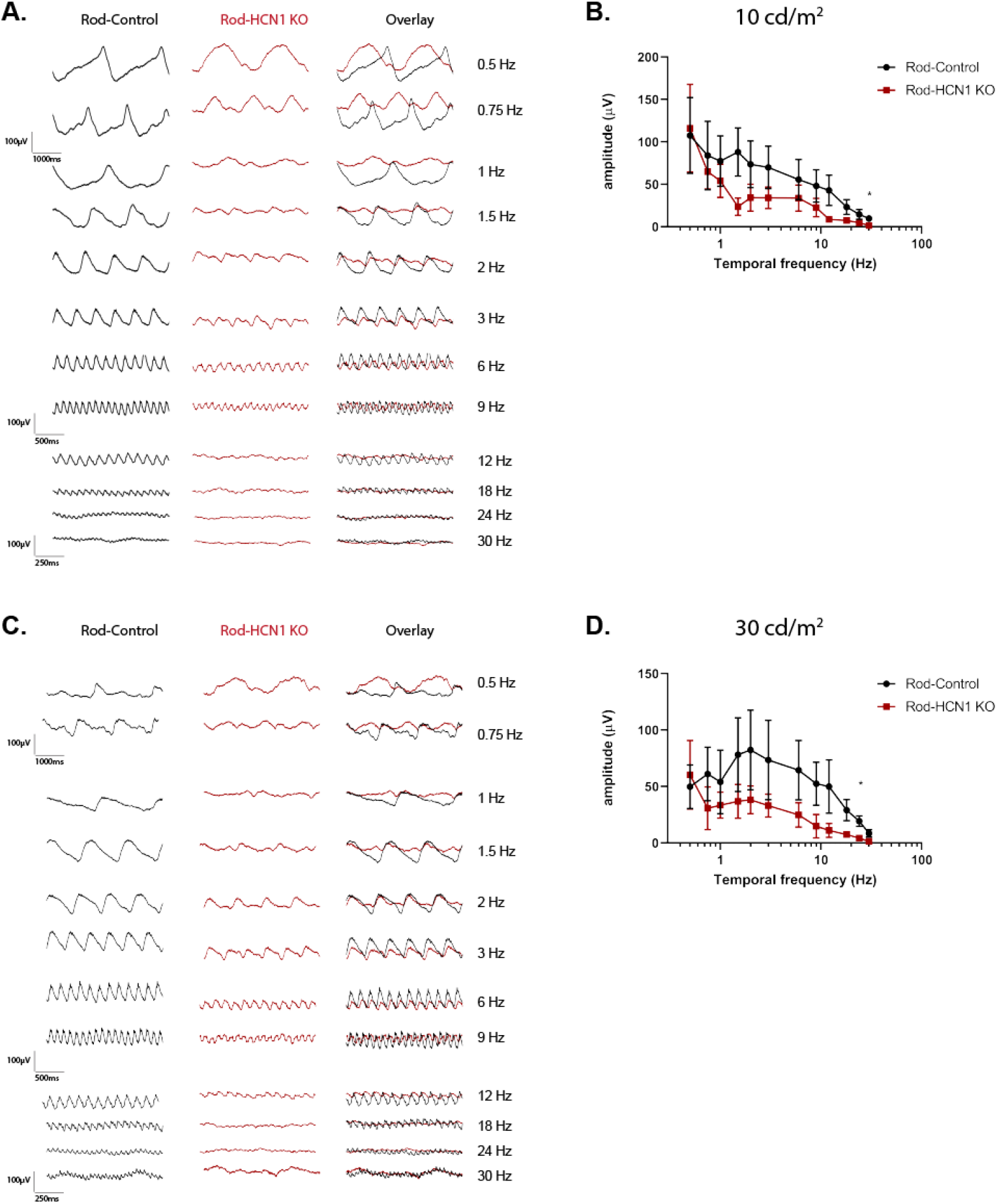
Bright sinusoidal flicker ERG: Rod-HCN1 KO. Representative family of ERG traces from Rod-Control (black) and Rod-HCN1 KO (red) mice using a sinusoidal flicker at the given frequency with a mean illumination of 10 cd/m^2^ **(A)** or 30 cd/m^2^ **(D)**. **B)** Amplitude of the responses from the data shown in (A) plotted against stimulus frequency; difference between genotypes is p = 0.0229. **C)** Data presented in (B) normalized to mean amplitude of the Rod-Control. **E)** Amplitude of the responses from the data shown (D) plotted against stimulus frequency; difference between genotypes p = 0.0097. **F)** Data presented in (E) normalized to mean amplitude of the Rod-Control. Data is presented as mean ± SD. For sample size and detailed statistics see Supplemental Tables 4.1-4.2.

It was previously reported that whole-body HCN1 KO mice have a more drastic inability to respond to mesopic flicker (Seeliger et al., 2011). That study used a flash flicker ERG protocol that results in an effective increase in average background light as the frequency increases that could lead to light-adaptation of rods during the experiment. To determine if the different responses to mesopic flicker were due to the mice or the protocol used, we tested the Rod-HCN1 KO mice using the same flash flicker as described by Seeliger and colleagues. Dark adapted Rod-HCN1 KO mice were stimulated with a repetitive 3 cd.s/m^2^ flash at frequencies ranging from 0.5 to 30 Hz (Fig. 5) At 0.5 Hz, the Rod-HCN1 KO mice generated a similar response amplitude as controls (adj p > 0.9999), though with a prolonged b-wave recovery phase. At 1 Hz, the b-wave was suppressed (adj p = 0.0067) and there was effectively no response from the HCN1-Rod KO mice at higher frequencies (Fig. 5B; ANOVA p < 0.0001), in agreement with the results obtained using the whole-body HCN1 KO (Seeliger et al., 2011). An interesting additional benefit of using this protocol is that it is known that rods are the primary driver of the ERG response up to 3 Hz at which point cones drive the response primarily through the ON-bipolar cells (3-15 Hz) then OFF-bipolar cells (>15 Hz) (Tanimoto et al., 2009).

**Figure 5:**
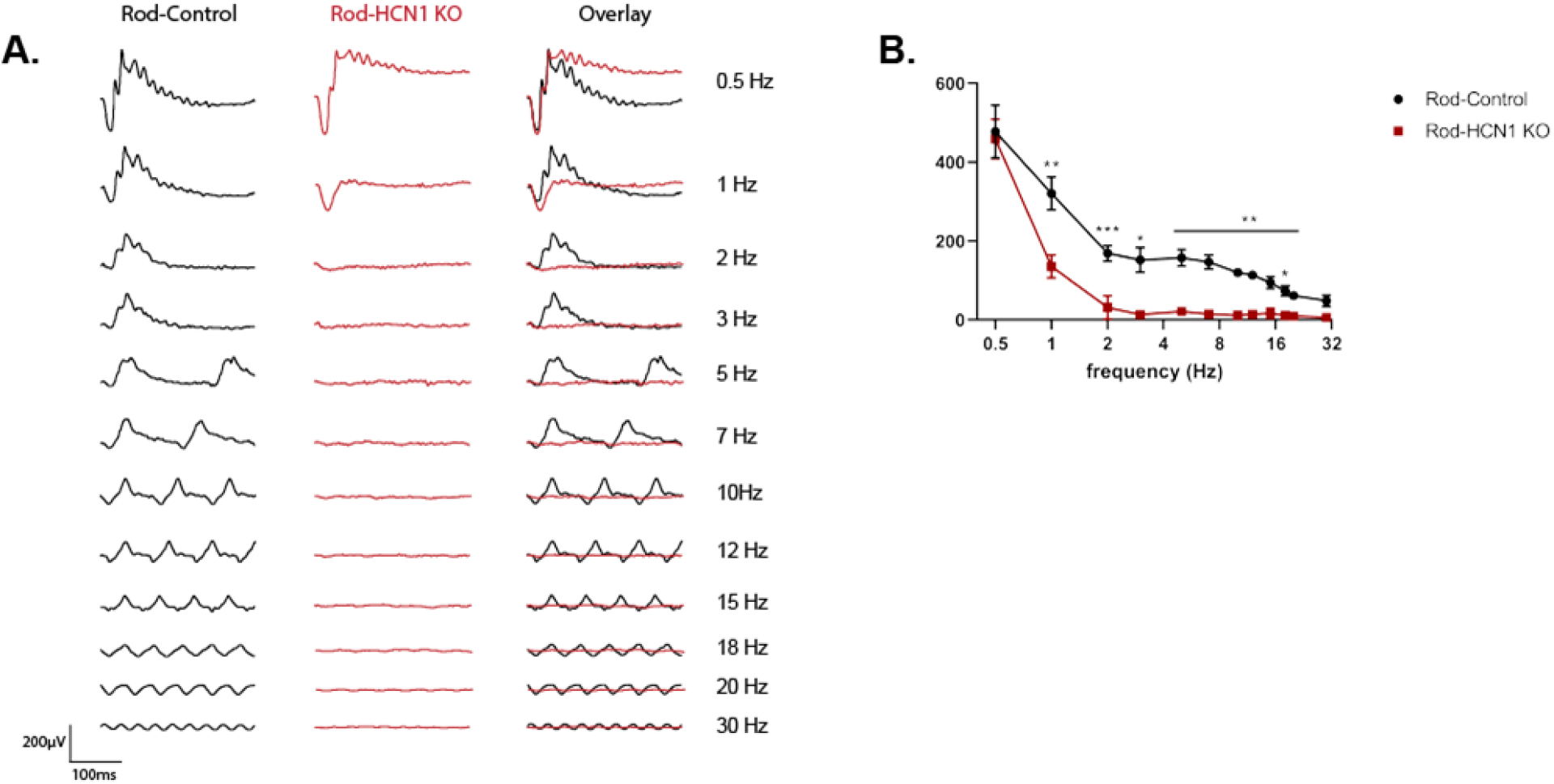
Flash flicker ERG: Rod-HCN1 KO. **A)** Representative family of ERG traces from dark adapted Rod-Control (black) and Rod-HCN1 KO (red) mice following a 3 cd.s/m^2^ flash flicker at the given frequency. **B)** Response amplitude plotted against frequency; p <0.0001 for difference between genotypes. Data is presented as mean ± SD. For sample size and detailed statistics see Supplemental Table 5.

### Rod-HCN1 KO mice have no photopic b-wave

The flicker ERG tests indicated cone dysfunction. To further examine cone-driven signaling in the Rod-HCN1 KO mice, we examined the light-adapted (photopic) single flash ERG response of these mice. The single flash response is well characterized with the a-wave reflecting cone activation and the b-wave reflecting cone-driven activation of downstream neurons, primarily ON-bipolar cells. The photopic ERG was obtained by stimulating light-adapted Rod-Control and Rod-HCN1 KO mice with flashes of light ranging from 0.23 cd.s/m^2^ to 724 cd.s/m^2^ in the presence of a rod-saturating background light of 30 cd/m^2^ (Fig. 6). Despite having a normal a-wave amplitude (Fig. 6B; ANOVA p = 0.6359), the Rod-HCN1 KO mice did not exhibit a photopic b-wave at any intensity tested (Fig. 6C; ANOVA p < 0.0001). This indicates that under rod saturating conditions, cones are functional but unable to propagate signals onto bipolar cells when HCN1 is lost in rods. While this phenotype is more severe than what was reported in the global HCN1 KO (Knop et al., 2008), it is consistent with the global HCN1 KO having a suppressed photopic b-wave and is consistent with our flicker ERG data demonstrating that loss of HCN1 in rods impairs cone-driven signaling.

**Figure 6:**
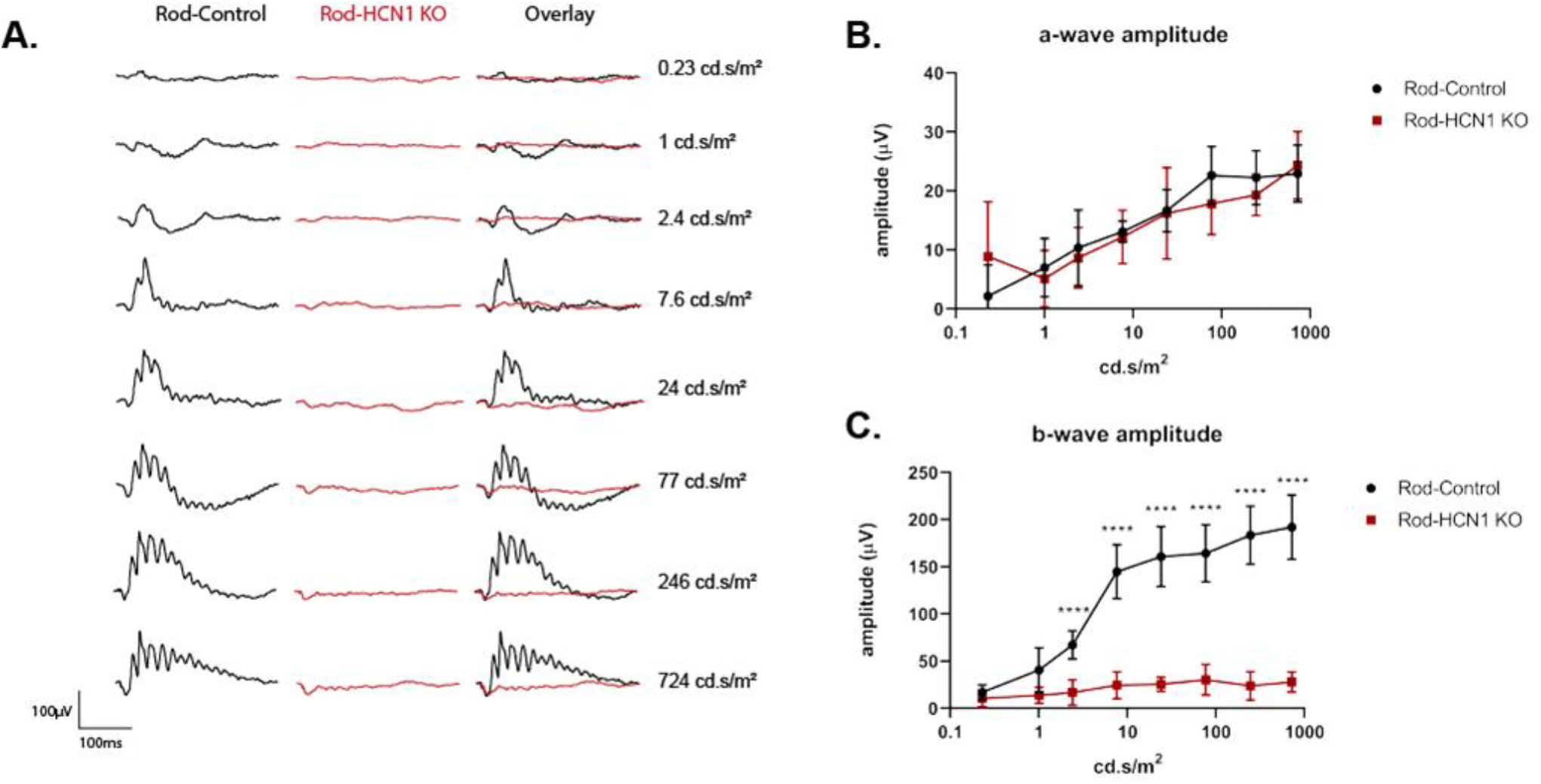
Light-adapted ERG: Rod-HCN1 KO. **A)** Representative family of ERG traces from light (30 cd/m^2^) adapted Rod-Control (black) and Rod-HCN1 KO (red) mice following a flash at the given intensity. **B)** Amplitude of a-wave plotted against stimulus intensity; p = 0.6359. **C)** Amplitude of b-wave plotted against stimulus intensity; p <0.0001 for difference between genotypes. Data is presented as mean ± SD. For sample size and detailed statistics see Supplemental Tables 6.1-6.2.

In conclusion, selective ablation of HCN1 in rods leads to prolonged rod-driven ERG responses that suppress transmission of cone-driven signaling. This confirms the prevailing model for HCN1 function in rods and unexpectedly extends the phenomenon from the mesopic range into photopic conditions.

#### LanBehavioral responses of Rod-HCN1 KO mice

To examine how loss of HCN1 in rods impacts vision beyond summed retinal electrical patterns, we used the optomotor response, a visually dependent innate reflexive response, to measure the temporal contrast sensitivity of Rod-HCN1 KO.

We examined Rod-HCN1 KO mice under dim conditions using a 0.002 cd/m^2^ background (producing an estimated 1.4 R*/rod/s) and a 0.031 c/d spatial frequency at 0.1, 0.4, and 1.5 Hz. We could not use higher frequencies due to spatiotemporal limitations in the Optomotry system. Under dim conditions, we do not expect rods to be driven to sufficient hyperpolarized potentials to significantly activate HCN1 channels. Consistent with this prediction, the temporal contrast sensitivity function for Rod-HCN1 KO and Rod-Control mice were identical (Fig. 7A; p = 0.377). We then repeated this experiment under brighter conditions, 70 cd/m^2^ background (1500 R*/rod/s), where HCN1 channels would likely play a role shaping the rod voltage response. We observed a reduction in the contrast sensitivity of the Rod-HCN1 KO at 0.1 Hz (adj p = 0.005) but no difference at 0.4 and 1.5 Hz frequencies (Fig. 7B; p = 0.23). Thus, we see that HCN1 channels in rods impact contrast sensitivity at low temporal frequencies in a background illumination dependent manner.

**Figure 7:**
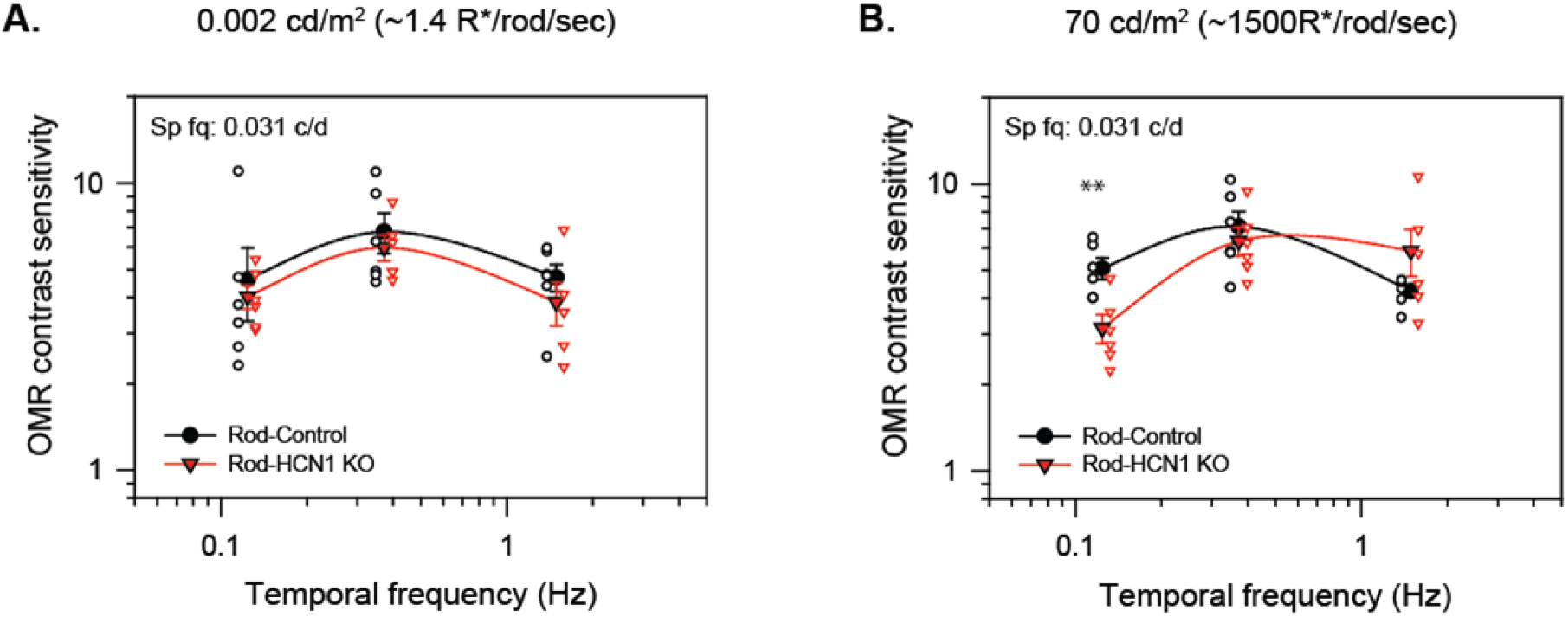
Optomotor temporal contrast sensitivity: Rod-HCN1 KO. **A)** Contrast sensitivity of Rod-Control (black) and Rod-HCN1 KO (red) mice under scotopic conditions with a background light of 0.002 cd/m^2^ (estimated 1.4 R*/rod/s) or **B)** under mesopic conditions with a background light of 70 cd/m^2^ (estimated 1500 R*/rod/s) and a spatial frequency of 0.031 cycles/degree. Animals were tested at frequencies of 0.1, 0.4, and 1.5 Hz. The only significant difference between genotypes was at 0.1 Hz under mesopic conditions p = 0.005 (**). Data is presented as mean ± SEM with each individual mouse average shown. For sample size and detailed statistics see Supplemental Tables 7.1-7.2.

#### Generation of Cone-HCN1 KO mice

To ablate HCN1 from cones, the conditional HCN1 KO line was crossed to the cone-specific HRGP-Cre to generate littermate control HCN1^fl/fl^:HRGP-Cre^−^ (Cone-Control) and experimental HCN1^fl/fl^:HRGP-Cre^+^ (Cone-HCN1 KO) animals. Cre was not detected in Cone-Control retina (Fig. 8A) and Cre expression in Cone-HCN1 KO was restricted to cones (Fig. 8D), verified by double labeling for Cone Arrestin. HCN1 labeling throughout the inner segment and outer nuclear layers confirmed HCN1 expression in rods of Cone-HCN1 KO mice (Fig. 8B, E). An absence of HCN1 in cones was not immediately obvious by visual examination, but this is likely due to the abundance of rods in the murine retina and the proximity of rod and cone inner segment plasma membranes. We measured the fluorescence intensity along the length of short lines (~10 μm long) drawn laterally across the inner segments. In Cone-Control retinas, the intensity of HCN1 signal was similar in Cone Arrestin positive (cone) and negative (rod) areas (Fig. 8C). In Cone-HCN1 KO, there was a small decrease in HCN1 signal where the cone arrestin signal peaked (Fig. 8F)

**Figure 8:**
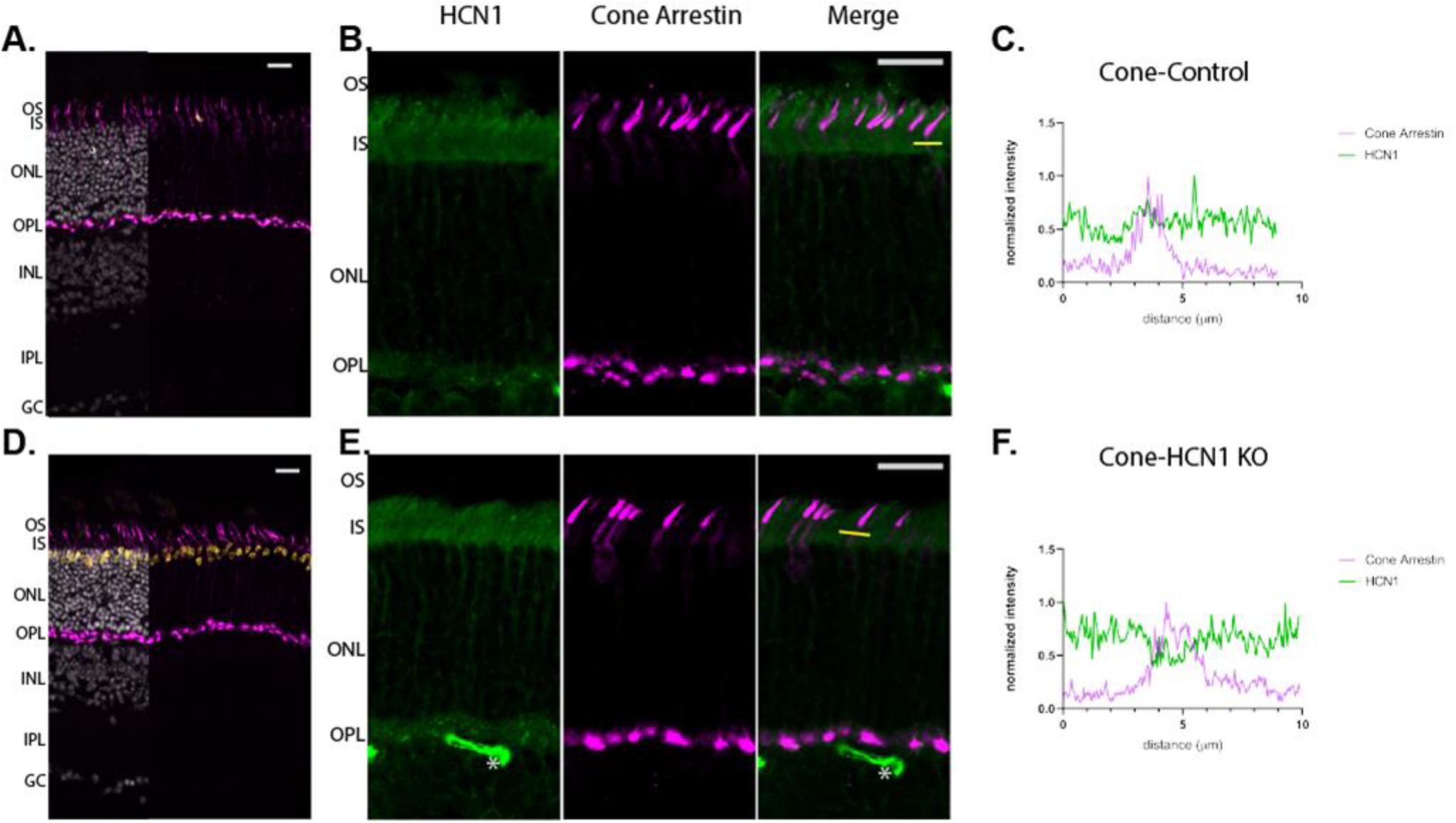
Validation of Cone-HCN1 KO line. Low magnification view of retina from **A)** Cone-Control or **D)** Cone-HCN1 KO immunolabeled for Cre (yellow) and Cone Arrestin (magenta). Overlay with Hoechst labeled nuclei (grey) only partially shown to increase visibility of Cre labeling which is detected only in the Cone-HCN1 KO. Higher magnification view of the photoreceptor layer from **B)** Cone-Control or **E)** Cone-HCN1 KO immunolabeled for HCN1 (green) and Cone Arrestin (magenta). HCN1 staining is detected in rods and largely obscures cones. Line scan analysis of **C)** Cone-Control or **F)** Cone-HCN1 KO; the normalized cone arrestin and HCN1 signal as a function of distance along the line shown in **C** or **F**, respectively. Both scale bars are 20 μm. Abbreviations are OS, outer segment; IS, inner segment; ONL, outer nuclear layer; OPL, outer plexiform layer; INL, inner nuclear layer; IPL, inner plexiform layer; GC, ganglion cell layer. Asterisks (*) are blood vessels non-specifically stained with secondary antibody.

To get a more representative assessment of HCN1 expression in cones, we repeated this analysis using longer lines (~100 μm) across the inner segment to transect several cones (Fig. S3A, B). HCN1 signal intensity was plotted against cone arrestin intensity for each pixel and compared the correlation between the two signals using Pearson’s correlation (Fig. S3C, D). This was repeated across four separate retinas for each genotype. There was a consistent reduction in the correlation between HCN1 and cone arrestin staining intensity which would be expected when HCN1 protein level is reduced in cones (Fig. S3E; Cone-Control: r = 0.11 ± 0.01 vs Cone-HCN1 KO: r = −0.067 ± 0.077; t-test p = 0.0040). This analysis demonstrates a reliable reduction of HCN1 protein in cones. Animals were two-months-old and all subsequent experiments used mice of this age or older.

#### Cone-HCN1 KO mice have unaltered ERG responses

To examine the extent to which cone expressed HCN1 contributes to the ERG response we performed the same set of ERG tests as described for the Rod-HCN1 KO. We began with the dark-adapted flash series of increasing light intensity to ensure rods responses were normal. As expected, there were no differences between Cone-Control and Cone-HCN1 KO animals at the lower flash intensities (Fig. S4). Nor was there a difference at higher light intensities where cones contribute to the response waveform. To probe cone function directly, animals were light adapted and flashes delivered on a 30 cd/m^2^ background light to saturate rods and minimize their contribution to the response.

The Cone-Control and Cone-HCN1 KO mice exhibited an identical ERG response at all flash intensities tested (Fig. 9). Quantitation of a-wave and b-wave amplitudes (Fig. 9B; ANOVA a-wave p = 0.5257; b-wave p = 0.2928) and implicit times (Fig. 9C; ANOVA a-wave p = 0.0953; b-wave p = 0.2928) were the same. Despite expectations from the whole-body HCN1 KO, the b-wave was not prolonged in the Cone-HCN1 KO mice as b-wave amplitudes were identical to Cone-Controls at 100 or 150 ms after the flash (Fig. 9D; ANOVA p = 0.7315 and 0.5606). At 200 ms after the flash there was an apparent statistically significant difference (ANOVA p = 0.0105), but multiple comparisons test revealed no difference in amplitude at any individual intensity tested. The apparent difference may be due to the low signal-to-noise ratio at that point in the b-wave recovery.

**Figure 9:**
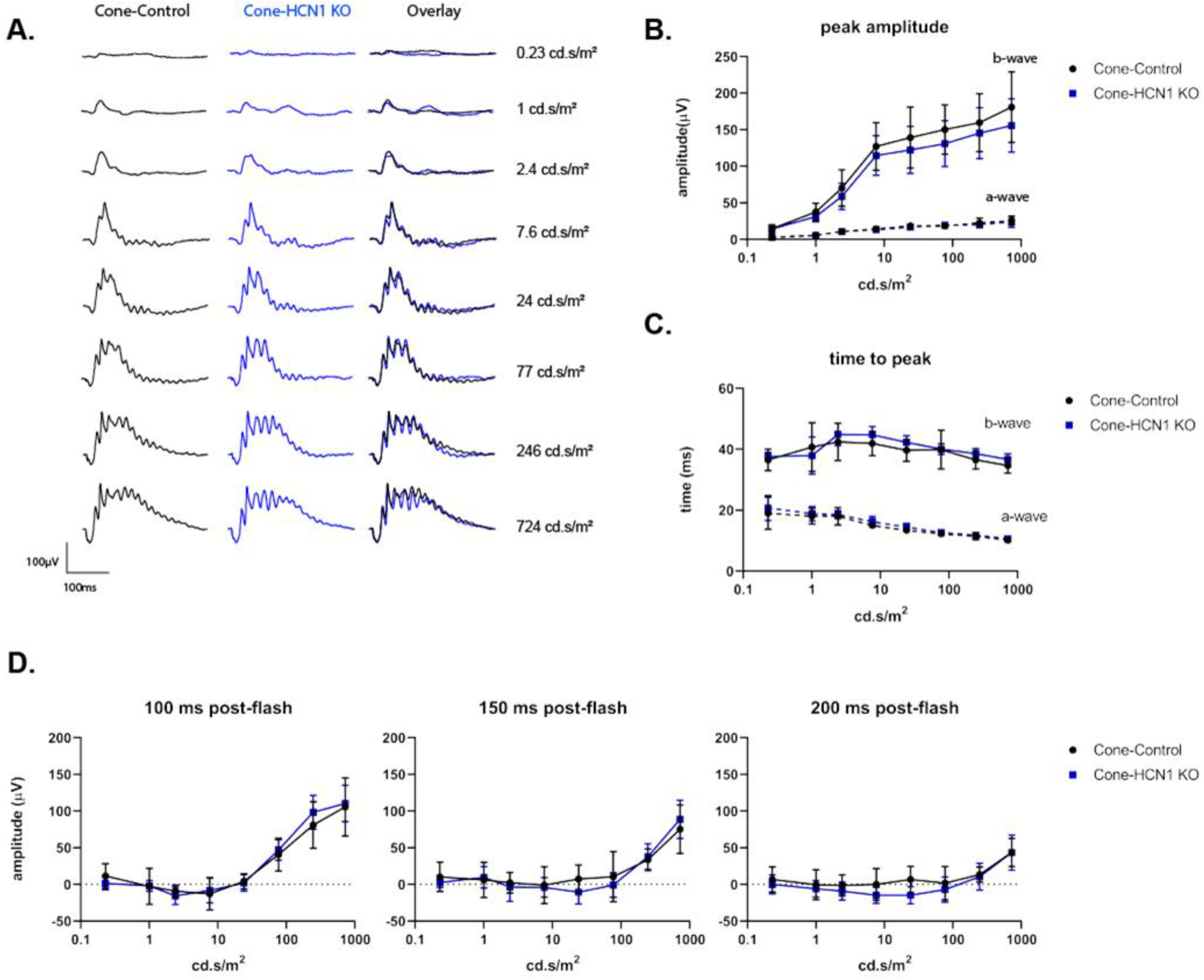
Light adapted ERG: Cone-HCN1 KO. **A)** Representative family of ERG traces from light (30 cd/m^2^) adapted Cone-Control (black) and Cone-HCN1 KO (blue) mice following a flash at the given intensity. **B)** Amplitude of a-wave (dashed line) and b-wave (solid line) plotted against stimulus intensity. **C)** Time to peak of a-wave (dashed line) and b-wave (solid line) plotted against stimulus intensity. **D)** Amplitude of the b-wave relative to baseline at 100, 150 and 200 ms after the flash. Data is presented as mean ± SD. For sample size and detailed statistics see Supplemental Tables 9.1-9.7.

We next tested both flash and sinusoidal flicker ERG protocols. The responses of the Cone-HCN1 KO mice to a 3 cd.s/m^2^ flash flicker at frequencies from 0.5 to 30 Hz were indistinguishable from Cone-Controls (Fig. 10A, B; ANOVA p = 0.2971). We tested the mice using a 30 Hz flicker with the brightest intensity our apparatus could deliver (3 to 724 cd.s/m^2^ (Fig. S5A) and again observed no difference between Cone-HCN1 KO and Cone-Controls (Fig. S5B; ANOVA p = 0.5766). Based on the studies using single cone recordings (Barrow and Wu, 2009), we expected Cone-HCN1 KO mice to have increased amplitude responses to low frequency sinusoidal flicker stimulation. However, Cone-HCN1 KO responded identically to Cone-Controls to all frequencies when a sinusoidally modulated flicker was used under photopic conditions, 30 cd/m^2^ (Fig. 10C,D; ANOVA p = 0.1892). There was similarly no difference when we analyzed the amplitude of the fundamental component (Fig. S6C ANOVA p = 0.0911). Taken together, the loss of HCN1 in cones does not appear to drive any component of the ERG phenotype associated with the global HCN1 KO. Further, this suggests that HCN1 does not accelerate the cone voltage response to the extent that we are able to detect any changes in the cone-dominated ERG response.

**Figure 10:**
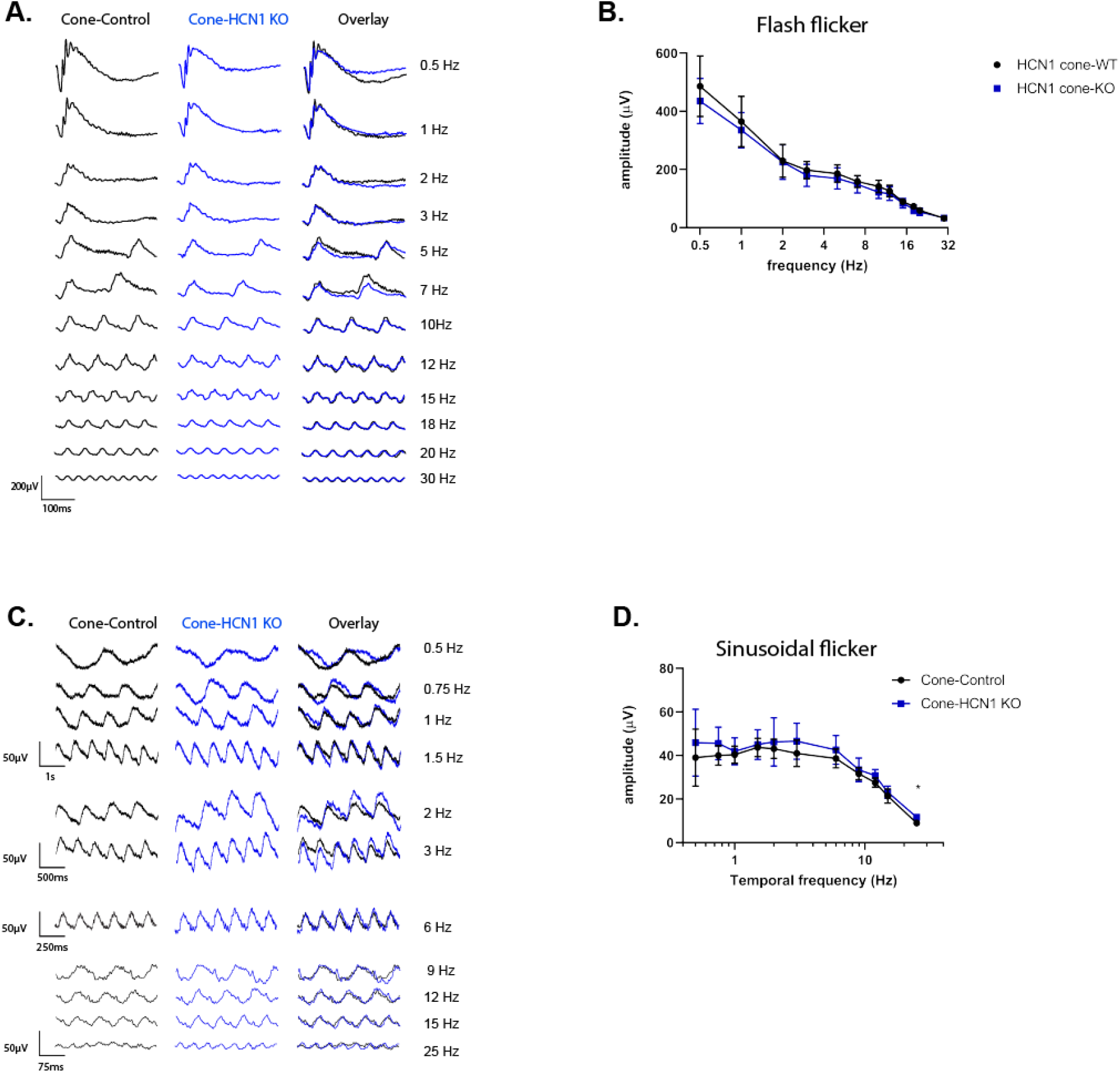
Flicker ERG: Cone-HCN1 KO. **A)** Representative family of ERG traces and **B)** response amplitude at each frequency for dark adapted Cone-Control (black) and Cone-HCN1 KO (blue) mice using a 3 cd.s/m^2^ flash flicker at the given frequency. **C)** Representative family of ERG traces and **D)** response amplitude at each frequency using a sinusoidal flicker at the given frequency with 100% contrast and a background of 30 cd/m^2^. Data is presented as mean ± SD. For sample size and detailed statistics see Supplemental Tables 10.1-10.2.

#### Cone-HCN1 KO mice have unaltered cone ON and OFF responses

To further explore the functional role of HCN1 in cones, we examined whether HCN1 is required for cone adaptation to a prolonged pulse of light. Animals were initially adapted to a 30 cd/m^2^ rod saturating background light then stimulated with one-second pulses of light ranging from 57.5 cd/m^2^ to 25,000 cd/m^2^ against this background (Fig. 11A). The initial phase of the response was unchanged as a-wave and b-wave amplitudes (Fig. 11B; ANOVA a-wave = 0.3296; b-wave p = 0.3001) and time to peak (Fig. 11C; ANOVA a-wave p = 0.0475) were not different than Cone-Controls. Note, the time to peak for the b-wave indicated a difference (ANOVA, p = 0.0365) but multiple comparisons of the individual flash intensities did not reveal a significant difference. This is consistent with the response to a brief flash. To quantify the rate of adaptation, we measured the amplitude of the decaying b-wave at 150, 250, and 500 ms after light onset and again found no significant difference in the amplitude any of these time-points (Fig. 11D; ANOVA p = 0.5624, 0.1073, 0.1336). We also examined the negative OFF response (change in voltage measured in the traces at 1000 ms after the stimulus to the trough of the sharp negative inflection). Neither the magnitude (Fig. 11E; Mixed-effects p = 0.4009) nor the timing (Fig. 11F; Mixed-effects p = 0.8846) of the OFF response was altered in the Cone-HCN1 KO mice. Thus, we see no alteration in the response to a prolonged pulse of light in the Cone-HCN1 KO mice.

**Figure 11:**
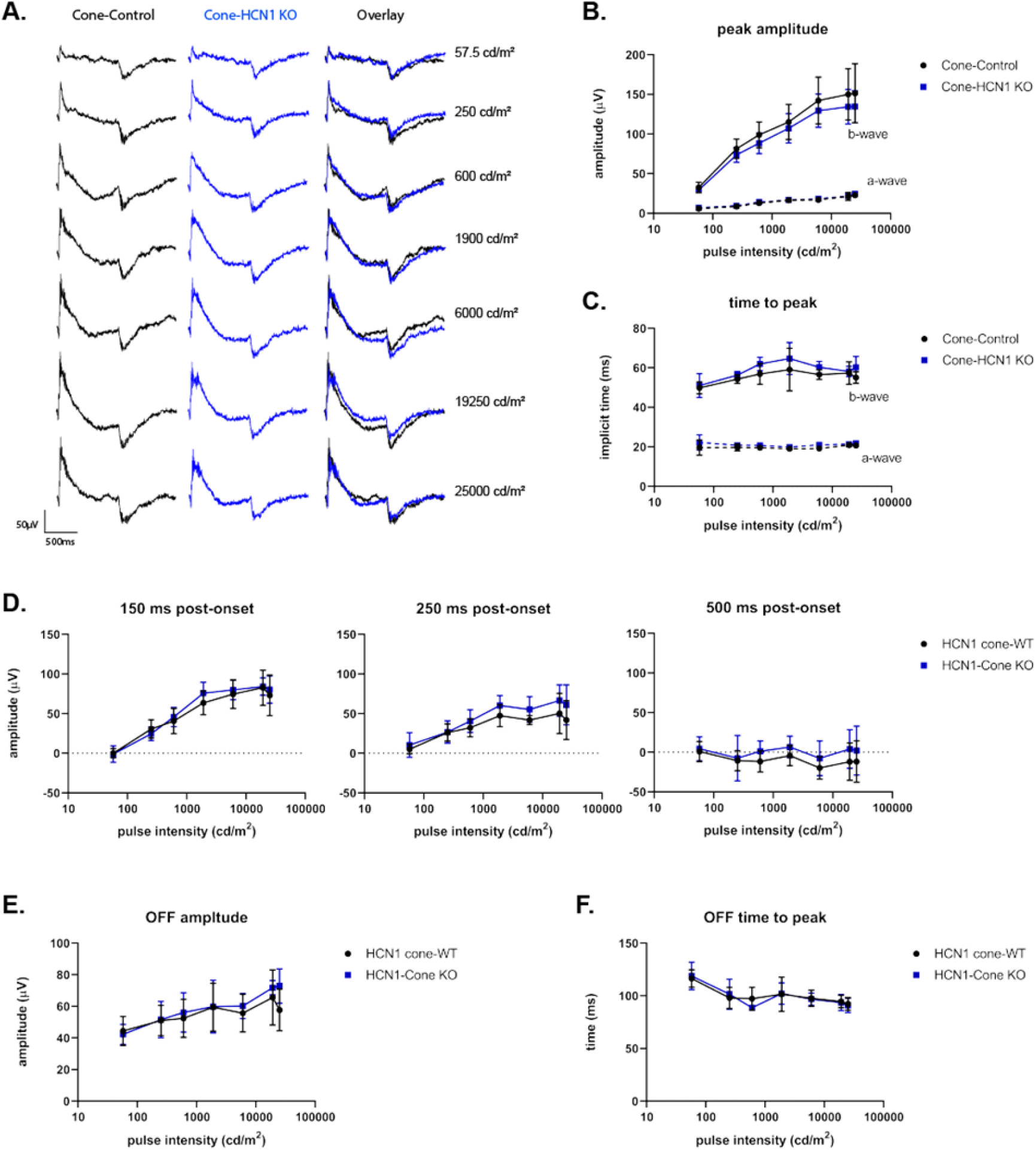
Photopic long pulse ERG: Cone-HCN1 KO. **A)** Representative family of ERG traces from light (30cd/m^2^) adapted Cone-Control (black) and Cone-HCN1 KO (blue) mice following a 1 sec pulse at the given intensity. **B)** Amplitude of the a-wave (dashed line) and b-wave (solid line) plotted against stimulus intensity. **C)** Time to peak of the a-wave (dashed line) and b-wave (solid line) plotted against stimulus intensity. **D)** Amplitude of the b-wave relative to baseline at 150, 250 and 500 ms after light onset. **E)** Magnitude of the negative inflection at light offset. **F)** Time to peak for the negative inflection at light offset. Data is presented as mean ± SD. For sample size and detailed statistics see Supplemental Tables 11.1-11.9.

#### Isolated cone responses in Cone-HCN1 KO are similar to Cone-Control Mice

We next examined directly whether cone function was altered in the Cone-HCN1-KO mice. For this, we used *ex vivo* transretinal recordings, where pharmacological manipulation in conjunction with a double flash stimulus allowed us to first dissect out the photoreceptor component from the whole retina response and then suppress the rod response to record isolated cone responses. Consistent with the in vivo ERG findings, we did not observe any significant differences in the cone response amplitude between Cone-Control and Cone-HCN1 KO mice (Fig. 12A-C). The intensity to produce half maximum response, which is a measure of sensitivity, and the dark-adapted fractional sensitivity of cone responses were also not found to be significantly different between Cone-Control and Cone-HCN1 KO mice (Table 1). This was also evident in the normalized intensity response curves which were comparable for the Cone-HCN1 KO and Cone-Control retinas under both dim light (Fig. 12D) and the brightest stimulus (Fig. 12E). Interestingly, the kinetics of the dim flash response were slightly slower in the Cone-HCN1 KO as compared to controls (Fig. 12D) though the differences were only significant for the time to peak of the response (Table 1). We also saw differences in the kinetics of the bright flash response; however, these too turned out to be statistically insignificant (Fig. 12F). Thus, there were no notable changes in cone function between Cone-HCN1 KO and Cone-Control mice.

**Table 1:** Isolated cone response parameters measured by transretinal ERG.

| | $R_{max}$ ($\mu\text{V}$) | $I_{1/2}$ (Phot/$\mu\text{m}^2$) | $S_{fD}$ (Phot$^{-1}/\mu\text{m}^{-2}$) | $t_p$ (ms) | $t_{int}$ (ms) | N |
| --- | --- | --- | --- | --- | --- | --- |
| Cone-Control | $34 \pm 5$ | $10,000 \pm 2820$ | $1.6 \pm 0.2$ | $45 \pm 1$ | $81 \pm 4$ | 7 retinas<br>(4 mice) |
| Cone-HCN1 KO | $44 \pm 8$ | $6,660 \pm 1549$ | $1.7 \pm 0.2$ | $52 \pm 1$ | $87 \pm 3$ | 13 retinas<br>(7 mice) |
| *P* value | 0.29 | 0.32 | 0.68 | 0.0035 | 0.19 |  |

$R_{max}$ , saturated response amplitude measured at the plateau; $I_{1/2}$ , intensity required to produce half of the saturated response; $S_{fD}$ , fractional dark-adapted sensitivity; $t_p$ , time to peak of a dim flash response; $t_{int}$ , integration time of the response.

**Figure 12:**
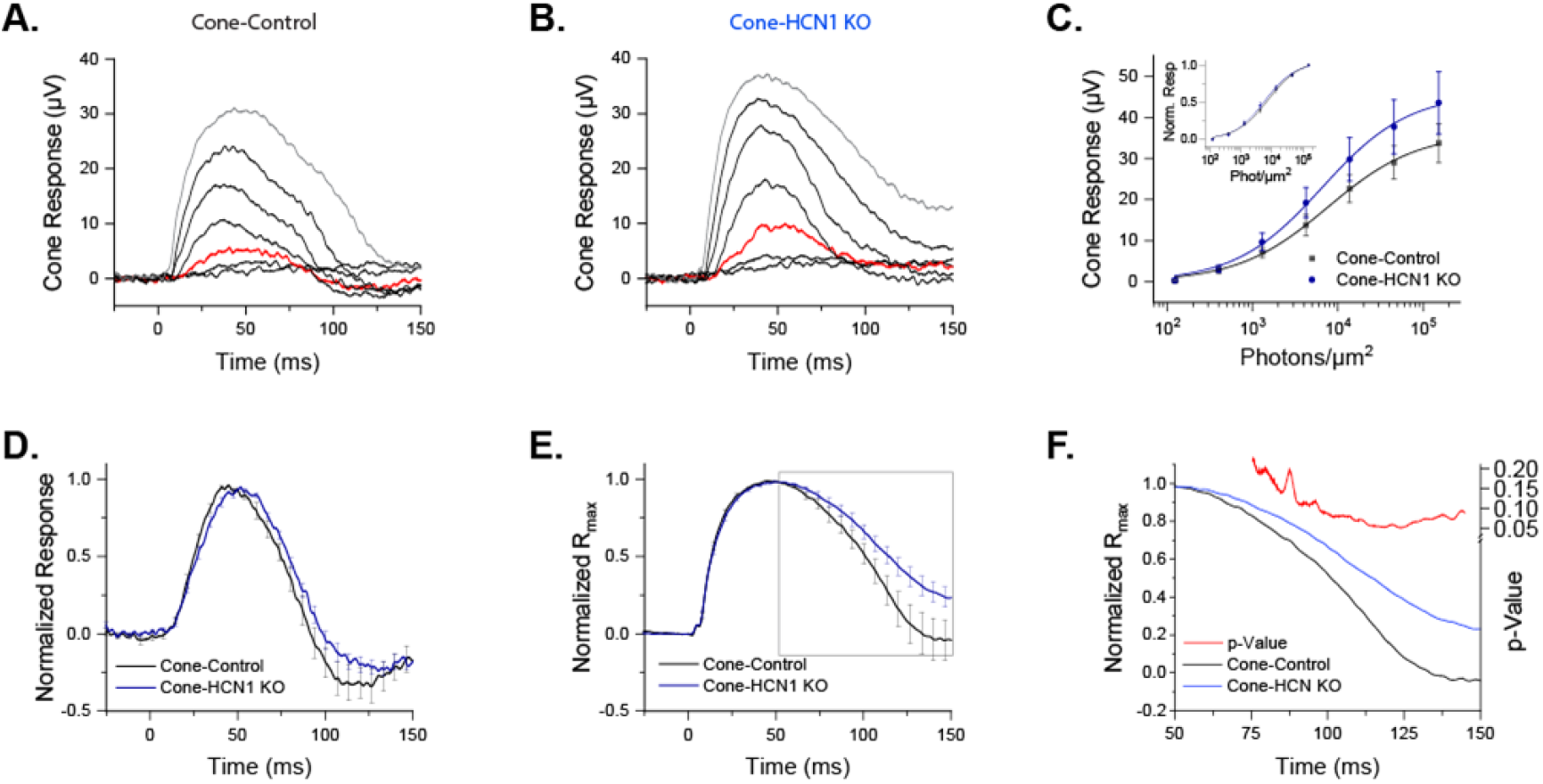
Isolated cone responses: Cone-HCN1 KO. Representative responses to a family of flashes (123, 401, 1295, 4223, 13717, 45465 and 151808 photons μm^−2^ presented at T_0_ from individual Cone-Control (**A**) and Cone-HCN1 KO retinas obtained by transretinal ERG recordings. (**B**). The red traces in the two panels show responses to the same test flash intensity (1295 photons μm^−2^), highlighted here for comparison. **C)** Ensemble-averaged intensity-response curves for cones from Cone-Control (black) and Cone-HCN1 KO (blue) fit to a Naka-Rushton function; **inset** shows the corresponding normalized intensity response curves. **D)** Ensemble-averaged normalized dim flash responses (red trace in A and B). **E)** Ensemble-averaged normalized bright flash responses (grey trace in A and B). **F)** Enlarged segment of boxed portion of E) from 50 to 150 ms. Red trace plotted on the right y-axis represents p-value for amplitude difference at each timepoint; note p > 0.05 for all timepoints. Data is presented as mean ± SEM.

#### Behavioral responses are unaltered in the Cone-HCN1 KO mice

Finally, we tested the behavioral responses of Cone-HCN1 KO animals using the optomotor response. Temporal contrast sensitivity of Cone-HCN1 KO and Cone-Controls was measured using a background of 70 cd/m^2^ with a 0.383 c/d spatial frequency at temporal frequencies of 1.5, 3, and 6 Hz. The higher spatial and temporal frequencies in the stimulus were selected to favor cone-driven responses (Umino et al., 2008). Consistent the prior experiments, we found that the Cone-HCN1 KO mice behaved the same as Cone-Controls (Fig. S7; p = 0.427). Altogether, this data demonstrates that selective ablation of HCN1 from cones does not impact visual function.

## DISCUSSION

Photoreceptor HCN1 channels carry a current that acts to reset photovoltage after light triggered hyperpolarization. Prior studies of HCN1 have led to several open-ended questions and some contradictory findings. Studies of HCN1 whole-body knock out mice are complicated by the expression of HCN1 in multiple neurons within the retina, and numerous other regions of the CNS. Here, we generated mouse lines with HCN1 knocked out selectively from rods or cones and characterized retina function using ERG and OMR. This revealed several novel findings. In dim light where vision is dependent on rods, loss of HCN1 enhances the rod response. Under high mesopic and into photopic conditions, loss of HCN1 in rods prevents transmission of cone-driven signals. This illustrates the importance of voltage modulation as an adaptation mechanism to control rod output. On the other hand, loss of HCN1 selectively from cones does not change retinal responses.

### The role of HCN1 in rods under scotopic conditions

HCN1 has a well characterized role accelerating the rod voltage recovery and as such is well positioned to extend the temporal resolution of rods (Barrow and Wu, 2009; Fain et al., 1978; Knop et al., 2008; Sothilingam et al., 2016). However, there are conflicting reports regarding the role HCN1 plays in shaping the response to high temporal frequency stimulation (Barrow and Wu, 2009; Della Santina et al., 2012; Gargini et al., 1999; Knop et al., 2008; Seeliger et al., 2011; Sothilingam et al., 2016). Using a dim, sinusoidal flicker, we observed that rod specific knockout of HCN1 does not appear to dampen the ERG response to a high frequency flicker. Instead, we see a trend toward elevated response amplitudes to low frequencies in the Rod-HCN1 KO. This is consistent with direct rod recordings demonstrating the voltage response to a flicker is larger when *I*_h_ is blocked and suggests that under dim conditions, HCN1 acts to limit the amplitude of the rod hyperpolarization response under low frequencies (Barrow and Wu, 2009). Also, a previous flicker ERG study showed that global HCN1 KO mice exhibited a slightly left shifted frequency response curve (Della Santina et al., 2012). We do not see this in our data and the difference between the global HCN1 KO and the rod specific HCN1 KO may reflect the function of HCN1 in bipolar cells which is poorly understood and merits further investigation (Cangiano et al., 2007; Della Santina et al., 2012).

### The role of HCN1 in rods under mesopic conditions

Under brighter conditions, where rods would be driven to more hyperpolarized potentials, we would imagine that HCN1 would exert a greater influence on rod function and would be required to accelerate recovery following initial light stimulation and thus be necessary to form the response to high frequency flicker. Instead, we found that the rod-specific HCN1 KO had altered responses to low but not high-frequency modulation. The ERG response to a low frequency mesopic sinusoidal flicker is biphasic with an initial sharp “b-wave-like” component followed by a broader “c-wave-like” component (Inamdar et al., 2021). This contrasts with the response to the same frequency stimulus under dim conditions where the response consists of a single broad peak consistent with the “b-wave-like” and “c-wave-like” components being temporally superimposed. The fact that the b-wave-like and c-wave-like components appear superimposed under mesopic conditions in the Rod-HCN1 KO suggests that classical adaptation of the phototransduction cascade is not sufficient to accelerate the kinetics of the rod-driven response. Instead, this acceleration depends on HCN1 mediated voltage adaptation.

The prevailing model for the role of HCN1 in integration of rod and cone driven signaling is that under mesopic conditions HCN1 acts to accelerate rod voltage responses so that rod output does not saturate the downstream circuitry needed to transmit cone-driven signals (Seeliger et al., 2011). When using a flash flicker, there was prolonged rod-driven activity and suppression of cone responses in Rod-HCN1 KO mice similar to the global HCN1 KO. However, the flash flicker ERG protocol results in an increase in mean illumination as the frequency increases so to separate these components. To address this, we used a sinusoidal flicker that allowed us to hold the mean illumination constant while varying temporal frequency. This revealed that the rod-mediated saturation of downstream circuitry occurs under bright conditions (high mesopic and photopic) even beyond the point of rod saturation and is not dependent on delayed rod voltage recovery. Thus, we do not disagree with the model proposed by Seeliger and colleagues, but note it is incomplete as it does not account for differing lighting environments.

### How does HCN1 mediated modulation of rod function affect vision?

Taken together, our data suggest that HCN1 is an essential component of an adaptation mechanism that acts at the level of rod voltage. This voltage adaptation accelerates rod response kinetics as light levels increase through the scotopic and low mesopic range where rod-signaling dominates the retinal circuit and limits rod hyperpolarization, effectively limiting the output of rods onto the retinal circuit, under elevated lighting conditions where cone-derived signaling dominates. This complements classical phototransduction adaptation which accelerates rod kinetics while suppressing sensitivity to the point of saturation. Interestingly, we see that the requirement for HCN1 extends beyond the point of rod saturation as rod expressed HCN1 remains essential for cone-derived signals to transmit across the retinal circuit under photopic conditions.

While our ERG tests demonstrate a clear role for rod expressed HCN1 in shaping retinal signaling, it is unclear how this would impact visual function. Using the OMR, we observed a relatively minor impact on vision as Rod-HCN1 KO mice had largely normal OMR responses with only low frequency stimulation under mesopic conditions eliciting a detectable reduction in contrast sensitivity. Given the severity of rod saturation under high mesopic and photopic conditions, it will be interesting to test the Rod-HCN1 KO mice with more rigorous psychophysical assays, such as the forced-choice operant assay (Umino et al., 2018). Human HCN1 variants do not report visual dysfunction, however these patients exhibit a range of neurological problems which may preclude reporting of visual impairments (Bonzanni et al., 2018; Marini et al., 2018). This is particularly true if foveal cones, which are devoid of coupled rods, are not impacted by loss of HCN1 as this study would suggest.

### The role of HCN1 in cones

Previously, little was known regarding the role of HCN1 in shaping cone-driven, photopic responses. *I*_h_ can be recorded from cones and while there is some question regarding potential expression of HCN2 and HCN3, the data suggests that cone *I*_h_ is mediated by HCN1 channels (Barrow and Wu, 2009; Della Santina et al., 2012; Fain and Sampath, 2021; Ingram et al., 2020; Müller et al., 2003; Voigt et al., 2019). The function of *I*_h_ in cones has not been studied in a mammalian system but in salamander, *I*_h_ block results in a significant overshoot in light induced hyperpolarization with minor impact on voltage recovery rate (Barrow and Wu, 2009). Using a variety of ERG tests, we did not observe any impact on the cone-driven responses when HCN1 was knocked out in cones. The isolated cone a-wave measured by transretinal ERG was also not significantly altered, though, we did see a small but statistically significant delay in the HCN1 KO cone response kinetics which is consistent with earlier findings in amphibian cones. Thus, HCN1 may have a subtle role in shaping the cone response that was not apparent in the assays performed here. Cone-expressed HCN1 may have additional functions not examined in this study. For instance, experiments from goldfish suggest HCN1 also contributes to contrast-activated adaptation of cones (Howlett et al., 2017). HCN1 is evolutionarily conserved, and it seems unlikely that this channel would be fully dispensable for cone function and more precise and direct tests than the full field ERG used in this study may be needed to parse out the function of this channel in cones.

In conclusion, we see that HCN1 is unexpectedly dispensable for intrinsic cone function and is not required to drive retinal response to a high frequency flickering light. Instead, HCN1 primarily mediates rod voltage adaptation that suppresses rod output onto the retinal circuit as light intensity increases. This allows cone-derived signals to dominate the circuit. Thus, HCN1 channels appears to limit rod excitability, consistent with their well-established role limiting neuronal excitability in the central brain, despite rods operating under an inverted voltage paradigm compared to traditional neurons.

## MATERIALS AND METHODS

### Animals

HCN1 floxed mice (129S/SvEv-Hcn1tm1Kndl/J, JAX: 028300) were obtained from The Jackson Laboratory. Rod specific Cre mice, Rho-iCre, mice (B6.Cg-Pde6b+ Tg(Rho-icre)1Ck/Boc, JAX: 015850) were obtained from JAX Labs and cone specific Cre mice, HRGP-Cre, mice (STOCK Tg(OPN1LW-cre)4Yzl/J (JAX: 032911)) were a generous gift from Yun Zheng Le, University of Oklahoma. The HCN1 rod-KO and HCN1 cone-KO lines used in this study were generated by crossing the rod and cone specific Cre driver lines onto the HCN1 floxed background. Both Cre and iCre transgenes were maintained as hemizygous in these lines and Cre negative litter mates were used as controls. Animals were genotyped using published protocols or through the services of Transnetyx (Cordova, TN). Mice were housed in a central vivarium, maintained on a standard 12/12-hour light/dark cycle, with food and water provided ad libitum in accordance with the Guide for the Care and Use of Laboratory Animals of the National Institutes of Health. All procedures adhered to the ARVO Statement for the Use of Animals in Ophthalmic and Vision Research and were approved by the University of Iowa IACUC committee. For all experiments, both male and female mice were used.

### Immunohistochemistry

Animals were euthanized by CO_2_ asphyxiation followed by cervical dislocation. Immediately after death, the eyes were enucleated, dissected into eye cups, and fixed for 1 hour in 4% paraformaldehyde in PBS (137 mM NaCl, 2.7 mM KCl, 10 mM Na_2_HPO_4_, 1.8 mM KH_2_PO_4_, pH 7.4). Fixed retinas were incubated overnight in a 30% sucrose solution prior to freezing in O.C.T. compound. Immunostaining was performed as previously described (Inamdar et al., 2018).

Microscopes used were either a THUNDER Imager 3D Tissue Fully automated upright research microscope Leica DM6 B equipped with a Leica DFC9000 GT camera, or a Zeiss 710 scanning laser confocal microscope, each equipped with 1.4 NA 40X objective. Image analysis including THUNDER computational clearing was performed using the LASx software.

Primary antibodies used: Mouse anti-HCN1 (Neuromab Cat# N70/28, RRID:AB_2877279) used at 1:500; guinea pig anti-Cre (Synaptic Systems Cat# 257 004, RRID:AB_2782969) used at 1:750, rabbit anti-cone arrestin (Millipore Cat# AB15282, RRID:AB_1163387) used at 1:750.

Secondary antibodies used: goat anti-Guinea pig Alexa488 (Molecular Probes Cat# A-11073, RRID:AB_2534117), used at 1:500; goat anti-mouse Alexa488 (Thermo Fisher Scientific Cat# A-11001, RRID:AB_2534069), and goat anti-rabbit Alexa647 (Thermo Fisher Scientific Cat# A-21245, RRID:AB_2535813) all used at 1:500.

Line scan analysis was performed using FIJI (imageJ). In order to compare cone arrestin and HCN1 signal intensity simultaneously, the signal intensity was normalized to the maximum signal such that the maximum value was 1 for both cone arrestin and HCN1 signal. For examination of the correlation between HCN1 and cone arrestin, long line scans (~100um) were drawn across the inner segment and the normalized HCN1 intensity was plotted against the normalized cone arrestin signal. The was calculated using GraphPad Prism (ver 8). This analysis was performed on single optical sections across four retinas for both Cone-Control and Cone-HCN1 KO. The average r value was compared using a T-test.

### In vivo Electroretinography (ERG)

ERG recordings were obtained using an Espion V6 Diagnosys Celeris system (Diagnosys LLC, Massachusetts) with individual eye integrated light stimulators and voltage recording electrodes. Animals were between 7 and 10 weeks of age at the time of analysis. Between 3-11 animals of each genotype were used per test as detailed in the Supplemental Tables. Animals were dark adapted overnight (~18 hours) prior to recordings and all steps performed on the day of recording were under dim red light. Animals were anesthetized just prior to analysis by a mixture of ketamine (87.5 mg/kg) and xylazine (2.5 mg/kg). Tropicamide (1%) was used to dilate the pupils and Genteal gel (0.3% Hypromellose) was used to keep the eyes hydrated. Body temperature was maintained during testing by an internal heating system in the ERG machine or heating pads and by heating pads during animal recovery. The single flash ERG was performed with a reference electrode placed subcutaneously along the nasal ridge or in the mouth and a grounding electrode inserted into the haunch. The flash flicker, photopic sinusoidal flicker, and long pulse ERG tests were performed without using a ground electrode, instead using the non-stimulated eye as a reference.

The scotopic single flash ERG series consisted of 8 steps using a white flash ranging in intensity from 0.003 to 100 cd.s/m^2^. For intensities from 0.003-0.1 cd.s/m^2^ responses were averaged from ten sweeps collected at intervals of 5s, for 1 and 3 cd.s/m^2^ fifteen sweeps were collected at intervals of 10s, for 10 cd.s/m^2^ five sweeps were collected at intervals of 10s, for 30 cd.s/m^2^ three sweeps were collected at an interval of 10 s, and after 30 s a single recording was collected at the highest intensity (100 cd.s/m^2^).

The photopic ERG series consisted of a 10-minute adaptation step to the 30 cd/m^2^ white light background followed by eight steps using a flashing white light ranging from intensity of 0.23-724 cd.s/m^2^ above this background. Responses were averaged from ten sweeps with an inter-sweep interval of 5 s.

The flash flicker series consisted of 12 steps using 3 cd.s/m^2^ flickering white light with a frequency ranging from 0.5-30 Hz, for frequencies from 0.5-5 Hz response 3 sweeps were collected, for 7 Hz five sweeps were collected, for 10-15 Hz twenty sweeps were collected, and for 18-30 Hz fifty sweeps were collected. There was no delay between sweeps and the first sweep was always rejected.

The 30 Hz flicker series consisted of 6 steps using a 30 Hz flickering white light with intensity stepping from 3 cd.s/m^2^ to 724 cd.s/m^2^. For each step, 50 sweeps were collected, there was no delay between sweeps, and the first sweep was always rejected.

The long pulse protocol consisted of a 10-minute adaptation step to the 30 cd/m^2^ white light background followed by seven steps using a 1 second white light ranging from 57.5 cd/m^2^ to 25,000 cd/m^2^ followed by 1 second at the 30 cd/m^2^ background. Responses were averaged from ten sweeps with a 3.5 second inter-sweep interval.

The photopic sinusoidal flicker ERG performed on Cone-HCN1 KO and Cone-Control mice consisted of a 10-minute adaptation step to 30 cd/m^2^ white light. For stimulation, light intensity was modulated at 100% contrast with a mean illumination of 30 cd/m^2^. Recording time was limited to 4000 ms for 0.5 to 1.5 Hz, 2000 ms for 2 to 3 Hz, 1000 ms for 6 Hz, and 300 ms for 9 to 25 Hz. Responses were averaged from 20 sweeps for frequencies from 0.5 to 12 Hz and 50 sweeps for frequencies from 15 to 25 Hz.

Sinusoidal flicker ERGs performed on Rod-HCN1 KO and Rod-Control mice were recorded using the Espion E2 system and a Ganzfeld ColorDome stimulator (Diagnosys, Espion E^2^ system) as previously described (Umino et al., 2019). Following anesthesia with a mixture of ketamine/xylazine mixture at 120 and 10mg/kg, respectively, mice were placed on a heating pad (37°C) and reference and ground electrodes were placed in the mouth and intradermally next to the tail, respectively. A drop of 2.5% hypromellose GONAK solution (AKORN) was applied to the eye and custom-made conductive silver thread electrode and contact lens were placed on the cornea under infrared illumination. After completing the setup procedure mice were light-adapted for 1 hour prior to the start of the recordings. Booster shots with 30% of the original dosage of ketamine/xylazine was administered every 40 min to maintain the mice anesthetized over prolonged time required to complete a recording session. Flicker ERGs were elicited by a sinusoidally modulated monochromatic light stimulus (530 nm) at various levels of mean luminance (0.05, 1, 10 and 30 cd/m^2^) and temporal frequencies (from 0.5 Hz to 30 Hz). Contrast was set at 100%. Conversion from luminance to rate of rhodopsin excitation was performed as previously described (Umino et al., 2019). Note that rod effective collecting area values are likely to change following prolonged bleaching at the high irradiance levels used in our ERG experiments therefore the R*/rod/s values indicate initial photoisomerization rates at onset of background.

Traces were analyzed using the Espion software (Ver 6.58.17). In the rod-driven and cone-driven series, a-waves were identified as the negative peak following stimulus. A-wave amplitude was measured from baseline to the trough of the a-wave. B-waves were identified as the second major positive peak of the ascending positive inflection following the a-wave. B-wave amplitude was measured from the trough of the a-wave to the peak of the b-wave. For the flicker series (both flash and sinusoidal), the amplitude of the response was measured as the amplitude from trough to peak. For the sinusoidal flicker the fast Fourier transform was performed using MATLAB (R2021) and the amplitude of the fundamental component was assessed. For the long pulse protocol, the a-wave and b-wave were measured as before while the OFF-response was measured as the change in voltage from the start of light offset at 1000 ms to the trough of the negative peak.

### Optomotor response (OMR)

Optomotor contrast sensitivities of mice were measured with the OptoMotry© system using a two-alternative forced choice protocol as described previously (Prusky et al., 2004; Umino et al., 2008). Briefly, dark-adapted mice were placed on a pedestal at the center of the OptoMotry chamber. An observer monitored the reflex head movement of mice in response to the clockwise or counter-clockwise rotation of sinusoidal pattern gratings. The observer was blind of direction of pattern rotation. Auditory feedback indicated whether the selected direction was correct or incorrect. Trial durations were 5 sec. A computer program controlled contrast of the stimulus following a staircase paradigm (Umino et al., 2006). The threshold was set to 70% correct response. Optomotor contrast sensitivity was defined as the reciprocal of the contrast threshold value. Luminance within the Optomotry chamber was attenuated with neutral density filters (Lee filters) positioned between the computer monitors and the mice. Conversion of luminance values to rod photoisomerization rates were estimated using pupil areas values as previously (Bushnell et al., 2016). Optomotor contrast sensitivity of each mouse was determined as the average of four to five independent trials. Results from trials differing by >2 SD from the average were discarded.

### Transretinal Electroretinography (ERG)

Overnight dark-adapted animals were euthanized by CO_2_ inhalation followed by enucleation of the eyes in dim red light. The eyes were then dissected under infrared illumination in a Petri dish containing oxygenated Ames medium (Sigma) under a microscope. An incision was made close to the limbus using the tip of a scalpel, then the eyeballs were hemisected using micro-scissors and posterior eye cups were separated from the lens and cornea. The sclera and RPE were gently detached using forceps. The retinas were stored in oxygenated Ames medium in a dark chamber until recording. Recordings were made using previously described methods. The retina was mounted in a closed chamber with photoreceptors facing up. The chamber was then positioned under a microscope (Olympus BX51). During the recording, the retina was continuously supplied with heated Ames medium at 3-5 ml/min and maintained at 36-37 °C. Recordings were started after adapting the retina to these conditions for 15 to 20 minutes. Flash stimuli were generated using computer-controlled LEDs and were projected on the retina using the microscope optics. The Ames medium contained 50 μM DL-AP_4_ (Tocris) and 100 μM BaCl_2_ (Sigma) to isolate the photoreceptor response from that of the whole retina. The responses were amplified using a differential amplifier (Warner Instruments), low-pass filtered at 300 Hz (Krohn Hite Corp.), digitized using Digidata 1440 (Molecular Devices), and sampled at 10 kHz. Data was acquired and recorded using pClamp 10 software on a computer. To separate the cone response from the rod response, first a probe flash (12700 photons μm^−2^) was presented to saturate the rod response. Then after 350 ms, a second flash was presented to record the cone response as described earlier. Data are presented as mean ± SEM and student’s t-test was used to estimate the statistical significance. Dark adapted fractional sensitivity (S_fD_) was calculated by dividing the dark-adapted dim flash response, normalized to the maximum response, by the flash strength. The intensity-response data were fitted to a Naka-Rushton function using the following equation:

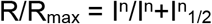

where, R_max_ is the maximum amplitude, I is the flash intensity, n is the Hill coefficient and I_1/2_ is the half saturating light intensity.

### Statistical Analysis

Statistical analysis was performed using GraphPad Prism (Ver. 8.4). For ERG and OMR analysis we compared experimental animals with littermate controls using a two-way ANOVA with a Sidak multiple comparison test to compare the genotype difference at individual flash intensities or flicker frequencies. When noted, a mixed effect analysis was used rather than the two-way ANOVA. This was limited to situations where a few data points were excluded due to technical issues. P-values reported in the text and figure legends refer to the genotype differences except where individual intensities/frequencies are compared, and the p-value presented reflects the adjusted p-value from the Sidak multiple comparison test. Full descriptive statistics including mean differences with 95% confidence intervals are available in Supplemental Tables. Sample sizes, also detailed in the supplemental tables, are individual retinas (trans-retinal ERG), individual animals (for the majority of ERG tests), or individual eyes (for the sinusoidal flicker ERG performed on Rod-HCN1 KO and Rod-Controls). Error bars shown on graphs are S.D. with the exception of the *ex* vivo transretinal ERG and OMR assays where bars represent SEM, as noted in the figure legend. Asterisks on graphs indicate adjusted p-value range as follows: p < 0.05 (*); p < 0.01 (**); p<0.001 (***); p < 0.0001 (****).

## Supporting information

Supplemental Statistical Tables

## ACKNOWLEDGEMENTS

This work was supported by the National Eye Institute (R01 EY020542 to SAB, R01s EY027387 and EY030912 to VJK, and R01 EY026216 to ES). We also acknowledge unrestricted grants from Research to Prevent Blindness to the Department of Ophthalmology and Visual Sciences at SUNY Upstate Medical University, NY and the Department of Ophthalmology at University of California, Irvine, CA.

## COMPETING INTERESTS

The authors have no competing interests to disclose.

## Notes

### Competing Interest Statement

The authors have declared no competing interest.

