## Supplemental Figures for "Cone-driven retinal responses are shaped by rod but not cone HCN1"

### SUPPLEMENTAL FIGURE LEGENDS

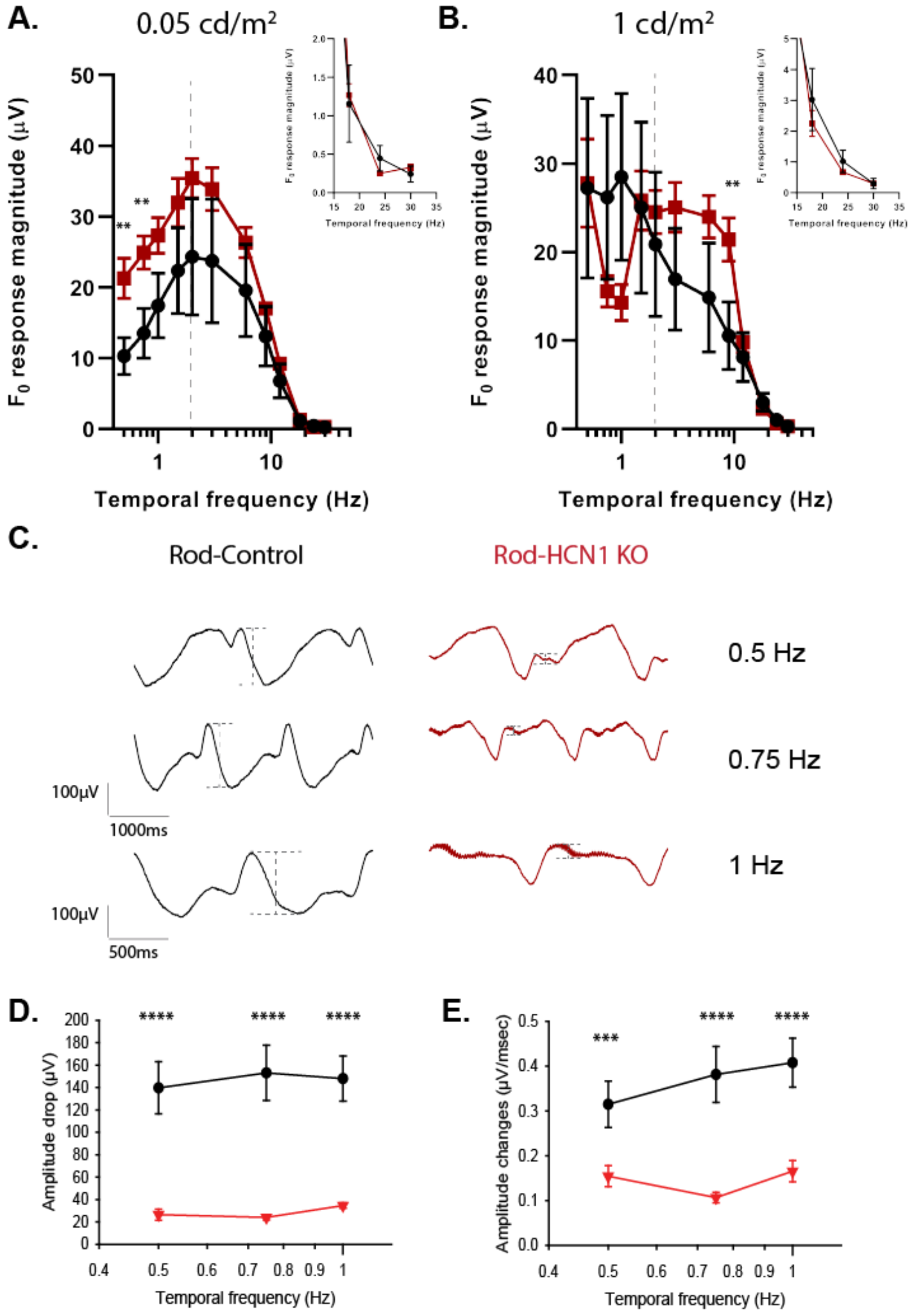


Figure S1 (related to Figure 3): Low light sinusoidal flicker ERG: Rod-HCN1 KO

**A)** Amplitude of the fundamental component ($F_0$) plotted against temporal frequency for the ERG response to a sinusoidal flicker with mean illumination of 0.05 cd/m$^2$ (data presented in Fig. 3A, B) or **B)** with mean illumination of 1 cd/m$^2$ (data presented in Fig 3C, D). Dashed line at 2 Hz below which responses were non-linear. Inset shows the data for the response to 18, 24, and 30 Hz flicker on a zoomed in scale. **C)** Low frequency responses were alternatively analyzed by measuring the drop in the b-wave-like peak as indicated by the grey dashed lines in the representative traces to a sinusoidal flicker with mean illumination of 1 cd/m$^2$ (taken from Figure 3). **D)** Magnitude of the b-wave-like voltage drop. **E)** Rate of the voltage drop from the b-wave-like peaks. Rod-Control (black), Rod-HCN1 KO (red). Data is presented as mean ± SD. For sample sizes and detailed statistics see Supplemental Tables S1.1-S1.4.

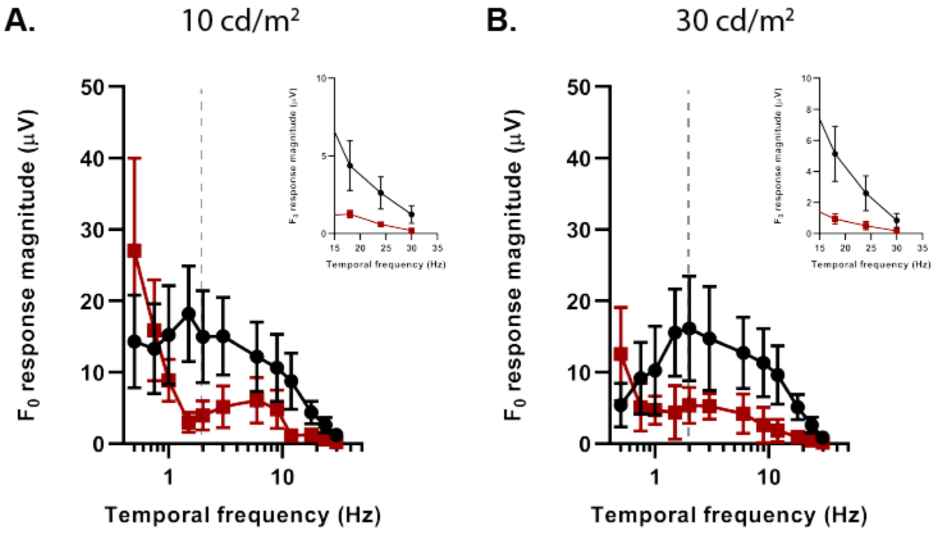


<u>Figure S2 (related to Figure 4): Bright sinusoidal flicker ERG: Rod-HCN1 KO</u>

**A)** Amplitude of the fundamental component ($F_0$) plotted against temporal frequency for the ERG response to a sinusoidal flicker with mean illumination of 10 cd/m$^2$ (data presented in Fig. 4A, B). **B)** Amplitude of the fundamental component ($F_0$) plotted against temporal frequency for the ERG response to a sinusoidal flicker with mean illumination of 30 cd/m$^2$ (data presented in Fig 4C, D). Dashed line at 2 Hz below which responses were non-linear. Inset shows the data for the response to 18, 24, and 30 Hz flicker on a zoomed in scale. Data is presented as mean ± SD. For sample sizes and detailed statistics see Supplemental Tables S2.1-S2.2.

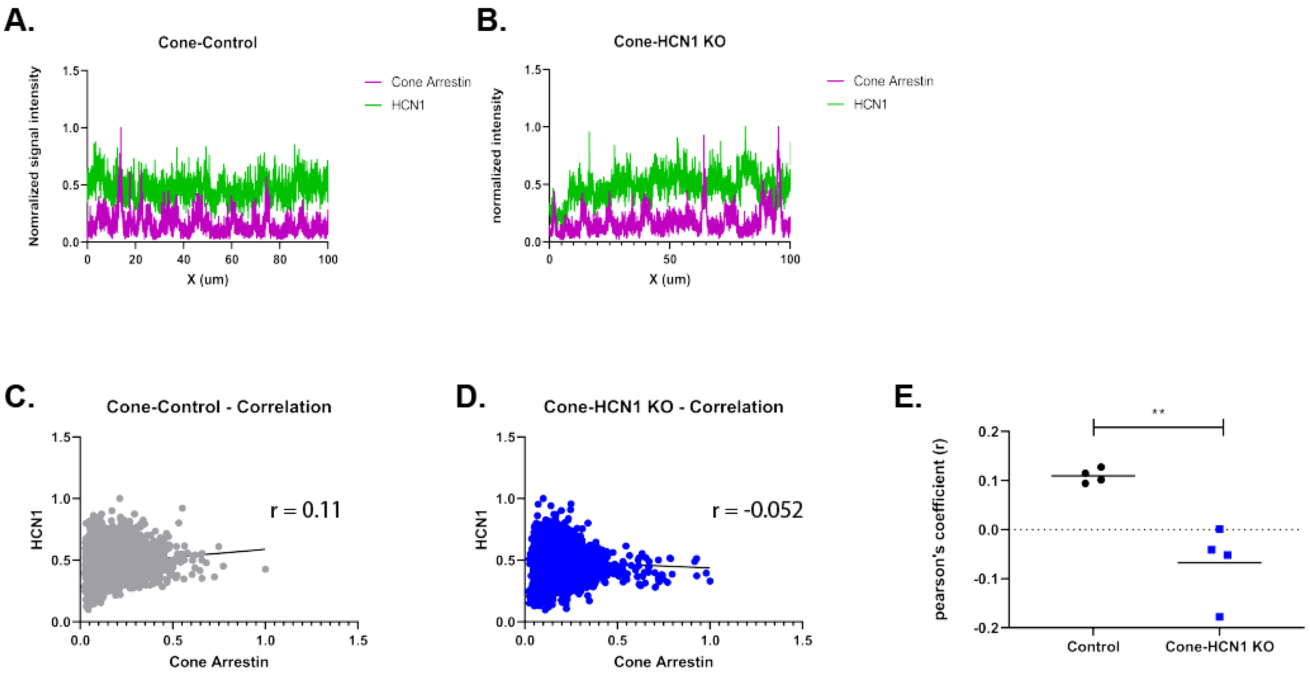


Figure S3 (related to Figure 8): Validation of Cone-HCN1 KO line

Representative profiles of HCN1 and cone arresting signal intensity across a 100 μm line through the inner segment of Cone-Control **(A)** and Cone-HCN1 KO **(B)** retina. **C, D)** Normalized HCN1 signal intensity plotted against normalized cone arrestin intensity from plots (A) and (B) with the Pearson's correlation coefficient (r). **E)** Pearson's correlation coefficient calculated from long line scans as in C and D from 4 individual retina for each genotype with data presented as Data is presented mean ± SD. T-test p = 0.0040. For sample sizes and detailed statistics see Supplemental Tables S3.

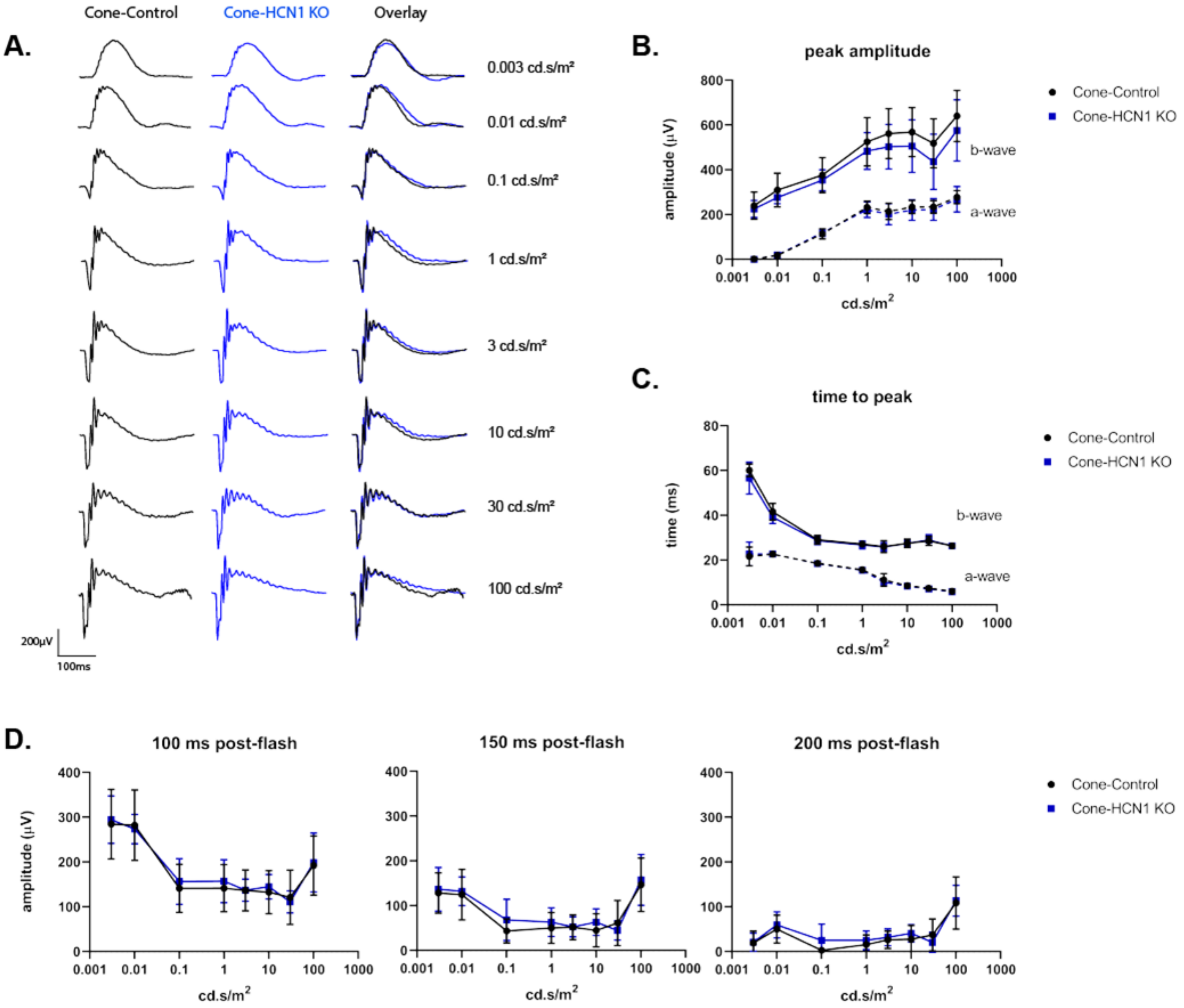


<u>Figure S4 – Dark adapted ERG: Cone-HCN1 KO</u>

**A)** Representative family of ERG traces from dark adapted Cone-Control (black) and Cone-HCN1 KO (blue) mice following a flash at the given intensity. **B)** Amplitude of a-wave (dashed line) and b-wave (solid line) plotted against stimulus intensity. **C)** Time to peak of a-wave (dashed line) and b-wave (solid line) plotted against stimulus intensity. **D)** Amplitude of the b-wave relative to baseline at 100, 150, or 200 ms after the flash. Data is presented as mean ± SD For sample size and detailed statistics see Supplemental Tables S4.1-S4.7.

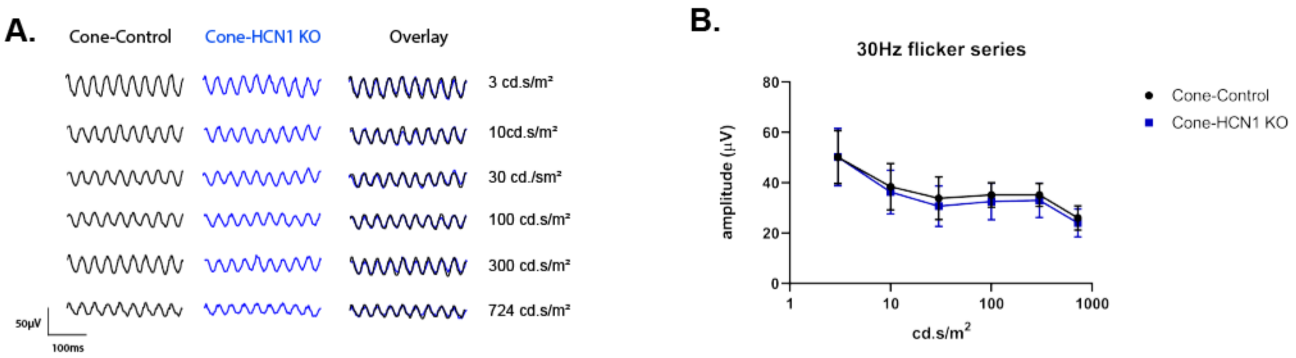


Figure S5: Bright Flicker ERG: Cone-HCN1 KO

**A)** Representative family of ERG traces from dark adapted Cone-Control (black) and Cone-HCN1 KO (blue) mice stimulated with a 30 Hz flicker at the given light intensity. **B)** Response amplitude plotted against intensity. Data is presented as mean ± SD. For sample size and detailed statistics see Supplemental Table S5.

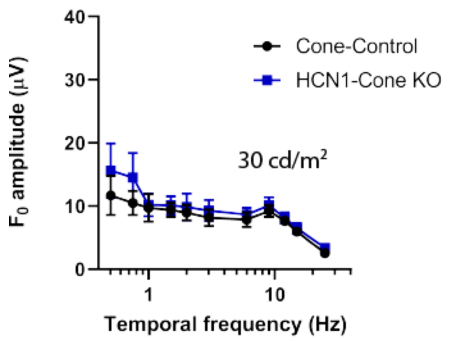


Figure S6 (related to Figure 10): Flicker ERG: Cone-HCN1 KO

**C)** Amplitude of the fundamental component ($F_0$) plotted against temporal frequency for the ERG response to a sinusoidal flicker with mean illumination of 30 cd/$m^2$ (data presented in Fig 10C, D). Data is presented as mean ± SD. For sample size and detailed statistics see Supplemental Table S6.

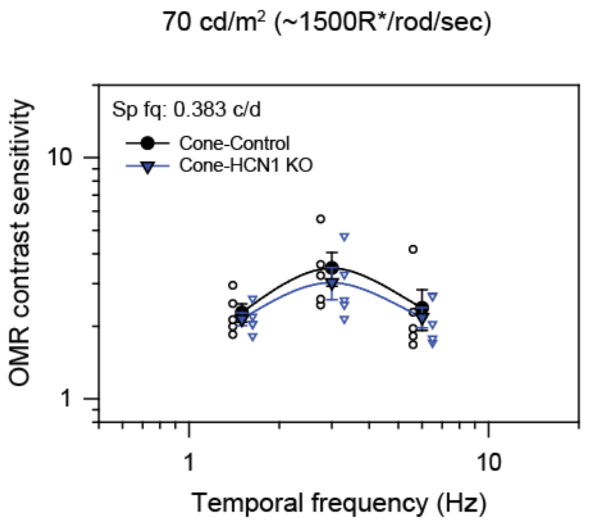


Figure S7: Optomotor temporal contrast sensitivity: Rod-HCN1 KO

**A)** Contrast sensitivity of Cone-Control (black) and Cone-HCN1 KO (blue) mice under photopic conditions with a background light of 70 cd/$m^2$ and a spatial frequency of 0.383 cycles/degree. Animals were tested at frequencies of 1.5, 3, and 6 Hz. Data is presented as mean ± SEM with each individual mouse average shown. For sample size and detailed statistics see Supplemental Table S7.
