## Supplemental Statistical Tables for "Cone-driven retinal responses are shaped by rod but not cone HCN1"

### Main text figures

**Supplemental Table 2.1 related to Figure 2B: Rod-HCN1 KO scotopic a-wave amplitude (µV)**

| | Rod-Control | | | Rod-HCN1 KO | | | Rod-Control - Rod-HCN1 KO | | | |
|---|---|---|---|---|---|---|---|---|---|---|
| cd.s/m$^2$ | mean | st dev | N | mean | st dev | N | mean | 95% CI | adj p | significant |
| 0.003 | 2.142429 | 19.71107 | 7 | 4.973111 | 6.366025829 | 9 | -2.831 | -32.60 to 26.93 | >0.9999 | ns |
| 0.01 | 31.38 | 5.64794 | 7 | 49.20222 | 8.776978805 | 9 | -17.82 | -29.47 to -6.171 | 0.0019 | ** |
| 0.1 | 159.25 | 20.37678 | 7 | 195.5056 | 16.905111 | 9 | -36.26 | -67.90 to -4.606 | 0.0212 | * |
| 1 | 291.3714 | 24.91783 | 7 | 320.1444 | 16.56574863 | 9 | -28.77 | -66.31 to 8.767 | 0.1835 | ns |
| 3 | 244.5571 | 11.90754 | 7 | 267.6056 | 13.91367233 | 9 | -23.05 | -43.79 to -2.307 | 0.025 | * |
| 10 | 287.2071 | 24.03406 | 7 | 313.7278 | 22.43230248 | 9 | -26.52 | -64.95 to 11.91 | 0.295 | ns |
| 30 | 287.2429 | 26.88251 | 7 | 311.3056 | 25.12786606 | 9 | -24.06 | -67.06 to 18.94 | 0.5358 | ns |
| 100 | 335.4714 | 36.15145 | 7 | 369.9333 | 20.55884846 | 9 | -34.46 | -88.50 to 19.58 | 0.3406 | ns |

| ANOVA table | SS | DF | MS | F (DFn, DFd) | P value |
|---|---|---|---|---|---|
| Intensity x Genotype | 3054 | 7 | 436.3 | F (7, 98) = 1.955 | P=0.0690 |
| Intensity | 1888580 | 7 | 269797 | F (2.034, 28.48) = 1209 | P<0.0001 |
| Genotype | 18481 | 1 | 18481 | F (1, 14) = 11.35 | P=0.0046 |
| Subject | 22794 | 14 | 1628 | F (14, 98) = 7.296 | P<0.0001 |
| Residual | 21869 | 98 | 223.2 | | |

**Supplemental Table 2.2 related to Figure 2B: rod-HCN1 KO scotopic b-wave amplitude (µV)**

| | Rod-Control | | | Rod-HCN1 KO | | | Rod-Control - Rod-HCN1 KO | | | |
|---|---|---|---|---|---|---|---|---|---|---|
| cd.s/m$^2$ | mean | st dev | N | mean | st dev | N | mean | 95% CI | adj p | significant |
| 0.003 | 296.65 | 28.20476 | 7 | 300.2778 | 24.90066655 | 9 | -3.628 | -48.05 to 40.80 | >0.9999 | ns |
| 0.01 | 408.2214 | 33.40627 | 7 | 375.0222 | 21.55114717 | 9 | 33.2 | -17.02 to 83.42 | 0.0019 | ** |
| 0.1 | 533.4143 | 54.83626 | 7 | 540.9444 | 37.39456381 | 9 | -7.53 | -90.32 to 75.26 | 0.0212 | * |
| 1 | 719.85 | 59.772 | 7 | 754.7444 | 40.25659453 | 9 | -34.89 | -125.0 to 55.25 | 0.1835 | ns |
| 3 | 724.3286 | 68.37495 | 7 | 763.8389 | 49.44314145 | 9 | -39.51 | -143.4 to 64.37 | 0.025 | * |
| 10 | 720.0429 | 84.11092 | 7 | 754.8278 | 80.43498582 | 9 | -34.78 | -170.2 to 100.7 | 0.295 | ns |
| 30 | 647.3 | 100.4894 | 7 | 644.8111 | 79.65456978 | 9 | 2.489 | -152.3 to 157.2 | 0.5358 | ns |
| 100 | 808.8143 | 116.6496 | 7 | 806.4833 | 66.62971372 | 9 | 2.331 | -172.1 to 176.7 | 0.3406 | ns |

| ANOVA table | SS | DF | MS | F (DFn, DFd) | P value |
|---|---|---|---|---|---|
| Intensity x Genotype | 17030 | 7 | 2433 | F (7, 98) = 1.388 | P=0.2189 |
| Intensity | 3792372 | 7 | 541767 | F (2.298, 32.17) = 309.1 | P<0.0001 |
| Genotype | 3336 | 1 | 3336 | F (1, 14) = 0.1654 | P=0.6904 |
| Subject | 282323 | 14 | 20166 | F (14, 98) = 11.51 | P<0.0001 |
| Residual | 171772 | 98 | 1753 | | |

**Supplemental Table 2.3 related to Figure 2C: a-wave time to peak (ms)**

| cd.s/m² | Rod-Control | | | Rod-HCN1 KO | | | Rod-Control - Rod-HCN1 KO | | | |
|---|---|---|---|---|---|---|---|---|---|---|
| | mean | st dev | N | mean | st dev | N | mean | 95% CI | adj p | significant |
| 0.003 | 21.42857 | 2.967583 | 7 | 25.58333 | 4.128786141 | 9 | -4.155 | -9.844 to 1.534 | 0.2459 | ns |
| 0.01 | 22.5 | 1.689428 | 7 | 23.80556 | 1.599696152 | 9 | -1.306 | -4.018 to 1.407 | 0.7041 | ns |
| 0.1 | 18.17857 | 1.255938 | 7 | 19.02778 | 1.394433378 | 9 | -0.8492 | -2.987 to 1.288 | 0.8665 | ns |
| 1 | 15 | 1.25 | 7 | 16.19444 | 1.1843892 | 9 | -1.194 | -3.202 to 0.8131 | 0.4639 | ns |
| 3 | 9.321429 | 0.965167 | 7 | 10.16667 | 1.536025716 | 9 | -0.8452 | -2.870 to 1.180 | 0.8337 | ns |
| 10 | 8 | 0.661438 | 7 | 8.25 | 0.760345316 | 9 | -0.25 | -1.393 to 0.8933 | 0.9957 | ns |
| 30 | 6.857143 | 0.475595 | 7 | 6.972222 | 0.403973321 | 9 | -0.1151 | -0.8575 to 0.6273 | 0.9995 | ns |
| 100 | 5.642857 | 0.475595 | 7 | 6.055556 | 0.242956329 | 9 | -0.4127 | -1.123 to 0.2975 | 0.4304 | ns |

| ANOVA table | SS | DF | MS | F (DFn, DFd) | P value |
|---|---|---|---|---|---|
| Intensity x Genotype | 45.92 | 7 | 6.56 | F (7, 98) = 4.264 | P=0.0004 |
| Intensity | 5782 | 7 | 826 | F (1.513, 21.18) = 536.8 | P<0.0001 |
| Genotype | 41 | 1 | 41 | F (1, 14) = 3.636 | P=0.0773 |
| Subject | 157.9 | 14 | 11.28 | F (14, 98) = 7.328 | P<0.0001 |
| Residual | 150.8 | 98 | 1.539 | | |

**Supplemental Table 2.4 related to Figure 2C: b-wave time to peak (ms)**

| cd.s/m² | Rod-Control | | | Rod-HCN1 KO | | | Rod-Control - Rod-HCN1 KO | | | |
|---|---|---|---|---|---|---|---|---|---|---|
| | mean | st dev | N | mean | st dev | N | mean | 95% CI | adj p | significant |
| 0.003 | 53.78571 | 6.392425 | 7 | 57.63889 | 7.002355754 | 9 | -3.853 | -14.67 to 6.964 | 0.9204 | ns |
| 0.01 | 39.10714 | 2.597503 | 7 | 41.77778 | 2.895158794 | 9 | -2.671 | -7.099 to 1.758 | 0.4563 | ns |
| 0.1 | 28.32143 | 1.951038 | 7 | 31.11111 | 2.945099508 | 9 | -2.79 | -6.734 to 1.155 | 0.2768 | ns |
| 1 | 26.14286 | 2.135304 | 7 | 27.33333 | 2.080414622 | 9 | -1.19 | -4.650 to 2.269 | 0.9307 | ns |
| 3 | 24 | 2.577951 | 7 | 25.19444 | 2.23529143 | 9 | -1.194 | -5.237 to 2.849 | 0.9679 | ns |
| 10 | 26.28571 | 2.631653 | 7 | 27.47222 | 2.501388503 | 9 | -1.187 | -5.417 to 3.044 | 0.9775 | ns |
| 30 | 27.60714 | 3.210845 | 7 | 31.36111 | 6.550418774 | 9 | -3.754 | -11.97 to 4.457 | 0.7484 | ns |
| 100 | 25.17857 | 2.907298 | 7 | 27.52778 | 3.241441723 | 9 | -2.349 | -7.306 to 2.608 | 0.7281 | ns |

| ANOVA table | SS | DF | MS | F (DFn, DFd) | P value |
|---|---|---|---|---|---|
| Intensity x Genotype | 33.69 | 7 | 4.813 | F (7, 98) = 0.6858 | P=0.6838 |
| Intensity | 12344 | 7 | 1763 | F (2.445, 34.23) = 251.3 | P<0.0001 |
| Genotype | 177.5 | 1 | 177.5 | F (1, 14) = 2.692 | P=0.1231 |
| Subject | 922.8 | 14 | 65.91 | F (14, 98) = 9.392 | P<0.0001 |
| Residual | 687.7 | 98 | 7.018 | | |

**Supplemental Table 2.5 related to Figure 2D: amplitude 100 ms post flash (µV)**

| cd.s/m$^2$ | Rod-Control mean | Rod-Control st dev | N | Rod-HCN1 KO mean | Rod-HCN1 KO st dev | N | Rod-Control - Rod-HCN1 KO mean | 95% CI | adj p | significant |
|---|---|---|---|---|---|---|---|---|---|---|
| 0.003 | 368.6369 | 49.60996 | 7 | 417.0267 | 50.57801097 | 9 | -48.39 | -130.0 to 33.24 | 0.4729 | ns |
| 0.01 | 349.9117 | 39.28257 | 7 | 438.9241 | 28.94185499 | 9 | -89.01 | -148.8 to -29.19 | 0.0033 | ** |
| 0.1 | 189.8169 | 20.60044 | 7 | 306.4628 | 30.74612939 | 9 | -116.6 | -158.0 to -75.32 | <0.0001 | **** |
| 1 | 186.1405 | 33.27181 | 7 | 313.8773 | 27.48012646 | 9 | -127.7 | -179.4 to -76.11 | <0.0001 | **** |
| 3 | 180.2049 | 22.80942 | 7 | 299.3378 | 33.78252501 | 9 | -119.1 | -164.7 to -73.61 | <0.0001 | **** |
| 10 | 143.2031 | 41.04597 | 7 | 246.8248 | 32.85075389 | 9 | -103.6 | -166.9 to -40.31 | 0.0014 | ** |
| 30 | 172.9495 | 29.66218 | 7 | 204.972 | 36.9157365 | 9 | -32.02 | -85.35 to 21.31 | 0.4642 | ns |
| 100 | 241.8504 | 63.02082 | 7 | 284.0138 | 59.95206596 | 9 | -42.16 | -143.5 to 59.17 | 0.83 | ns |

| ANOVA table | SS | DF | MS | F (DFn, DFd) | P value |
|---|---|---|---|---|---|
| Intensity x Genotype | 40704 | 7 | 5815 | F (7, 98) = 4.393 | P=0.0003 |
| Intensity | 702147 | 7 | 100307 | F (2.346, 32.85) = 75.78 | P<0.0001 |
| Genotype | 226735 | 1 | 226735 | F (1, 14) = 72.10 | P<0.0001 |
| Subject | 44023 | 14 | 3145 | F (14, 98) = 2.376 | P=0.0069 |
| Residual | 129719 | 98 | 1324 | | |

**Supplemental Table 2.6 related to Figure 2D: amplitude 150 ms post flash (µV)**

| cd.s/m$^2$ | Rod-Control mean | Rod-Control st dev | N | Rod-HCN1 KO mean | Rod-HCN1 KO st dev | N | Rod-Control - Rod-HCN1 KO mean | 95% CI | adj p | significant |
|---|---|---|---|---|---|---|---|---|---|---|
| 0.003 | 138.8324 | 48.86097 | 7 | 227.8495 | 56.12080004 | 9 | -89.02 | -173.4 to -4.590 | 0.0357 | * |
| 0.01 | 151.8354 | 43.57117 | 7 | 303.4182 | 40.01065833 | 9 | -151.6 | -220.9 to -82.24 | <0.0001 | **** |
| 0.1 | 83.86192 | 22.00704 | 7 | 171.84 | 31.20692608 | 9 | -87.98 | -130.7 to -45.28 | <0.0001 | **** |
| 1 | 74.88416 | 44.24306 | 7 | 189.3981 | 22.01146591 | 9 | -114.5 | -180.6 to -48.44 | 0.0016 | ** |
| 3 | 94.66356 | 18.82412 | 7 | 204.1751 | 21.89617934 | 9 | -109.5 | -142.2 to -76.79 | <0.0001 | **** |
| 10 | 58.87832 | 40.02074 | 7 | 178.6583 | 25.49668979 | 9 | -119.8 | -179.9 to -59.67 | 0.0004 | *** |
| 30 | 100.9358 | 29.04662 | 7 | 158.4265 | 33.78626086 | 9 | -57.49 | -108.0 to -7.004 | 0.021 | * |
| 100 | 167.2868 | 80.35195 | 7 | 219.1114 | 63.90743316 | 9 | -51.82 | -175.6 to 71.98 | 0.8133 | ns |

| ANOVA table | SS | DF | MS | F (DFn, DFd) | P value |
|---|---|---|---|---|---|
| Intensity x Genotype | 30336 | 7 | 4334 | F (7, 98) = 2.822 | P=0.0101 |
| Intensity | 168854 | 7 | 24122 | F (2.217, 31.04) = 15.71 | P<0.0001 |
| Genotype | 300753 | 1 | 300753 | F (1, 14) = 91.67 | P<0.0001 |
| Subject | 45930 | 14 | 3281 | F (14, 98) = 2.136 | P=0.0158 |
| Residual | 150484 | 98 | 1536 | | |

**Supplemental Table 2.7 related to Figure 2D: amplitude 200 ms post flash (μV)**

| | Rod-Control | | | Rod-HCN1 KO | | | Rod-Control - Rod-HCN1 KO | | | |
|---|---|---|---|---|---|---|---|---|---|---|
| cd.s/m$^2$ | mean | st dev | N | mean | st dev | N | mean | 95% CI | adj p | significant |
| 0.003 | 25.89355 | 23.25215 | 7 | 31.60213 | 26.19387859 | 9 | -5.709 | -45.54 to 34.12 | 0.9998 | ns |
| 0.01 | 82.25778 | 27.01809 | 7 | 61.95363 | 46.09224373 | 9 | 20.3 | -39.39 to 80.00 | 0.936 | ns |
| 0.1 | 22.41113 | 23.73584 | 7 | 87.59731 | 18.14876073 | 9 | -65.19 | -101.5 to -28.85 | 0.0007 | *** |
| 1 | 34.60326 | 29.2558 | 7 | 114.4266 | 21.49604854 | 9 | -79.82 | -124.4 to -35.29 | 0.0007 | *** |
| 3 | 75.46384 | 14.74884 | 7 | 148.9172 | 10.55642697 | 9 | -73.45 | -95.83 to -51.07 | <0.0001 | **** |
| 10 | 52.35867 | 22.8127 | 7 | 133.7399 | 18.02396117 | 9 | -81.38 | -116.5 to -46.27 | <0.0001 | **** |
| 30 | 93.28995 | 23.28766 | 7 | 121.3715 | 29.28759817 | 9 | -28.08 | -70.19 to 14.02 | 0.3412 | ns |
| 100 | 117.4964 | 71.7496 | 7 | 180.6713 | 59.28606304 | 9 | -63.17 | -174.5 to 48.18 | 0.5088 | ns |

| ANOVA table | SS | DF | MS | F (DFn, DFd) | P value |
|---|---|---|---|---|---|
| Intensity x Genotype | 39943 | 7 | 5706 | F (7, 98) = 5.215 | P<0.0001 |
| Intensity | 153274 | 7 | 21896 | F (2.381, 33.33) = 20.01 | P<0.0001 |
| Genotype | 69771 | 1 | 69771 | F (1, 14) = 66.13 | P<0.0001 |
| Subject | 14771 | 14 | 1055 | F (14, 98) = 0.9643 | P=0.4950 |
| Residual | 107226 | 98 | 1094 | | |

**Supplemental Table 3.1 related to Figure 3B: amplitude (μV)**

| | Rod-Control | | | Rod-HCN1 KO | | | Rod-Control - Rod HCN1 KO | | | |
|---|---|---|---|---|---|---|---|---|---|---|
| Hz | mean | st dev | N | mean | st dev | N | mean | 95% CI | adj p | significant |
| 0.5 | 73.947 | 15.03549 | 5 | 125.2833 | 18.17464351 | 6 | -51.34 | -89.32 to -13.36 | 0.0074 | ** |
| 0.75 | 99.29366 | 26.49955 | 5 | 140.3917 | 15.31243979 | 6 | -41.1 | -100.1 to 17.95 | 0.2278 | ns |
| 1 | 96.682 | 25.97926 | 5 | 144.9333 | 12.51266026 | 6 | -48.25 | -107.4 to 10.93 | 0.1182 | ns |
| 1.5 | 108.242 | 24.07347 | 5 | 141.8361 | 15.27508862 | 6 | -33.59 | -86.85 to 19.66 | 0.3287 | ns |
| 2 | 114.5053 | 37.03039 | 5 | 167.6445 | 12.68271148 | 6 | -53.14 | -141.7 to 35.39 | 0.3035 | ns |
| 3 | 104.2233 | 36.26997 | 5 | 153.9778 | 13.08907396 | 6 | -49.75 | -135.9 to 36.35 | 0.3425 | ns |
| 6 | 98.74132 | 29.98207 | 5 | 126.45 | 10.13139041 | 6 | -27.71 | -99.51 to 44.09 | 0.7467 | ns |
| 9 | 61.102 | 19.34847 | 5 | 86.51445 | 5.564438402 | 6 | -25.41 | -72.66 to 21.84 | 0.3897 | ns |
| 12 | 33.944 | 7.832244 | 5 | 46.44723 | 2.044487986 | 6 | -12.5 | -31.82 to 6.818 | 0.2291 | ns |
| 18 | 6.42514 | 1.709206 | 5 | 7.397883 | 1.662251629 | 6 | -0.9727 | -4.915 to 2.969 | 0.9959 | ns |
| 24 | 4.38066 | 1.228726 | 5 | 2.544783 | 1.125451942 | 6 | 1.836 | -0.9534 to 4.625 | 0.3274 | ns |
| 30 | 3.52288 | 0.894843 | 5 | 3.006783 | 1.006445902 | 6 | 0.5161 | -1.664 to 2.696 | 0.9974 | ns |

| ANOVA table | SS | DF | MS | F (DFn, DFd) | P value |
|---|---|---|---|---|---|
| Frequency x Genotype | 13476 | 11 | 1225 | F (11, 99) = 7.710 | P<0.0001 |
| Frequency | 344288 | 11 | 31299 | F (1.776, 15.98) = 197.0 | P<0.0001 |
| Genotype | 26492 | 1 | 26492 | F (1, 9) = 14.67 | P=0.0040 |
| Subject | 16256 | 9 | 1806 | F (9, 99) = 11.37 | P<0.0001 |
| Residual | 15730 | 99 | 158.9 | | |

**Supplemental Table 3.2 related to Figure 3D: amplitude (µV)**

| Hz | Rod-Control mean | Rod-Control st dev | N | Rod-HCN1 KO mean | Rod-HCN1 KO st dev | N | Rod-Control - Rod HCN1 KO mean | 95% CI | adj p | significant |
|---|---|---|---|---|---|---|---|---|---|---|
| 0.5 | 142.81 | 46.77147 | 5 | 153.4333 | 31.445042 | 6 | -10.62 | -113.8 to 92.59 | >0.9999 | ns |
| 0.75 | 159.9187 | 51.61292 | 5 | 116.9083 | 9.176296457 | 6 | 43.01 | -88.04 to 174.1 | 0.8272 | ns |
| 1 | 147.517 | 45.57596 | 5 | 113.1533 | 11.45563035 | 6 | 34.36 | -78.48 to 147.2 | 0.8913 | ns |
| 1.5 | 135.4093 | 40.95045 | 5 | 142.3583 | 11.17235054 | 6 | -6.949 | -107.5 to 93.62 | >0.9999 | ns |
| 2 | 108.8466 | 34.237 | 5 | 126.6 | 9.601505877 | 6 | -17.75 | -101.6 to 66.09 | 0.9898 | ns |
| 3 | 82.57268 | 27.62486 | 5 | 120.0644 | 13.52698648 | 6 | -37.49 | -100.3 to 25.30 | 0.3476 | ns |
| 6 | 65.86598 | 17.50769 | 5 | 114.8222 | 13.95244666 | 6 | -48.96 | -87.80 to -10.11 | 0.0135 | * |
| 9 | 48.81268 | 17.11814 | 5 | 105.3972 | 13.75623908 | 6 | -56.58 | -94.60 to -18.57 | 0.0047 | ** |
| 12 | 38.2733 | 10.89306 | 5 | 48.90388 | 5.470200731 | 6 | -10.63 | -35.31 to 14.05 | 0.7077 | ns |
| 18 | 14.9744 | 2.4347 | 5 | 11.72138 | 1.127018384 | 6 | 3.253 | -2.325 to 8.831 | 0.3633 | ns |
| 24 | 5.7008 | 2.316908 | 5 | 4.55655 | 1.300047848 | 6 | 1.144 | -4.034 to 6.323 | 0.9956 | ns |
| 30 | 3.6308 | 1.024842 | 5 | 2.936283 | 1.068366357 | 6 | 0.6945 | -1.725 to 3.114 | 0.9865 | ns |

| ANOVA table | SS | DF | MS | F (DFn, DFd) | P value |
|---|---|---|---|---|---|
| Frequency x Genotype | 26430 | 11 | 2403 | F (11, 99) = 8.790 | P<0.0001 |
| Frequency | 361279 | 11 | 32844 | F (2.029, 18.26) = 120.2 | P<0.0001 |
| Genotype | 2579 | 1 | 2579 | F (1, 9) = 0.8250 | P=0.3874 |
| Subject | 28134 | 9 | 3126 | F (9, 99) = 11.44 | P<0.0001 |
| Residual | 27062 | 99 | 273.4 | | |

**Supplemental Table 4.1 related to Figure 4B: amplitude (μV)**

| | Rod-Control | | | Rod-HCN1 KO | | | Rod-Control - Rod HCN1 KO | | | |
|---|---|---|---|---|---|---|---|---|---|---|
| Hz | mean | st dev | N | mean | st dev | N | mean | 95% CI | adj p | significant |
| 0.5 | 107.512 | 44.67225 | 5 | 116.0536 | 51.6436552 | 7 | -8.542 | -112.7 to 95.65 | >0.9999 | ns |
| 0.75 | 83.886 | 40.3998 | 5 | 64.94141 | 20.44427788 | 7 | 18.94 | -73.22 to 111.1 | 0.9965 | ns |
| 1 | 77.347 | 29.66952 | 5 | 54.02714 | 19.63712574 | 7 | 23.32 | -42.09 to 88.73 | 0.8967 | ns |
| 1.5 | 88.02834 | 28.38278 | 5 | 23.51553 | 10.24627536 | 7 | 64.51 | -3.503 to 132.5 | 0.0618 | ns |
| 2 | 73.636 | 27.56347 | 5 | 34.18857 | 15.85320518 | 7 | 39.45 | -22.30 to 101.2 | 0.2949 | ns |
| 3 | 69.95 | 25.06298 | 5 | 34.1262 | 12.67875281 | 7 | 35.82 | -21.36 to 93.00 | 0.2973 | ns |
| 6 | 55.65532 | 23.67338 | 5 | 33.71129 | 15.30496165 | 7 | 21.94 | -30.36 to 74.25 | 0.7721 | ns |
| 9 | 48.03666 | 18.9939 | 5 | 22.5661 | 10.88482176 | 7 | 25.47 | -17.10 to 68.04 | 0.3588 | ns |
| 12 | 42.948 | 17.86043 | 5 | 8.769943 | 1.862958424 | 7 | 34.18 | -12.10 to 80.46 | 0.1412 | ns |
| 18 | 23.2518 | 8.607204 | 5 | 7.351257 | 1.863302968 | 7 | 15.9 | -5.795 to 37.60 | 0.1496 | ns |
| 24 | 14.59872 | 5.748964 | 5 | 4.7948 | 1.118474264 | 7 | 9.804 | -4.782 to 24.39 | 0.1951 | ns |
| 30 | 9.83034 | 2.653332 | 5 | 1.356157 | 1.277736534 | 7 | 8.474 | 2.376 to 14.57 | 0.011 | * |

| ANOVA table | SS | DF | MS | F (DFn, DFd) | P value |
|---|---|---|---|---|---|
| Frequency x Genotype | 10858 | 11 | 987.1 | F (11, 110) = 3.609 | P=0.0002 |
| Frequency | 118200 | 11 | 10745 | F (1.767, 17.67) = 39.29 | P<0.0001 |
| Genotype | 20339 | 1 | 20339 | F (1, 10) = 7.202 | P=0.0229 |
| Subject | 28242 | 10 | 2824 | F (10, 110) = 10.33 | P<0.0001 |
| Residual | 30083 | 110 | 273.5 | | |

**Supplemental Table 4.2 related to Figure 4D: amplitude (µV)**

| Hz | Rod-Control mean | Rod-Control st dev | N | Rod-HCN1 KO mean | Rod-HCN1 KO st dev | N | Rod-Control - Rod HCN1 KO mean | 95% CI | adj p | significant |
|---|---|---|---|---|---|---|---|---|---|---|
| 0.5 | 49.869 | 19.25759 | 5 | 60.2985 | 30.40235339 | 7 | -10.43 | -63.33 to 42.48 | 0.9996 | ns |
| 0.75 | 61.08 | 23.76964 | 5 | 30.70989 | 18.93399323 | 7 | 30.37 | -21.61 to 82.35 | 0.4429 | ns |
| 1 | 54.037 | 28.19038 | 5 | 33.49786 | 11.56661498 | 7 | 20.54 | -45.82 to 86.90 | 0.9137 | ns |
| 1.5 | 78.171 | 32.69592 | 5 | 36.81167 | 15.04765548 | 7 | 41.36 | -34.30 to 117.0 | 0.4191 | ns |
| 2 | 82.274 | 35.25692 | 5 | 38.22571 | 12.27743459 | 7 | 44.05 | -40.84 to 128.9 | 0.4366 | ns |
| 3 | 73.38134 | 35.19597 | 5 | 33.08381 | 10.08459426 | 7 | 40.3 | -46.36 to 127.0 | 0.5305 | ns |
| 6 | 64.43666 | 26.30174 | 5 | 24.82456 | 10.80620037 | 7 | 39.61 | -22.29 to 101.5 | 0.2591 | ns |
| 9 | 52.49266 | 19.10842 | 5 | 14.87461 | 10.38597531 | 7 | 37.62 | -5.522 to 80.76 | 0.0909 | ns |
| 12 | 49.92732 | 23.59977 | 5 | 11.078 | 6.096305411 | 7 | 38.85 | -19.83 to 97.53 | 0.2109 | ns |
| 18 | 29.04332 | 9.45528 | 5 | 7.4122 | 1.457055983 | 7 | 21.63 | -2.618 to 45.88 | 0.0754 | ns |
| 24 | 19.25464 | 4.723523 | 5 | 4.343414 | 2.263166404 | 7 | 14.91 | 4.046 to 25.78 | 0.0117 | * |
| 30 | 8.5725 | 3.212282 | 5 | 1.3592 | 1.391759216 | 7 | 7.213 | -0.2882 to 14.71 | 0.0589 | ns |

| ANOVA table | SS | DF | MS | F (DFn, DFd) | P value |
|---|---|---|---|---|---|
| Frequency x Genotype | 9059 | 11 | 823.6 | F (11, 110) = 5.803 | P<0.0001 |
| Frequency | 44111 | 11 | 4010 | F (2.879, 28.79) = 28.25 | P<0.0001 |
| Genotype | 25834 | 1 | 25834 | F (1, 10) = 10.15 | P=0.0097 |
| Subject | 25455 | 10 | 2546 | F (10, 110) = 17.94 | P<0.0001 |
| Residual | 15612 | 110 | 141.9 | | |

**Supplemental Table 5 related to Figure 5B: amplitude (μV)**

| | Rod-Control | | | Rod-HCN1 KO | | | Rod-Control - Rod-HCN1 KO | | | |
|---|---|---|---|---|---|---|---|---|---|---|
| Hz | mean | st dev | N | mean | st dev | N | mean | 95% CI | adj p | significant |
| 0.5 | 477.5625 | 66.55822 | 4 | 458.5833 | 50.26661583 | 6 | 18.98 | -168.8 to 206.7 | >0.9999 | ns |
| 1 | 320.7625 | 41.28616 | 4 | 135.6367 | 29.26686038 | 6 | 185.1 | 67.08 to 303.2 | 0.0067 | ** |
| 2 | 169.135 | 19.5547 | 4 | 31.8335 | 29.61918467 | 6 | 137.3 | 75.92 to 198.7 | 0.0003 | *** |
| 3 | 152.1425 | 31.26696 | 4 | 13.68992 | 3.233121811 | 6 | 138.5 | 17.02 to 259.9 | 0.0343 | * |
| 5 | 157.7106 | 20.53444 | 4 | 21.88158 | 9.521539242 | 6 | 135.8 | 69.81 to 201.8 | 0.0035 | ** |
| 7 | 147.3363 | 17.80757 | 4 | 14.994 | 5.843316687 | 6 | 132.3 | 70.12 to 194.6 | 0.0041 | ** |
| 10 | 120.2063 | 11.06577 | 4 | 11.71717 | 4.018543065 | 6 | 108.5 | 70.64 to 146.3 | 0.0014 | ** |
| 12 | 113.7913 | 9.447318 | 4 | 14.32017 | 3.138873391 | 6 | 99.47 | 66.54 to 132.4 | 0.0012 | ** |
| 15 | 94.705 | 15.13573 | 4 | 17.4885 | 13.19003219 | 6 | 77.22 | 35.39 to 119.0 | 0.0022 | ** |
| 18 | 73.8525 | 13.20825 | 4 | 11.13458 | 5.307188648 | 6 | 62.72 | 18.62 to 106.8 | 0.015 | * |
| 20 | 61.81125 | 8.782899 | 4 | 9.63275 | 3.547347118 | 6 | 52.18 | 22.89 to 81.47 | 0.0067 | ** |
| 30 | 48.91475 | 14.37856 | 4 | 6.462667 | 1.00928255 | 6 | 42.45 | -13.80 to 98.71 | 0.1089 | ns |

| ANOVA table | SS | DF | MS | F (DFn, DFd) | P value |
|---|---|---|---|---|---|
| Frequency x Genotype | 63800 | 11 | 5800 | F (11, 88) = 12.51 | P<0.0001 |
| Frequency | 1612931 | 11 | 146630 | F (1.731, 13.85) = 316.2 | P<0.0001 |
| Genotype | 283484 | 1 | 283484 | F (1, 8) = 232.0 | P<0.0001 |
| Subject | 9774 | 8 | 1222 | F (8, 88) = 2.634 | P=0.0123 |
| Residual | 40812 | 88 | 463.8 | | |

**Supplemental Table 6.1: related to Figure 6B: a-wave amplitude (μV)**

| | Rod-Control | | | Rod-HCN1 KO | | | Rod-Control - Rod-HCN1 KO | | | |
|---|---|---|---|---|---|---|---|---|---|---|
| $cd.s/m^2$ | mean | st dev | N | mean | st dev | N | mean | 95% CI | adj p | significant |
| 0.23 | 2.155625 | 5.304819 | 8 | 8.891045 | 9.246536783 | 11 | -6.735 | -17.24 to 3.768 | 0.4002 | ns |
| 1 | 6.997125 | 4.944541 | 8 | 5.100364 | 4.778097504 | 11 | 1.897 | -5.282 to 9.075 | 0.9864 | ns |
| 2.4 | 10.35081 | 6.404693 | 8 | 8.641182 | 5.143478158 | 11 | 1.71 | -7.185 to 10.60 | 0.9981 | ns |
| 7.6 | 13.08863 | 1.807457 | 8 | 12.198 | 4.563019039 | 11 | 0.8906 | -3.976 to 5.757 | 0.9988 | ns |
| 24 | 16.66188 | 3.577363 | 8 | 16.19764 | 7.74497 | 11 | 0.4642 | -7.954 to 8.882 | >0.9999 | ns |
| 77 | 22.61563 | 4.885213 | 8 | 17.80314 | 5.199704579 | 11 | 4.812 | -2.515 to 12.14 | 0.3691 | ns |
| 246 | 22.23813 | 4.578621 | 8 | 19.30227 | 3.529867111 | 11 | 2.936 | -3.378 to 9.250 | 0.738 | ns |
| 724 | 22.92563 | 4.84294 | 8 | 24.30091 | 5.738443525 | 11 | -1.375 | -8.971 to 6.220 | 0.999 | ns |

| ANOVA table | SS | DF | MS | F (DFn, DFd) | P value |
|---|---|---|---|---|---|
| Intensity x Genotype | 388.7 | 7 | 55.53 | F (7, 119) = 2.046 | P=0.0548 |
| Intensity | 6280 | 7 | 897.1 | F (3.589, 61.01) = 33.05 | P<0.0001 |
| Genotype | 12.24 | 1 | 12.24 | F (1, 17) = 0.2324 | P=0.6359 |
| Subject | 895.6 | 17 | 52.68 | F (17, 119) = 1.941 | P=0.0206 |
| Residual | 3230 | 119 | 27.14 | | |

**Supplemental Table 6.2 related to Figure 6C: b-wave amplitude (μV)**

| | Rod-Control | | | Rod-HCN1 KO | | | Rod-Control - Rod-HCN1 KO | | | |
|---|---|---|---|---|---|---|---|---|---|---|
| $cd.s/m^2$ | mean | st dev | N | mean | st dev | N | mean | 95% CI | adj p | significant |
| 0.23 | 16.94938 | 7.851929 | 8 | 10.48764 | 9.057170568 | 11 | 6.462 | -5.706 to 18.63 | 0.6273 | ns |
| 1 | 40.50688 | 23.45804 | 8 | 13.83505 | 8.273079362 | 11 | 26.67 | -4.694 to 58.04 | 0.1104 | ns |
| 2.4 | 67.21313 | 15.0234 | 8 | 16.80964 | 13.46092423 | 11 | 50.4 | 29.04 to 71.76 | <0.0001 | **** |
| 7.6 | 144.7506 | 28.67422 | 8 | 24.49168 | 14.17610187 | 11 | 120.3 | 81.97 to 158.5 | <0.0001 | **** |
| 24 | 160.7375 | 31.7789 | 8 | 25.51409 | 7.54574046 | 11 | 135.2 | 92.48 to 178.0 | <0.0001 | **** |
| 77 | 164.3375 | 30.31873 | 8 | 30.27 | 16.33176598 | 11 | 134.1 | 93.51 to 174.6 | <0.0001 | **** |
| 246 | 183.3875 | 30.82767 | 8 | 23.898 | 14.95964498 | 11 | 159.5 | 118.3 to 200.6 | <0.0001 | **** |
| 724 | 191.9813 | 33.96499 | 8 | 27.74727 | 10.71655247 | 11 | 164.2 | 118.7 to 209.7 | <0.0001 | **** |

| ANOVA table | SS | DF | MS | F (DFn, DFd) | P value |
|---|---|---|---|---|---|
| Intensity x Genotype | 125338 | 7 | 17905 | F (7, 119) = 125.3 | P<0.0001 |
| Intensity | 184966 | 7 | 26424 | F (3.189, 54.22) = 184.9 | P<0.0001 |
| Genotype | 367578 | 1 | 367578 | F (1, 17) = 179.6 | P<0.0001 |
| Subject | 34795 | 17 | 2047 | F (17, 119) = 14.32 | P<0.0001 |
| Residual | 17005 | 119 | 142.9 | | |

**Supplemental Table 7.1: related to Figure 7A: OMR contrast sensitivity (0.002 cd/m2)**

| | Rod-Control | | | Rod-HCN1 KO | | | Rod-Control - Rod-HCN1 KO | | | |
|---|---|---|---|---|---|---|---|---|---|---|
| Temporal Frequency (Hz) | log mean | st dev | N | log mean | st dev | N | mean | t | adj p | significant |
| 1.488 | 0.6558 | 0.137373 | 6 | 0.557817 | 0.164722669 | 6 | 0.098 | 1.07 | 0.293 | ns |
| 0.372 | 0.8052 | 0.161448 | 6 | 0.76545 | 0.103716667 | 6 | 0.0398 | 0.434 | 0.667 | ns |
| 0.124 | 0.600167 | 0.241986 | 6 | 0.595767 | 0.097889788 | 6 | 0.00437 | 0.0477 | 0.962 | ns |

| ANOVA table | SS | DF | MS | F | P value |
|---|---|---|---|---|---|
| Intensity x frequency | 2 | 0.0134 | 0.0067 | 0.266 | 0.768 |
| Frequency | 2 | 0.268 | 0.134 | 5.329 | 0.01 |
| Genotype | 1 | 0.0202 | 0.0202 | 0.803 | 0.377 |
| Residual | 30 | 0.755 | 0.0252 | | |
| Total | 35 | 1.057 | 0.0302 | | |

**Supplemental Table 7.2: related to Figure 7B: OMR contrast sensitivity (70 cd/m2)**

| | Rod-Control | | | Rod-HCN1 KO | | | Rod-Control - Rod-HCN1 KO | | | |
|---|---|---|---|---|---|---|---|---|---|---|
| Temporal Frequency (Hz) | log mean | st dev | N | log mean | st dev | N | mean | t | adj p | significant |
| 1.488 | 0.624667 | 0.049754 | 6 | 0.733483 | 0.182915581 | 6 | 0.109 | 1.546 | 0.133 | ns |
| 0.372 | 0.833767 | 0.138716 | 6 | 0.78875 | 0.113055486 | 6 | 0.045 | 0.64 | 0.527 | ns |
| 0.124 | 0.698217 | 0.089763 | 6 | 0.48485 | 0.114808619 | 6 | 0.213 | 3.03 | 0.005 | ** |

| ANOVA table | SS | DF | MS | F | P value |
|---|---|---|---|---|---|
| Intensity x frequency | 2 | 0.156 | 0.0779 | 5.239 | 0.011 |
| Frequency | 2 | 0.294 | 0.147 | 9.876 | <0.001 |
| Genotype | 1 | 0.0224 | 0.0224 | 1.504 | 0.23 |
| Residual | 30 | 0.446 | 0.0149 | | |
| Total | 35 | 0.918 | 0.0262 | | |

**Supplemental Table 9.1 related to Figure 9B: a-wave amplitude (μV)**

| | Cone-Control | | | Cone-HCN1 KO | | | Cone-Control - Cone HCN1 KO | | | |
|---|---|---|---|---|---|---|---|---|---|---|
| $cd.s/m^2$ | mean | st dev | N | mean | st dev | N | mean | 95% CI | adj p | significant |
| 0.23 | 2.969556 | 3.02582 | 9 | 2.745944 | 3.358551766 | 9 | 0.2236 | -4.593 to 5.040 | >0.9999 | ns |
| 1 | 5.729278 | 1.775084 | 9 | 5.223111 | 3.481097302 | 9 | 0.5062 | -3.877 to 4.889 | >0.9999 | ns |
| 2.4 | 10.95767 | 3.562094 | 9 | 10.92794 | 4.224100396 | 9 | 0.02972 | -5.873 to 5.933 | >0.9999 | ns |
| 7.6 | 14.46883 | 3.514285 | 9 | 13.16694 | 4.571393841 | 9 | 1.302 | -4.893 to 7.497 | 0.9983 | ns |
| 24 | 18.48128 | 3.981286 | 9 | 16.83272 | 3.879822379 | 9 | 1.649 | -4.265 to 7.562 | 0.9877 | ns |
| 77 | 18.47944 | 4.28484 | 9 | 19.46 | 4.436766841 | 9 | -0.9806 | -7.543 to 5.582 | 0.9999 | ns |
| 246 | 22.67039 | 6.803215 | 9 | 20.22556 | 4.435262425 | 9 | 2.445 | -6.411 to 11.30 | 0.9869 | ns |
| 724 | 25.62667 | 6.158576 | 9 | 24.14822 | 7.534705183 | 9 | 1.478 | -8.935 to 11.89 | >0.9999 | ns |

| ANOVA table | SS | DF | MS | F (DFn, DFd) | P value |
|---|---|---|---|---|---|
| Intensity x Genotype | 41.86 | 8 | 5.232 | F (8, 128) = 0.3331 | P=0.9518 |
| Intensity | 12316 | 8 | 1540 | F (4.349, 69.58) = 98.02 | P<0.0001 |
| Genotype | 20.67 | 1 | 20.67 | F (1, 16) = 0.4208 | P=0.5257 |
| Subject | 785.8 | 16 | 49.11 | F (16, 128) = 3.127 | P=0.0002 |
| Residual | 2010 | 128 | 15.71 | | |

**Supplemental Table 9.2 related to Figure 9B: b-wave amplitude (μV)**

| | Cone-Control | | | Cone-HCN1 KO | | | Cone-Control - Cone HCN1 KO | | | |
|---|---|---|---|---|---|---|---|---|---|---|
| $cd.s/m^2$ | mean | st dev | N | mean | st dev | N | mean | 95% CI | adj p | significant |
| 0.23 | 14.62794 | 5.528958 | 9 | 14.17106 | 3.765202123 | 9 | 0.4569 | -6.674 to 7.588 | >0.9999 | ns |
| 1 | 37.57 | 11.97675 | 9 | 31.33222 | 7.256910978 | 9 | 6.238 | -8.871 to 21.35 | 0.8389 | ns |
| 2.4 | 70.40333 | 24.86712 | 9 | 58.79 | 18.00779623 | 9 | 11.61 | -20.94 to 44.16 | 0.9235 | ns |
| 7.6 | 127.2494 | 32.61878 | 9 | 114.6983 | 27.16817244 | 9 | 12.55 | -32.03 to 57.13 | 0.9805 | ns |
| 24 | 139.2789 | 41.67986 | 9 | 122.47 | 32.13636774 | 9 | 16.81 | -38.72 to 72.34 | 0.9694 | ns |
| 77 | 150.4033 | 33.86394 | 9 | 130.9833 | 31.19511069 | 9 | 19.42 | -28.74 to 67.58 | 0.8685 | ns |
| 246 | 159.7033 | 39.53317 | 9 | 145.2094 | 34.96360683 | 9 | 14.49 | -40.78 to 69.77 | 0.9876 | ns |
| 724 | 180.8461 | 48.26683 | 9 | 155.7856 | 36.42818452 | 9 | 25.06 | -38.84 to 88.96 | 0.8802 | ns |

| ANOVA table | SS | DF | MS | F (DFn, DFd) | P value |
|---|---|---|---|---|---|
| Intensity x Genotype | 1835 | 7 | 262.1 | F (7, 112) = 1.047 | P=0.4027 |
| Intensity | 418604 | 7 | 59801 | F (1.521, 24.33) = 238.9 | P<0.0001 |
| Genotype | 6397 | 1 | 6397 | F (1, 16) = 1.183 | P=0.2928 |
| Subject | 86502 | 16 | 5406 | F (16, 112) = 21.60 | P<0.0001 |
| Residual | 28038 | 112 | 250.3 | | |

**Supplemental Table 9.3 related to Figure 9C: a-wave time to peak (ms)**

| cd.s/m$^2$ | Cone-Control | | | Cone-HCN1 KO | | | Cone-Control - Cone HCN1 KO | | | |
|---|---|---|---|---|---|---|---|---|---|---|
| | mean | st dev | N | mean | st dev | N | mean | 95% CI | adj p | significant |
| 0.23 | 19.02778 | 5.287728 | 9 | 20.72222 | 4.049391244 | 9 | -1.694 | -8.725 to 5.336 | 0.9925 | ns |
| 1 | 18.05556 | 2.576267 | 9 | 18.86111 | 2.385211964 | 9 | -0.8056 | -4.478 to 2.867 | 0.9962 | ns |
| 2.4 | 18 | 2.899353 | 9 | 18.55556 | 1.379412113 | 9 | -0.5556 | -4.118 to 3.007 | 0.9995 | ns |
| 7.6 | 15 | 0.856957 | 9 | 16.22222 | 1.6414763 | 9 | -1.222 | -3.254 to 0.8100 | 0.4451 | ns |
| 24 | 13.41667 | 1.045825 | 9 | 14.47222 | 1.201850425 | 9 | -1.056 | -2.725 to 0.6141 | 0.4138 | ns |
| 77 | 12.25 | 0.984251 | 9 | 12.69444 | 0.596575878 | 9 | -0.4444 | -1.686 to 0.7973 | 0.9169 | ns |
| 246 | 11.47222 | 1.20185 | 9 | 11.80556 | 0.596575878 | 9 | -0.3333 | -1.815 to 1.148 | 0.9938 | ns |
| 724 | 10.16667 | 0.216506 | 9 | 10.66667 | 0.450693909 | 9 | -0.5 | -1.054 to 0.05412 | 0.0887 | ns |

| ANOVA table | SS | DF | MS | F (DFn, DFd) | P value |
|---|---|---|---|---|---|
| Intensity x Genotype | 6.894 | 7 | 0.9849 | F (7, 112) = 0.2272 | P=0.9781 |
| Intensity | 1559 | 7 | 222.7 | F (2.189, 35.02) = 51.37 | P<0.0001 |
| Genotype | 24.59 | 1 | 24.59 | F (1, 16) = 3.142 | P=0.0953 |
| Subject | 125.2 | 16 | 7.824 | F (16, 112) = 1.805 | P=0.0388 |
| Residual | 485.5 | 112 | 4.335 | | |

**Supplemental Table 9.4 related to Figure 9C: b-wave time to peak (ms)**

| cd.s/m$^2$ | Cone-Control | | | Cone-HCN1 KO | | | Cone-Control - Cone HCN1 KO | | | |
|---|---|---|---|---|---|---|---|---|---|---|
| | mean | st dev | N | mean | st dev | N | mean | 95% CI | adj p | significant |
| 0.23 | 36.5 | 3.535534 | 9 | 37.60714 | 1.375811449 | 7 | -1.107 | -5.444 to 3.230 | 0.985 | ns |
| 1 | 40.72222 | 7.982837 | 9 | 37.91667 | 6.101997624 | 9 | 2.806 | -7.802 to 13.41 | 0.9864 | ns |
| 2.4 | 42.41667 | 6.159342 | 9 | 44.88889 | 3.644583119 | 9 | -2.472 | -10.21 to 5.270 | 0.9537 | ns |
| 7.6 | 41.94444 | 4.003471 | 9 | 44.77778 | 2.716973398 | 9 | -2.833 | -7.993 to 2.326 | 0.5721 | ns |
| 24 | 39.69444 | 3.633133 | 9 | 42.27778 | 2.198879134 | 9 | -2.583 | -7.166 to 1.999 | 0.5329 | ns |
| 77 | 39.86111 | 6.369006 | 9 | 40.05556 | 1.788757328 | 9 | -0.1944 | -7.923 to 7.534 | >0.9999 | ns |
| 246 | 36.5 | 3.041381 | 9 | 38.55556 | 1.721998193 | 9 | -2.056 | -5.855 to 1.744 | 0.5763 | ns |
| 724 | 34.69444 | 2.524189 | 9 | 36.66667 | 1.789727633 | 9 | -1.972 | -5.259 to 1.314 | 0.4684 | ns |

Mixed-effects model

| Fixed effects (type III) | P value | P value sur | Statistically | F (DFn, DFd) | Geisser-Greenhouse's epsilon |
|---|---|---|---|---|---|
| Intensity | <0.0001 | **** | Yes | F (1.991, 31.29) = 14.25 | 0.2845 |
| Genotype | 0.3547 | ns | No | F (1, 16) = 0.9086 | |
| Intensity x Genotype | 0.1732 | ns | No | F (7, 110) = 1.504 | |

| Random effects | SD | Variance |
|---|---|---|
| Subject | 2.598 | 6.75 |
| Residual | 3.242 | 10.51 |

**Supplemental Table 9.5 related to Figure 9D: amplitude 100 ms post flash (µV)**

| $cd.s/m^2$ | Cone-Control | | | Cone-HCN1 KO | | | Cone-Control - Cone HCN1 KO | | | |
|---|---|---|---|---|---|---|---|---|---|---|
| | mean | st dev | N | mean | st dev | N | mean | 95% CI | adj p | significant |
| 0.23 | 11.56443 | 16.60269 | 9 | 1.63324 | 8.935951533 | 9 | 9.931 | -10.69 to 30.55 | 0.6993 | ns |
| 1 | -2.530668 | 24.73266 | 9 | -2.005667 | 7.345093986 | 9 | -0.525 | -30.54 to 29.49 | >0.9999 | ns |
| 2.4 | -9.021187 | 8.321828 | 9 | -15.44444 | 11.93438893 | 9 | 6.423 | -9.053 to 21.90 | 0.8423 | ns |
| 7.6 | -12.74602 | 22.2246 | 9 | -8.312808 | 16.85765156 | 9 | -4.433 | -33.90 to 25.03 | 0.9997 | ns |
| 24 | 4.367302 | 10.4749 | 9 | 1.927949 | 11.23671428 | 9 | 2.439 | -13.63 to 18.50 | 0.9997 | ns |
| 77 | 40.80709 | 22.32406 | 9 | 46.95683 | 14.05855283 | 9 | -6.15 | -34.50 to 22.20 | 0.9959 | ns |
| 246 | 80.92096 | 31.67354 | 9 | 98.4837 | 23.09215037 | 9 | -17.56 | -59.10 to 23.97 | 0.8312 | ns |
| 724 | 105.7741 | 39.7288 | 9 | 110.2858 | 24.89316494 | 9 | -4.512 | -54.91 to 45.89 | >0.9999 | ns |

| ANOVA table | SS | DF | MS | F (DFn, DFd) | P value |
|---|---|---|---|---|---|
| Intensity x Genotype | 2279 | 7 | 325.6 | F (7, 112) = 0.9561 | P=0.4669 |
| Intensity | 280054 | 7 | 40008 | F (3.172, 50.74) = 117.5 | P<0.0001 |
| Genotype | 116.5 | 1 | 116.5 | F (1, 16) = 0.1220 | P=0.7315 |
| Subject | 15277 | 16 | 954.8 | F (16, 112) = 2.804 | P=0.0008 |
| Residual | 38143 | 112 | 340.6 | | |

**Supplemental Table 9.6 related to Figure 9D: amplitude 150 ms post flash (µV)**

| $cd.s/m^2$ | Cone-Control | | | Cone-HCN1 KO | | | Cone-Control - Cone HCN1 KO | | | |
|---|---|---|---|---|---|---|---|---|---|---|
| | mean | st dev | N | mean | st dev | N | mean | 95% CI | adj p | significant |
| 0.23 | 10.39693 | 20.07318 | 9 | 2.530199 | 6.946075289 | 9 | 7.867 | -16.52 to 32.25 | 0.9375 | ns |
| 1 | 6.168491 | 24.07933 | 9 | 9.669185 | 14.66061849 | 9 | -3.501 | -33.90 to 26.90 | >0.9999 | ns |
| 2.4 | 2.350781 | 14.00563 | 9 | -3.406787 | 19.51625767 | 9 | 5.758 | -19.73 to 31.25 | 0.9949 | ns |
| 7.6 | -1.000099 | 25.01369 | 9 | -4.077285 | 13.31778238 | 9 | 3.077 | -27.95 to 34.11 | >0.9999 | ns |
| 24 | 7.473344 | 19.16236 | 9 | -10.42678 | 16.33904713 | 9 | 17.9 | -8.514 to 44.31 | 0.3322 | ns |
| 77 | 10.48504 | 34.1844 | 9 | -0.903391 | 16.69590109 | 9 | 11.39 | -30.69 to 53.47 | 0.9802 | ns |
| 246 | 33.69312 | 14.90545 | 9 | 37.82417 | 17.48855361 | 9 | -4.131 | -28.23 to 19.97 | 0.9993 | ns |
| 724 | 75.37418 | 32.88882 | 9 | 88.92356 | 26.11660212 | 9 | -13.55 | -57.77 to 30.67 | 0.9674 | ns |

| ANOVA table | SS | DF | MS | F (DFn, DFd) | P value |
|---|---|---|---|---|---|
| Intensity x Genotype | 3108 | 7 | 443.9 | F (7, 112) = 1.233 | P=0.2905 |
| Intensity | 107418 | 7 | 15345 | F (2.930, 46.89) = 42.64 | P<0.0001 |
| Genotype | 346.2 | 1 | 346.2 | F (1, 16) = 0.3532 | P=0.5606 |
| Subject | 15683 | 16 | 980.2 | F (16, 112) = 2.723 | P=0.0011 |
| Residual | 40311 | 112 | 359.9 | | |

**Supplemental Table 9.7 related to Figure 9D: amplitude 200 ms post flash (µV)**

| cd.s/m$^2$ | Cone-Control | | | Cone-HCN1 KO | | | Cone-Control - Cone HCN1 KO | | | |
|---|---|---|---|---|---|---|---|---|---|---|
| | mean | st dev | N | mean | st dev | N | mean | 95% CI | adj p | significant |
| 0.23 | 6.787196 | 17.17775 | 9 | 0.194447 | 13.00633973 | 9 | -16.17 to 29.36 | | 0.9762 | ns |
| 1 | -0.271912 | 20.50053 | 9 | -5.90986 | 11.31754071 | 9 | -19.89 to 31.17 | | 0.9949 | ns |
| 2.4 | -1.269383 | 14.38806 | 9 | -9.094441 | 12.10764552 | 9 | -11.91 to 27.56 | | 0.8769 | ns |
| 7.6 | 0.096353 | 21.77493 | 9 | -14.69915 | 10.96259163 | 9 | -12.07 to 41.67 | | 0.5463 | ns |
| 24 | 6.865423 | 17.94966 | 9 | -14.67516 | 11.65882368 | 9 | -1.383 to 44.46 | | 0.0725 | ns |
| 77 | 1.727729 | 22.79126 | 9 | -6.952172 | 17.13759782 | 9 | -21.46 to 38.82 | | 0.9769 | ns |
| 246 | 13.41339 | 10.29675 | 9 | 10.61999 | 18.34638189 | 9 | -20.10 to 25.69 | | >0.9999 | ns |
| 724 | 43.73402 | 19.33088 | 9 | 43.39839 | 23.98346024 | 9 | -32.07 to 32.74 | | >0.9999 | ns |

| ANOVA table | SS | DF | MS | F (DFn, DFd) | P value |
|---|---|---|---|---|---|
| Intensity x Genotype | 1446 | 7 | 206.5 | F (7, 112) = 0.7228 | P=0.6529 |
| Intensity | 35918 | 7 | 5131 | F (4.161, 66.58) = 17.96 | P<0.0001 |
| Genotype | 2616 | 1 | 2616 | F (1, 16) = 8.393 | P=0.0105 |
| Subject | 4988 | 16 | 311.7 | F (16, 112) = 1.091 | P=0.3720 |
| Residual | 31999 | 112 | 285.7 | | |

**Supplemental Table 10.1 related to Figure 10B: 3cd.s/m2 flash flicker amplitude (μV)**

| Hz | Cone-Control | | | Cone-HCN1 KO | | | Cone-Control - Cone HCN1 KO | | | |
|---|---|---|---|---|---|---|---|---|---|---|
| | mean | st dev | N | mean | st dev | N | mean | 95% CI | adj p | significant |
| 0.5 | 486.1438 | 104.2111 | 8 | 435.0591 | 77.70898216 | 11 | 51.08 | -101.3 to 203.5 | 0.9747 | ns |
| 1 | 364.7438 | 86.50585 | 8 | 335.2136 | 60.38884454 | 11 | 29.53 | -96.04 to 155.1 | 0.9986 | ns |
| 2 | 229.5306 | 56.85062 | 8 | 225.1368 | 59.83873228 | 11 | 4.394 | -85.76 to 94.55 | >0.9999 | ns |
| 3 | 197.782 | 29.4951 | 8 | 179.6705 | 38.15042362 | 11 | 18.11 | -33.11 to 69.34 | 0.9729 | ns |
| 5 | 185.8475 | 29.98932 | 8 | 169.4482 | 36.65812934 | 11 | 16.4 | -34.23 to 67.03 | 0.986 | ns |
| 7 | 158.7097 | 20.49293 | 8 | 148.8396 | 29.49522381 | 11 | 9.87 | -27.94 to 47.68 | 0.9979 | ns |
| 10 | 141.7275 | 21.00553 | 8 | 121.5947 | 21.76457902 | 11 | 20.13 | -13.00 to 53.26 | 0.522 | ns |
| 12 | 125.5706 | 20.12505 | 8 | 116.6753 | 23.33269138 | 11 | 8.895 | -24.27 to 42.06 | 0.9972 | ns |
| 15 | 89.51125 | 10.29024 | 8 | 83.65455 | 16.56922033 | 11 | 5.857 | -14.57 to 26.28 | 0.995 | ns |
| 18 | 73.17969 | 9.600178 | 8 | 63.47941 | 15.36557699 | 11 | 9.7 | -9.276 to 28.68 | 0.7522 | ns |
| 20 | 59.02313 | 8.62371 | 8 | 54.38127 | 12.62053215 | 11 | 4.642 | -11.43 to 20.72 | 0.9948 | ns |
| 30 | 32.128 | 4.521464 | 8 | 33.33795 | 9.114041226 | 11 | -1.21 | -11.86 to 9.436 | >0.9999 | ns |

| ANOVA table | SS | DF | MS | F (DFn, DFd) | P value |
|---|---|---|---|---|---|
| Frequency x Genotype | 10229 | 11 | 929.9 | F (11, 187) = 0.9504 | P=0.4935 |
| Frequency | 3216439 | 11 | 292404 | F (1.664, 28.29) = 298.9 | P<0.0001 |
| Genotype | 12148 | 1 | 12148 | F (1, 17) = 1.157 | P=0.2971 |
| Subject | 178435 | 17 | 10496 | F (17, 187) = 10.73 | P<0.0001 |
| Residual | 182955 | 187 | 978.4 | | |

**Supplemental Table 10.2 related to Figure 10D: photopic sinusoidal flicker amplitude (μV)**

| Hz | Cone-Control | | | Cone-HCN1 KO | | | Cone-Control - Cone HCN1 KO | | | |
|---|---|---|---|---|---|---|---|---|---|---|
| | mean | st dev | N | mean | st dev | N | mean | 95% CI | adj p | significant |
| 0.5 | 39.0225 | 13.06926 | 6 | 45.89167 | 15.36812795 | 6 | -6.869 | -36.93 to 23.19 | 0.9977 | ns |
| 0.75 | 40.00625 | 4.512552 | 6 | 45.57597 | 7.503623781 | 6 | -5.57 | -19.33 to 8.195 | 0.847 | ns |
| 1 | 40.36861 | 3.906007 | 6 | 41.88272 | 6.230128703 | 6 | -1.514 | -12.98 to 9.949 | >0.9999 | ns |
| 1.5 | 43.7135 | 4.18824 | 6 | 45.03383 | 6.776187524 | 6 | -1.32 | -13.77 to 11.13 | >0.9999 | ns |
| 2 | 43.10944 | 4.384918 | 6 | 46.22 | 11.07222168 | 6 | -3.111 | -23.58 to 17.36 | 0.9998 | ns |
| 3 | 40.99483 | 6.087678 | 6 | 46.54417 | 8.279556568 | 6 | -5.549 | -21.13 to 10.03 | 0.9331 | ns |
| 6 | 38.65183 | 4.321849 | 6 | 42.57417 | 6.622737 | 6 | -3.922 | -16.16 to 8.311 | 0.9617 | ns |
| 9 | 31.61444 | 2.744669 | 6 | 33.45445 | 5.425004452 | 6 | -1.84 | -11.76 to 8.079 | 0.9993 | ns |
| 12 | 27.53361 | 2.087894 | 6 | 30.78722 | 2.827185701 | 6 | -3.254 | -8.578 to 2.071 | 0.4242 | ns |
| 15 | 21.39208 | 3.273195 | 6 | 23.16167 | 2.779944769 | 6 | -1.77 | -8.170 to 4.631 | 0.9892 | ns |
| 25 | 8.967167 | 0.935324 | 6 | 11.63067 | 1.127342125 | 6 | -2.664 | -4.851 to -0.4757 | 0.0146 | * |

| ANOVA table | SS | DF | MS | F (DFn, DFd) | P value |
|---|---|---|---|---|---|
| Frequency x Genotype | 105.8 | 10 | 10.58 | F (10, 100) = 0.3544 | P=0.9629 |
| Frequency | 14979 | 10 | 1498 | F (2.353, 23.53) = 50.18 | P<0.0001 |
| Genotype | 381.1 | 1 | 381.1 | F (1, 10) = 1.985 | P=0.1892 |
| Subject | 1920 | 10 | 192 | F (10, 100) = 6.434 | P<0.0001 |
| Residual | 2985 | 100 | 29.85 | | |

**Supplemental Table 11.1 related to Figure 11B: a-wave amplitude (µV)**

| cd.s/m | Cone-Control | | | Cone-HCN1 KO | | | Cone-Control - Cone HCN1 KO | | | |
|---|---|---|---|---|---|---|---|---|---|---|
| | mean | st dev | N | mean | st dev | N | mean | 95% CI | adj p | significant |
| 57.5 | 5.998417 | 0.88154 | 6 | 7.032714 | 2.290713805 | 7 | -1.034 | -4.385 to 2.317 | 0.9194 | ns |
| 250 | 8.559417 | 1.551734 | 6 | 9.6115 | 1.929523581 | 7 | -1.052 | -4.225 to 2.121 | 0.9172 | ns |
| 600 | 12.90725 | 2.679398 | 6 | 14.07571 | 0.906611956 | 7 | -1.168 | -5.735 to 3.398 | 0.9496 | ns |
| 1900 | 16.4095 | 3.174181 | 6 | 16.82357 | 1.671326106 | 7 | -0.4141 | -5.715 to 4.887 | >0.9999 | ns |
| 6000 | 17.13333 | 3.124011 | 6 | 18.33786 | 1.474920498 | 7 | -1.205 | -6.437 to 4.028 | 0.9768 | ns |
| 19250 | 21.17833 | 5.008789 | 6 | 21.81357 | 3.43465393 | 7 | -0.6352 | -9.074 to 7.804 | >0.9999 | ns |
| 25000 | 22.68583 | 3.141844 | 6 | 24.82786 | 3.260517414 | 7 | -2.142 | -8.003 to 3.719 | 0.8717 | ns |

| ANOVA table | SS | DF | MS | F (DFn, DFd) | P value |
|---|---|---|---|---|---|
| Intensity x Genotype | 5.797 | 6 | 0.9661 | F (6, 66) = 0.2421 | P=0.9608 |
| Intensity | 3053 | 6 | 508.9 | F (3.106, 34.17) = 127.5 | P<0.0001 |
| Genotype | 27.02 | 1 | 27.02 | F (1, 11) = 1.041 | P=0.3296 |
| Subject | 285.5 | 11 | 25.96 | F (11, 66) = 6.504 | P<0.0001 |
| Residual | 263.4 | 66 | 3.991 | | |

**Supplemental Table 11.2 related to Figure 11B: b-wave amplitude (µV)**

| cd.s/m | Cone-Control | | | Cone-HCN1 KO | | | Cone-Control - Cone HCN1 KO | | | |
|---|---|---|---|---|---|---|---|---|---|---|
| | mean | st dev | N | mean | st dev | N | mean | 95% CI | adj p | significant |
| 57.5 | 32.54333 | 6.679682 | 6 | 29.26357 | 3.056031896 | 7 | 3.28 | -7.923 to 14.48 | 0.9224 | ns |
| 250 | 81.4 | 12.20843 | 6 | 73.74571 | 9.453125334 | 7 | 7.654 | -13.25 to 28.56 | 0.8565 | ns |
| 600 | 98.9125 | 16.23439 | 6 | 88.40214 | 13.13286522 | 7 | 10.51 | -17.52 to 38.54 | 0.8454 | ns |
| 1900 | 115.3067 | 22.16818 | 6 | 107.19 | 18.50844559 | 7 | 8.117 | -30.42 to 46.65 | 0.9916 | ns |
| 6000 | 142.1975 | 29.48823 | 6 | 129.4779 | 21.04946829 | 7 | 12.72 | -37.18 to 62.62 | 0.9724 | ns |
| 19250 | 150.0683 | 32.3882 | 6 | 134.3379 | 21.8450879 | 7 | 15.73 | -38.76 to 70.22 | 0.9458 | ns |
| 25000 | 151.3758 | 37.25075 | 6 | 134.5293 | 20.2460141 | 7 | 16.85 | -45.35 to 79.04 | 0.9528 | ns |

| ANOVA table | SS | DF | MS | F (DFn, DFd) | P value |
|---|---|---|---|---|---|
| Intensity x Genotype | 446.5 | 6 | 74.42 | F (6, 66) = 0.5534 | P=0.7656 |
| Intensity | 131630 | 6 | 21938 | F (1.608, 17.68) = 163.1 | P<0.0001 |
| Genotype | 2586 | 1 | 2586 | F (1, 11) = 1.182 | P=0.3001 |
| Subject | 24062 | 11 | 2187 | F (11, 66) = 16.27 | P<0.0001 |
| Residual | 8875 | 66 | 134.5 | | |

**Supplemental Table 11.3 related to Figure 11C: a-wave time to peak (ms)**

| cd.s/m | Cone-Control | | | Cone-HCN1 KO | | | Cone-Control - Cone HCN1 KO | | | |
|---|---|---|---|---|---|---|---|---|---|---|
| | mean | st dev | N | mean | st dev | N | mean | 95% CI | adj p | significant |
| 57.5 | 19.625 | 3.901122 | 6 | 22.21429 | 3.855268504 | 7 | -2.589 | -9.726 to 4.547 | 0.8742 | ns |
| 250 | 19.66667 | 1.736855 | 6 | 20.85714 | 1.390871601 | 7 | -1.19 | -4.183 to 1.802 | 0.8052 | ns |
| 600 | 19.58333 | 1.169045 | 6 | 20.78571 | 1.509888044 | 7 | -1.202 | -3.648 to 1.244 | 0.6363 | ns |
| 1900 | 19 | 1.557241 | 6 | 19.89286 | 1.078744861 | 7 | -0.8929 | -3.519 to 1.733 | 0.8877 | ns |
| 6000 | 19.16667 | 1.136515 | 6 | 21.03571 | 1.523623501 | 7 | -1.869 | -4.305 to 0.5664 | 0.1826 | ns |
| 19250 | 20.91667 | 0.983192 | 6 | 21.39286 | 1.125991626 | 7 | -0.4762 | -2.397 to 1.445 | 0.9811 | ns |
| 25000 | 20.66667 | 1.200694 | 6 | 21.85714 | 1.329607891 | 7 | -1.19 | -3.497 to 1.116 | 0.5852 | ns |

| ANOVA table | SS | DF | MS | F (DFn, DFd) | P value |
|---|---|---|---|---|---|
| Intensity x Genotype | 9.208 | 6 | 1.535 | F (6, 66) = 0.5281 | P=0.7850 |
| Intensity | 33.68 | 6 | 5.614 | F (2.111, 23.23) = 1.932 | P=0.1658 |
| Genotype | 40.87 | 1 | 40.87 | F (1, 11) = 4.972 | P=0.0475 |
| Subject | 90.43 | 11 | 8.221 | F (11, 66) = 2.829 | P=0.0044 |
| Residual | 191.8 | 66 | 2.906 | | |

**Supplemental Table 11.4 related to Figure 11C: b-wave time to peak (ms)**

| cd.s/m | Cone-Control | | | Cone-HCN1 KO | | | Cone-Control - Cone HCN1 KO | | | |
|---|---|---|---|---|---|---|---|---|---|---|
| | mean | st dev | N | mean | st dev | N | mean | 95% CI | adj p | significant |
| 57.5 | 49.83333 | 2.969287 | 6 | 51 | 6.044970361 | 7 | -1.167 | -10.08 to 7.751 | 0.9995 | ns |
| 250 | 54.29167 | 2.129652 | 6 | 56.32143 | 1.902379462 | 7 | -2.03 | -5.797 to 1.738 | 0.5277 | ns |
| 600 | 56.875 | 5.245832 | 6 | 61.96429 | 3.423048627 | 7 | -5.089 | -13.89 to 3.714 | 0.4195 | ns |
| 1900 | 59.08333 | 10.80471 | 6 | 64.71429 | 8.052469305 | 7 | -5.631 | -24.02 to 12.76 | 0.933 | ns |
| 6000 | 56.54167 | 2.436271 | 6 | 60.28571 | 2.980591983 | 7 | -3.744 | -8.678 to 1.190 | 0.1919 | ns |
| 19250 | 57.375 | 5.667341 | 6 | 58.17857 | 2.6208868 | 7 | -0.8036 | -10.30 to 8.697 | >0.9999 | ns |
| 25000 | 55.125 | 2.914404 | 6 | 60.21429 | 5.544194733 | 7 | -5.089 | -13.32 to 3.139 | 0.3649 | ns |

| ANOVA table | SS | DF | MS | F (DFn, DFd) | P value |
|---|---|---|---|---|---|
| Intensity x Genotype | 78.83 | 6 | 13.14 | F (6, 66) = 0.5771 | P=0.7472 |
| Intensity | 1016 | 6 | 169.3 | F (2.622, 28.84) = 7.436 | P=0.0012 |
| Genotype | 256 | 1 | 256 | F (1, 11) = 5.661 | P=0.0365 |
| Subject | 497.5 | 11 | 45.23 | F (11, 66) = 1.987 | P=0.0438 |
| Residual | 1503 | 66 | 22.77 | | |

**Supplemental Table 11.5 related to Figure 11D: amplitude 150 ms post onset (μV)**

| cd.s/m | Cone-Control | | | Cone-HCN1 KO | | | Cone-Control - Cone HCN1 KO | | | |
|---|---|---|---|---|---|---|---|---|---|---|
| | mean | st dev | N | mean | st dev | N | mean | 95% CI | adj p | significant |
| 57.5 | 0.016983 | 5.942589 | 6 | -1.106568 | 10.41651531 | 7 | 1.124 | -14.50 to 16.74 | >0.9999 | ns |
| 250 | 30.55715 | 11.46024 | 6 | 24.52926 | 8.562593928 | 7 | 6.028 | -13.48 to 25.54 | 0.9303 | ns |
| 600 | 40.64609 | 15.83829 | 6 | 45.72274 | 12.18474903 | 7 | -5.077 | -32.16 to 22.01 | 0.9955 | ns |
| 1900 | 63.50355 | 15.13467 | 6 | 75.78252 | 14.04170185 | 7 | -12.28 | -39.36 to 14.80 | 0.7085 | ns |
| 6000 | 74.57188 | 18.06685 | 6 | 79.95179 | 12.40412161 | 7 | -5.38 | -35.82 to 25.06 | 0.9965 | ns |
| 19250 | 82.71765 | 22.19735 | 6 | 84.23606 | 10.82064171 | 7 | -1.518 | -38.66 to 35.62 | >0.9999 | ns |
| 25000 | 73.13669 | 25.7355 | 6 | 80.48854 | 17.15762222 | 7 | -7.352 | -50.61 to 35.90 | 0.9971 | ns |

| ANOVA table | SS | DF | MS | F (DFn, DFd) | P value |
|---|---|---|---|---|---|
| Intensity x Genotype | 691.4 | 6 | 115.2 | F (6, 66) = 0.8757 | P=0.5177 |
| Intensity | 77114 | 6 | 12852 | F (2.912, 32.03) = 97.66 | P<0.0001 |
| Genotype | 276 | 1 | 276 | F (1, 11) = 0.3567 | P=0.5624 |
| Subject | 8511 | 11 | 773.7 | F (11, 66) = 5.880 | P<0.0001 |
| Residual | 8686 | 66 | 131.6 | | |

**Supplemental Table 11.6 related to Figure 11D: amplitude 250 ms post onset (μV)**

| cd.s/m | Cone-Control | | | Cone-HCN1 KO | | | Cone-Control - Cone HCN1 KO | | | |
|---|---|---|---|---|---|---|---|---|---|---|
| | mean | st dev | N | mean | st dev | N | mean | 95% CI | adj p | significant |
| 57.5 | 5.000794 | 6.064873 | 6 | 10.49105 | 15.65717659 | 7 | -5.49 | -28.40 to 17.42 | 0.9771 | ns |
| 250 | 26.13638 | 10.84979 | 6 | 26.73742 | 13.95438013 | 7 | -0.601 | -23.24 to 22.04 | >0.9999 | ns |
| 600 | 32.19188 | 11.56786 | 6 | 40.58723 | 14.1540501 | 7 | -8.395 | -31.83 to 15.03 | 0.8833 | ns |
| 1900 | 47.36031 | 13.83062 | 6 | 60.30127 | 12.58959795 | 7 | -12.94 | -37.54 to 11.66 | 0.5554 | ns |
| 6000 | 41.88053 | 5.625767 | 6 | 55.27626 | 16.05192756 | 7 | -13.4 | -36.88 to 10.09 | 0.4178 | ns |
| 19250 | 50.21709 | 25.38731 | 6 | 66.50438 | 19.80082098 | 7 | -16.29 | -59.81 to 27.24 | 0.844 | ns |
| 25000 | 41.96162 | 24.53776 | 6 | 61.09302 | 25.18545818 | 7 | -19.13 | -64.69 to 26.43 | 0.7794 | ns |

| ANOVA table | SS | DF | MS | F (DFn, DFd) | P value |
|---|---|---|---|---|---|
| Intensity x Genotype | 803.7 | 6 | 134 | F (6, 66) = 0.7565 | P=0.6066 |
| Intensity | 25472 | 6 | 4245 | F (2.697, 29.67) = 23.97 | P<0.0001 |
| Genotype | 2683 | 1 | 2683 | F (1, 11) = 3.074 | P=0.1073 |
| Subject | 9599 | 11 | 872.6 | F (11, 66) = 4.928 | P<0.0001 |
| Residual | 11687 | 66 | 177.1 | | |

**Supplemental Table 11.7 related to Figure 11D: amplitude 500 ms post onset (µV)**

| cd.s/m | Cone-Control | | | Cone-HCN1 KO | | | Cone-Control - Cone HCN1 KO | | | |
|---|---|---|---|---|---|---|---|---|---|---|
| | mean | st dev | N | mean | st dev | N | mean | 95% CI | adj p | significant |
| 57.5 | 0.957643 | 12.78205 | 6 | 4.452842 | 15.09097158 | 7 | -3.495 | -28.87 to 21.88 | 0.9995 | ns |
| 250 | -10.61032 | 12.73099 | 6 | -7.452198 | 28.72645047 | 7 | -3.158 | -45.30 to 38.98 | >0.9999 | ns |
| 600 | -11.61492 | 13.30253 | 6 | 1.269819 | 13.0149597 | 7 | -12.88 | -37.13 to 11.36 | 0.5487 | ns |
| 1900 | -4.267244 | 12.70015 | 6 | 6.564782 | 13.71057128 | 7 | -10.83 | -34.94 to 13.27 | 0.7235 | ns |
| 6000 | -19.82973 | 14.28057 | 6 | -7.609468 | 21.84756368 | 7 | -12.22 | -45.84 to 21.40 | 0.8708 | ns |
| 19250 | -11.87322 | 23.59211 | 6 | 4.012451 | 24.46249287 | 7 | -15.89 | -59.88 to 28.11 | 0.878 | ns |
| 25000 | -11.72813 | 26.17798 | 6 | 2.015059 | 30.95957668 | 7 | -13.74 | -65.76 to 38.28 | 0.9734 | ns |

| ANOVA table | SS | DF | MS | F (DFn, DFd) | P value |
|---|---|---|---|---|---|
| Intensity x Genotype | 487.9 | 6 | 81.32 | F (6, 66) = 0.2541 | P=0.9559 |
| Intensity | 2455 | 6 | 409.2 | F (3.308, 36.39) = 1.279 | P=0.2965 |
| Genotype | 2407 | 1 | 2407 | F (1, 11) = 2.623 | P=0.1336 |
| Subject | 10094 | 11 | 917.6 | F (11, 66) = 2.868 | P=0.0039 |
| Residual | 21121 | 66 | 320 | | |

**Supplemental Table 11.8 related to Figure 11E: OFF response amplitude**

| cd.s/m | Cone-Control | | | Cone-HCN1 KO | | | Cone-Control - Cone HCN1 KO | | | |
|---|---|---|---|---|---|---|---|---|---|---|
| | mean | st dev | N | mean | st dev | N | mean | 95% CI | adj p | significant |
| 57.5 | 44.4743 | 9.181629 | 6 | 42.31567 | 6.398569021 | 7 | 2.159 | -13.34 to 17.65 | 0.9992 | ns |
| 250 | 51.00122 | 9.623204 | 6 | 51.65794 | 11.54062073 | 7 | -0.6567 | -19.93 to 18.61 | >0.9999 | ns |
| 600 | 52.42348 | 12.06045 | 6 | 56.15864 | 12.48469598 | 7 | -3.735 | -26.21 to 18.74 | 0.9982 | ns |
| 1900 | 59.49205 | 15.19434 | 6 | 59.84127 | 16.72444957 | 6 | -0.3492 | -31.37 to 30.67 | >0.9999 | ns |
| 6000 | 55.72472 | 11.93188 | 6 | 60.17804 | 7.988476386 | 7 | -4.453 | -24.51 to 15.61 | 0.9863 | ns |
| 19250 | 65.65248 | 17.42346 | 5 | 71.77071 | 4.480491969 | 7 | -6.118 | -43.79 to 31.56 | 0.99 | ns |
| 25000 | 57.71615 | 13.17859 | 6 | 72.8758 | 10.85303796 | 7 | -15.16 | -38.00 to 7.679 | 0.2995 | ns |

Mixed-effects analysis

| Fixed effects (type III) | P value | P value sur | Statistically | F (DFn, DFd) | Geisser-Greenhouse's epsilon |
|---|---|---|---|---|---|
| Intensity | <0.0001 | **** | Yes | F (2.818, 30.06) = 13.94 | 0.4697 |
| Genotype | 0.4009 | ns | No | F (1, 11) = 0.7636 | |
| Intensity x Genotype | 0.1536 | ns | No | F (6, 64) = 1.629 | |

| Random effects | SD | Variance |
|---|---|---|
| Subject | 8.455 | 71.48 |

| Residual | 7.963 | 63.41 |
|---|---|---|

**Supplemental Table 11.9 related to Figure 11F: OFF response time to peak**

| cd.s/m | Cone-Control | | | Cone-HCN1 KO | | | Cone-Control - Cone HCN1 KO | | | |
|---|---|---|---|---|---|---|---|---|---|---|
| | mean | st dev | N | mean | st dev | N | mean | 95% CI | adj p | significant |
| 57.5 | 116.4583 | 8.242294 | 6 | 118.8571 | 13.06462873 | 7 | -2.399 | -22.34 to 17.55 | 0.9998 | ns |
| 250 | 98.08333 | 9.984571 | 6 | 101.5357 | 14.31231155 | 7 | -3.452 | -25.85 to 18.95 | 0.9989 | ns |
| 600 | 97.33333 | 10.71059 | 6 | 88.66667 | 1.411264209 | 6 | 8.667 | -10.20 to 27.53 | 0.5388 | ns |
| 1900 | 101.5833 | 16.22549 | 6 | 102.1429 | 10.10760558 | 7 | -0.5595 | -27.72 to 26.60 | >0.9999 | ns |
| 6000 | 97.625 | 7.801843 | 6 | 96.85714 | 5.782259404 | 7 | 0.7679 | -12.50 to 14.03 | >0.9999 | ns |
| 19250 | 94.54167 | 5.900035 | 6 | 93.60714 | 7.649696539 | 7 | 0.9345 | -11.44 to 13.31 | >0.9999 | ns |
| 25000 | 92.33333 | 6.011794 | 6 | 90.96429 | 6.909276029 | 7 | 1.369 | -10.40 to 13.14 | 0.9998 | ns |

Mixed-effects analysis

| Fixed effects (type III) | P value | P value sur | Statistically | F (DFn, DFd) | Geisser-Greenhouse's epsilon |
|---|---|---|---|---|---|
| Intensity | <0.0001 | **** | Yes | F (3.495, 37.86) = 18.25 | 0.5825 |
| Genotype | 0.8846 | ns | No | F (1, 11) = 0.02206 | |
| Intensity x Genotype | 0.6667 | ns | No | F (6, 65) = 0.6794 | |

| Random effects | SD | Variance |
|---|---|---|
| Subject | 6.278 | 39.41 |
| Residual | 7.39 | 54.61 |

### Supplementary figures

**Supplemental Table S1.1 related to Figure S1A: amplitude of fundamental (μV)**

| | Rod-Control | | | Rod-HCN1 KO | | | Rod-Control - Rod HCN1 KO | | | |
|---|---|---|---|---|---|---|---|---|---|---|
| Hz | mean | st dev | N | mean | st dev | N | mean | 95% CI | adj p | significant |
| 0.5 | 10.2799 | 2.623981 | 5 | 21.29588 | 2.841366345 | 6 | -11.02 | -17.30 to -4.728 | 0.0012 | ** |
| 0.75 | 13.50292 | 3.511623 | 5 | 24.9197 | 2.354262452 | 6 | -11.42 | -19.17 to -3.667 | 0.006 | ** |
| 1 | 17.42472 | 4.556245 | 5 | 27.34618 | 2.52395866 | 6 | -9.921 | -20.12 to 0.2764 | 0.0568 | ns |
| 1.5 | 22.37452 | 6.056916 | 5 | 31.99067 | 3.446993319 | 6 | -9.616 | -23.13 to 3.902 | 0.2094 | ns |
| 2 | 24.33678 | 8.249398 | 5 | 35.43208 | 2.768239537 | 6 | -11.1 | -30.87 to 8.679 | 0.363 | ns |
| 3 | 23.75102 | 8.750894 | 5 | 33.87808 | 3.034311218 | 6 | -10.13 | -31.01 to 10.76 | 0.5164 | ns |
| 6 | 19.56768 | 6.511069 | 5 | 26.34333 | 2.15323645 | 6 | -6.776 | -22.41 to 8.861 | 0.6297 | ns |
| 9 | 13.10654 | 4.195972 | 5 | 17.00877 | 0.825351767 | 6 | -3.902 | -14.49 to 6.688 | 0.7388 | ns |
| 12 | 6.79644 | 2.402441 | 5 | 9.199767 | 0.430807083 | 6 | -2.403 | -8.501 to 3.694 | 0.6708 | ns |
| 18 | 1.15688 | 0.500406 | 5 | 1.267267 | 0.149256714 | 6 | -0.1104 | -1.327 to 1.107 | >0.9999 | ns |
| 24 | 0.4469 | 0.165675 | 5 | 0.250817 | 0.024747963 | 6 | 0.1961 | -0.2281 to 0.6202 | 0.5026 | ns |
| 30 | 0.24066 | 0.103573 | 5 | 0.327867 | 0.04269982 | 6 | -0.08721 | -0.3285 to 0.1541 | 0.829 | ns |

| ANOVA table | SS | DF | MS | F (DFn, DF | P value |
|---|---|---|---|---|---|
| Frequency x Genotype | 682.9 | 11 | 62.08 | F (11, 99) = | P<0.0001 |
| Frequency | 14804 | 11 | 1346 | F (1.483, 1: | P<0.0001 |
| Genotype | 1322 | 1 | 1322 | F (1, 9) = 1 | P=0.0030 |
| Subject | 734.8 | 9 | 81.64 | F (9, 99) = | P<0.0001 |
| Residual | 686 | 99 | 6.929 | | |

**Supplemental Table S1.2 related to Figure S1B: amplitude of fundamental (μV)**

| | Rod-Control | | | Rod-HCN1 KO | | | Rod-Control - Rod HCN1 KO | | | |
|---|---|---|---|---|---|---|---|---|---|---|
| Hz | mean | st dev | N | mean | st dev | N | mean | 95% CI | adj p | significant |
| 0.5 | 27.20804 | 10.18445 | 5 | 27.79125 | 4.98387431 | 6 | -0.5832 | -23.73 to 22.57 | >0.9999 | ns |
| 0.75 | 26.19206 | 9.256377 | 5 | 15.54122 | 1.796477987 | 6 | 10.65 | -12.73 to 34.03 | 0.5289 | ns |
| 1 | 28.48328 | 9.415438 | 5 | 14.29202 | 2.045259214 | 6 | 14.19 | -9.409 to 37.79 | 0.2762 | ns |
| 1.5 | 25.00642 | 9.662977 | 5 | 25.814 | 3.342982099 | 6 | -0.8076 | -23.88 to 22.26 | >0.9999 | ns |
| 2 | 20.86256 | 8.144536 | 5 | 24.49998 | 2.434850774 | 6 | -3.637 | -23.44 to 16.16 | 0.997 | ns |
| 3 | 16.93754 | 5.751995 | 5 | 25.06067 | 2.828473392 | 6 | -8.123 | -21.19 to 4.944 | 0.3092 | ns |
| 6 | 14.86848 | 6.162893 | 5 | 23.96608 | 2.41265186 | 6 | -9.098 | -23.56 to 5.366 | 0.2751 | ns |
| 9 | 10.53218 | 3.835136 | 5 | 21.3979 | 2.449981686 | 6 | -10.87 | -19.35 to -2.385 | 0.0137 | * |
| 12 | 8.11468 | 2.742832 | 5 | 9.785983 | 1.152142855 | 6 | -1.671 | -8.043 to 4.701 | 0.9719 | ns |
| 18 | 3.02474 | 1.01257 | 5 | 2.248683 | 0.421576344 | 6 | 0.7761 | -1.579 to 3.131 | 0.8905 | ns |
| 24 | 1.01788 | 0.365077 | 5 | 0.66725 | 0.090775013 | 6 | 0.3506 | -0.5542 to 1.255 | 0.7092 | ns |
| 30 | 0.29254 | 0.169276 | 5 | 0.2985 | 0.060067962 | 6 | -0.00596 | -0.4087 to 0.3968 | >0.9999 | ns |

| ANOVA table | SS | DF | MS | F (DFn, DF | P value |
|---|---|---|---|---|---|
| Frequency x Genotype | 1617 | 11 | 147 | F (11, 99) = | P<0.0001 |
| Frequency | 11535 | 11 | 1049 | F (1.988, 1 | P<0.0001 |
| Genotype | 17.69 | 1 | 17.69 | F (1, 9) = 0 | P=0.7475 |
| Subject | 1445 | 9 | 160.6 | F (9, 99) = | P<0.0001 |
| Residual | 1037 | 99 | 10.48 | | |

**Supplemental Table S1.3 related to Figure S1D: amplitude of recovery following b-wave-like component (μV)**

| | Rod-Control | | | Rod-HCN1 KO | | | Rod-Control - Rod HCN1 KO | | | |
|---|---|---|---|---|---|---|---|---|---|---|
| Hz | log mean | SEM | N | log mean | SEM | N | mean | t | adj p | significant |
| 0.5 | 2.122 | 0.0638 | 5 | 1.386 | 0.0582 | 6 | 0.736 | 8.521 | <0.001 | **** |
| 0.75 | 2.161 | 0.0638 | 5 | 1.371 | 0.0582 | 6 | 0.79 | 9.146 | <0.001 | **** |
| 1 | 2.153 | 0.0638 | 5 | 1.534 | 0.0582 | 6 | 0.619 | 7.161 | <0.001 | **** |

| **ANOVA table** | **SS** | **DF** | **MS** | **F** | **P** | |
|---|---|---|---|---|---|---|
| Genotype x T Freq | 0.042 | 2 | 0.021 | 1.031 | | 0.37 |
| T Freq | 0.0515 | 2 | 0.0258 | 1.265 | | 0.298 |
| Genotype | 4.183 | 1 | 4.183 | 205.487 | <0.001 | |
| Residual | 0.55 | 27 | 0.0204 | | | |
| Total | 4.835 | 32 | 0.151 | | | |

**Supplemental Table S1.4 related to Figure S1E: rate of recovery following b-wave-like component (μV/ms)**

| | Rod-Control | | | Rod-HCN1 KO | | | Rod-Control - Rod HCN1 KO | | | |
|---|---|---|---|---|---|---|---|---|---|---|
| Hz | mean rate | SEM | N | mean rate | SEM | N | mean | t | adj p | significant |
| 0.5 | -0.525 | 0.0657 | 5 | -0.833 | 0.06 | 6 | 0.308 | 3.463 | 0.002 | *** |
| 0.75 | -0.446 | 0.0657 | 5 | -0.986 | 0.06 | 6 | 0.54 | 6.075 | <0.001 | **** |
| 1 | -0.407 | 0.0657 | 5 | -0.8 | 0.06 | 6 | 0.393 | 4.422 | <0.001 | **** |

| **ANOVA table** | **SS** | **DF** | **MS** | **F** | **P** | |
|---|---|---|---|---|---|---|
| Genotype x T Freq | 2 | 0.0753 | 0.0377 | 1.746 | | 0.194 |
| T Freq | 2 | 0.0711 | 0.0356 | 1.649 | | 0.211 |
| Genotype | 1 | 1.401 | 1.401 | 64.957 | <0.001 | |
| Residual | 27 | 0.582 | 0.0216 | | | |
| Total | 32 | 2.137 | 0.0668 | | | |

**Supplemental Table S2.1 related to Figure S2A: amplitude of fundamental (μV)**

| | Rod-Control | | | Rod-HCN1 KO | | | Rod-Control - Rod HCN1 KO | | | |
|---|---|---|---|---|---|---|---|---|---|---|
| Hz | mean | st dev | N | mean | st dev | N | mean | 95% CI | adj p | significant |
| 0.5 | 14.31078 | 6.481983 | 5 | 27.03853 | 12.94727563 | 7 | -12.73 | -34.12 to 8.669 | 0.4684 | ns |
| 0.75 | 13.2923 | 6.285033 | 5 | 15.8979 | 7.078179665 | 7 | -2.606 | -17.13 to 11.92 | 0.9998 | ns |
| 1 | 15.22442 | 6.873902 | 5 | 8.888686 | 2.945758965 | 7 | 6.336 | -9.743 to 22.41 | 0.752 | ns |
| 1.5 | 18.18418 | 6.688489 | 5 | 2.990886 | 1.388227559 | 7 | 15.19 | -1.712 to 32.10 | 0.0739 | ns |
| 2 | 14.99298 | 6.450274 | 5 | 3.960486 | 2.073937507 | 7 | 11.03 | -4.650 to 26.71 | 0.1803 | ns |
| 3 | 15.05616 | 5.412316 | 5 | 5.155986 | 2.918268544 | 7 | 9.9 | -2.333 to 22.13 | 0.1231 | ns |
| 6 | 12.17234 | 4.844077 | 5 | 6.0765 | 3.185711613 | 7 | 6.096 | -4.590 to 16.78 | 0.4346 | ns |
| 9 | 10.62806 | 4.672774 | 5 | 4.835557 | 2.693969885 | 7 | 5.793 | -4.672 to 16.26 | 0.4437 | ns |
| 12 | 8.73336 | 3.958088 | 5 | 1.149386 | 0.795002795 | 7 | 7.584 | -2.440 to 17.61 | 0.134 | ns |
| 18 | 4.37438 | 1.593963 | 5 | 1.247157 | 0.241865816 | 7 | 3.127 | -0.9630 to 7.217 | 0.1275 | ns |
| 24 | 2.61104 | 1.03547 | 5 | 0.589614 | 0.184143698 | 7 | 2.021 | -0.6180 to 4.661 | 0.1277 | ns |
| 30 | 1.22312 | 0.567171 | 5 | 0.1873 | 0.079987541 | 7 | 1.036 | -0.4232 to 2.495 | 0.1614 | ns |

| ANOVA table | SS | DF | MS | F (DFn, DF | P value |
|---|---|---|---|---|---|
| Frequency x Genotype | 1664 | 11 | 151.3 | F (11, 110) | P<0.0001 |
| Frequency | 4198 | 11 | 381.6 | F (1.779, 1 | P<0.0001 |
| Genotype | 677.2 | 1 | 677.2 | F (1, 10) = | P=0.0440 |
| Subject | 1276 | 10 | 127.6 | F (10, 110) | P<0.0001 |
| Residual | 1517 | 110 | 13.79 | | |

**Supplemental Table S2.2 related to Figure S2B: amplitude of fundamental (μV)**

| | Rod-Control | | | Rod-HCN1 KO | | | Rod-Control - Rod HCN1 KO | | | |
|---|---|---|---|---|---|---|---|---|---|---|
| Hz | mean | st dev | N | mean | st dev | N | mean | 95% CI | adj p | significant |
| 0.5 | 5.4167 | 3.041029 | 5 | 12.56824 | 6.541456818 | 7 | -7.152 | -17.87 to 3.569 | 0.3243 | ns |
| 0.75 | 9.18672 | 5.012848 | 5 | 5.137443 | 3.343540024 | 7 | 4.049 | -6.994 to 15.09 | 0.8815 | ns |
| 1 | 10.25662 | 6.167346 | 5 | 4.693529 | 1.981776176 | 7 | 5.563 | -9.433 to 20.56 | 0.7667 | ns |
| 1.5 | 15.56738 | 6.124965 | 5 | 4.365957 | 3.760332177 | 7 | 11.2 | -2.408 to 24.81 | 0.1192 | ns |
| 2 | 16.1602 | 7.306047 | 5 | 5.372586 | 2.492706784 | 7 | 10.79 | -6.848 to 28.42 | 0.2818 | ns |
| 3 | 14.71584 | 7.282577 | 5 | 5.258714 | 1.829865533 | 7 | 9.457 | -8.694 to 27.61 | 0.4051 | ns |
| 6 | 12.7354 | 5.00241 | 5 | 4.218343 | 2.776510323 | 7 | 8.517 | -2.743 to 19.78 | 0.1615 | ns |
| 9 | 11.35548 | 4.796333 | 5 | 2.621214 | 2.478524064 | 7 | 8.734 | -2.174 to 19.64 | 0.127 | ns |
| 12 | 9.61028 | 4.108676 | 5 | 1.837429 | 1.615567929 | 7 | 7.773 | -1.958 to 17.50 | 0.1204 | ns |
| 18 | 5.1371 | 1.772866 | 5 | 0.937257 | 0.34046406 | 7 | 4.2 | -0.3014 to 8.701 | 0.0645 | ns |
| 24 | 2.59168 | 1.124021 | 5 | 0.500914 | 0.274005094 | 7 | 2.091 | -0.7177 to 4.899 | 0.1437 | ns |
| 30 | 0.8493 | 0.454445 | 5 | 0.167 | 0.078821275 | 7 | 0.6823 | -0.4775 to 1.842 | 0.2846 | ns |

| ANOVA table | SS | DF | MS | F (DFn, DF | P value |
|---|---|---|---|---|---|
| Frequency x Genotype | 873.7 | 11 | 79.43 | F (11, 110) | P<0.0001 |
| Frequency | 1477 | 11 | 134.3 | F (3.287, 3: | P<0.0001 |
| Genotype | 1056 | 1 | 1056 | F (1, 10) = | P=0.0097 |
| Subject | 1040 | 10 | 104 | F (10, 110) | P<0.0001 |
| Residual | 693.2 | 110 | 6.302 | | |

**Supplemental Table S3: related to Figure S3E: Pearson's Correlation**

| Cone-Control | | | Cone-HCN1 KO | | | t-test | |
|---|---|---|---|---|---|---|---|
| mean | st dev | N | mean | st dev | N | p | significant |
| 0.1095 | 0.01471 | 4 | -0.06743 | 0.07698 | 4 | 0.004 | ** |

**Supplemental Table S4.1 related to Figure S4B: a-wave amplitude (μV)**

| | Cone-Control | | | Cone-HCN1 KO | | | Cone-Control - Cone HCN1 KO | | | |
|---|---|---|---|---|---|---|---|---|---|---|
| cd.s/m$^2$ | mean | st dev | N | mean | st dev | N | mean | 95% CI | adj p | significant |
| 0.003 | 1.427813 | 2.402964 | 8 | 0.466111 | 5.922508858 | 9 | 0.9617 | -6.282 to 8.206 | 0.9998 | ns |
| 0.01 | 17.35225 | 8.036624 | 8 | 19.88628 | 6.62236115 | 9 | -2.534 | -14.11 to 9.039 | 0.9956 | ns |
| 0.1 | 113.8344 | 22.81502 | 8 | 118.7689 | 16.91340459 | 9 | -4.935 | -36.95 to 27.08 | 0.9996 | ns |
| 1 | 232.2563 | 28.14433 | 8 | 221.5278 | 35.0022777 | 9 | 10.73 | -37.89 to 59.34 | 0.9958 | ns |
| 3 | 214.2875 | 37.00129 | 8 | 201.3944 | 47.23690589 | 9 | 12.89 | -52.06 to 77.85 | 0.9979 | ns |
| 10 | 233.6313 | 29.10975 | 8 | 220.2278 | 45.85406274 | 9 | 13.4 | -45.84 to 72.65 | 0.9946 | ns |
| 30 | 234.025 | 28.28128 | 8 | 223.2667 | 49.06874642 | 9 | 10.76 | -51.43 to 72.95 | 0.9991 | ns |
| 100 | 277.9375 | 29.39378 | 8 | 268.2 | 56.69085574 | 9 | 9.737 | -61.00 to 80.48 | 0.9998 | ns |

| ANOVA table | SS | DF | MS | F (DFn, DFd) | P value |
|---|---|---|---|---|---|
| Intensity x Genotype | 1601 | 7 | 228.7 | F (7, 105) = 0.4881 | P=0.8414 |
| Intensity | 1285654 | 7 | 183665 | F (1.351, 20.27) = 392.0 | P<0.0001 |
| Genotype | 1378 | 1 | 1378 | F (1, 15) = 0.2583 | P=0.6187 |
| Subject | 80016 | 15 | 5334 | F (15, 105) = 11.39 | P<0.0001 |
| Residual | 49193 | 105 | 468.5 | | |

**Supplemental Table S4.2 related to Figure S4B: b-wave amplitude (μV)**

| | Cone-Control | | | Cone-HCN1 KO | | | Cone-Control - Cone HCN1 KO | | | |
|---|---|---|---|---|---|---|---|---|---|---|
| cd.s/m$^2$ | mean | st dev | N | mean | st dev | N | mean | 95% CI | adj p | significant |
| 0.003 | 239.9438 | 60.9299 | 8 | 224.8056 | 38.12498725 | 9 | 15.14 | -68.02 to 98.30 | 0.9985 | ns |
| 0.01 | 309.9063 | 75.49317 | 8 | 276.2222 | 27.79452634 | 9 | 33.68 | -67.17 to 134.5 | 0.9146 | ns |
| 0.1 | 376.425 | 78.39359 | 8 | 353.4611 | 46.75141561 | 9 | 22.96 | -83.47 to 129.4 | 0.9951 | ns |
| 1 | 524.7063 | 108.3221 | 8 | 483.7111 | 82.35259854 | 9 | 41 | -111.8 to 193.8 | 0.9832 | ns |
| 3 | 561.575 | 111.6077 | 8 | 503.0333 | 99.49906407 | 9 | 58.54 | -106.2 to 223.2 | 0.9236 | ns |
| 10 | 568.1063 | 109.6222 | 8 | 505.8444 | 117.6180562 | 9 | 62.26 | -112.4 to 236.9 | 0.9249 | ns |
| 30 | 518.3563 | 109.6047 | 8 | 435.7 | 123.2357446 | 9 | 82.66 | -96.15 to 261.5 | 0.7612 | ns |
| 100 | 640.475 | 113.8163 | 8 | 575.6167 | 136.9575299 | 9 | 64.86 | -127.9 to 257.6 | 0.9446 | ns |

| ANOVA table | SS | DF | MS | F (DFn, DFd) | P value |
|---|---|---|---|---|---|
| Intensity x Genotype | 15922 | 7 | 2275 | F (7, 105) = 1.043 | P=0.4062 |
| Intensity | 2064871 | 7 | 294982 | F (1.726, 25.89) = 135.2 | P<0.0001 |
| Genotype | 76890 | 1 | 76890 | F (1, 15) = 1.343 | P=0.2647 |
| Subject | 859019 | 15 | 57268 | F (15, 105) = 26.25 | P<0.0001 |
| Residual | 229079 | 105 | 2182 | | |

**Supplemental Table S4.3 related to Figure S4C: a-wave time to peak (ms)**

| | Cone-Control | | | Cone-HCN1 KO | | | Cone-Control - Cone HCN1 KO | | | |
|---|---|---|---|---|---|---|---|---|---|---|
| $cd.s/m^2$ | mean | st dev | N | mean | st dev | N | mean | 95% CI | adj p | significant |
| 0.003 | 21.625 | 4.210955 | 8 | 22.77778 | 5.287728193 | 9 | -1.153 | -8.470 to 6.164 | 0.9996 | ns |
| 0.01 | 22.625 | 1.052209 | 8 | 22.69444 | 1.483122644 | 9 | -0.06944 | -2.042 to 1.903 | >0.9999 | ns |
| 0.1 | 18.5625 | 1.006674 | 8 | 18.44444 | 1.059022086 | 9 | 0.1181 | -1.471 to 1.707 | >0.9999 | ns |
| 1 | 15.6875 | 1.058554 | 8 | 15.44444 | 1.044163674 | 9 | 0.2431 | -1.380 to 1.866 | 0.9997 | ns |
| 3 | 11.15625 | 2.82823 | 8 | 10.16667 | 1.17260394 | 9 | 0.9896 | -2.789 to 4.768 | 0.9783 | ns |
| 10 | 8.65625 | 0.94432 | 8 | 8.333333 | 0.661437828 | 9 | 0.3229 | -0.9876 to 1.633 | 0.9896 | ns |
| 30 | 7.40625 | 0.789863 | 8 | 7.166667 | 0.414578099 | 9 | 0.2396 | -0.8226 to 1.302 | 0.9927 | ns |
| 100 | 6.125 | 0.377964 | 8 | 5.805556 | 0.300462606 | 9 | 0.3194 | -0.2196 to 0.8585 | 0.4751 | ns |

| ANOVA table | SS | DF | MS | F (DFn, DFd) | P value |
|---|---|---|---|---|---|
| Intensity x Genotype | 10.68 | 7 | 1.526 | F (7, 105) = 0.4751 | P=0.8507 |
| Intensity | 5354 | 7 | 764.8 | F (1.382, 20.73) = 238.1 | P<0.0001 |
| Genotype | 0.5405 | 1 | 0.5405 | F (1, 15) = 0.05308 | P=0.8209 |
| Subject | 152.7 | 15 | 10.18 | F (15, 105) = 3.171 | P=0.0003 |
| Residual | 337.2 | 105 | 3.212 | | |

**Supplemental Table S4.4 related to Figure S4C: b-wave time to peak (ms)**

| | Cone-Control | | | Cone-HCN1 KO | | | Cone-Control - Cone HCN1 KO | | | |
|---|---|---|---|---|---|---|---|---|---|---|
| $cd.s/m^2$ | mean | st dev | N | mean | st dev | N | mean | 95% CI | adj p | significant |
| 0.003 | 60.125 | 2.793487 | 8 | 56.63889 | 7.150296225 | 9 | 3.486 | -5.245 to 12.22 | 0.8399 | ns |
| 0.01 | 41.59375 | 3.793645 | 8 | 39.13889 | 2.834252302 | 9 | 2.455 | -2.877 to 7.787 | 0.7489 | ns |
| 0.1 | 29.125 | 1.913299 | 8 | 28.72222 | 1.834753238 | 9 | 0.4028 | -2.497 to 3.303 | 0.9998 | ns |
| 1 | 27.1875 | 1.585368 | 8 | 26.77778 | 1.817641176 | 9 | 0.4097 | -2.204 to 3.023 | 0.9996 | ns |
| 3 | 26.0625 | 2.671777 | 8 | 25.80556 | 1.967619826 | 9 | 0.2569 | -3.487 to 4.001 | >0.9999 | ns |
| 10 | 27.5 | 1.899248 | 8 | 27.47222 | 1.839006374 | 9 | 0.02778 | -2.862 to 2.918 | >0.9999 | ns |
| 30 | 28.59375 | 2.030735 | 8 | 28.94444 | 2.42634556 | 9 | -0.3507 | -3.776 to 3.075 | >0.9999 | ns |
| 100 | 26.34375 | 1.523257 | 8 | 26.33333 | 1.489546911 | 9 | 0.01042 | -2.317 to 2.338 | >0.9999 | ns |

| ANOVA table | SS | DF | MS | F (DFn, DFd) | P value |
|---|---|---|---|---|---|
| Intensity x Genotype | 55.45 | 7 | 7.921 | F (7, 105) = 1.527 | P=0.1662 |
| Intensity | 15143 | 7 | 2163 | F (1.303, 19.55) = 417.0 | P<0.0001 |
| Genotype | 23.75 | 1 | 23.75 | F (1, 15) = 0.8408 | P=0.3737 |
| Subject | 423.7 | 15 | 28.25 | F (15, 105) = 5.445 | P<0.0001 |
| Residual | 544.7 | 105 | 5.188 | | |

**Supplemental Table S4.5 related to Figure S4D: amplitude 100 ms post flash (μV)**

| | Cone-Control | | | Cone-HCN1 KO | | | Cone-Control - Cone HCN1 KO | | | |
|---|---|---|---|---|---|---|---|---|---|---|
| $cd.s/m^2$ | mean | st dev | N | mean | st dev | N | mean | 95% CI | adj p | significant |
| 0.003 | 284.5891 | 77.73666 | 8 | 294.3811 | 53.10605706 | 9 | -9.792 | -117.2 to 97.63 | >0.9999 | ns |
| 0.01 | 282.0157 | 78.94025 | 8 | 273.5202 | 32.54037788 | 9 | 8.495 | -96.97 to 114.0 | >0.9999 | ns |
| 0.1 | 141.2729 | 53.66448 | 8 | 156.5187 | 50.99665917 | 9 | -15.25 | -96.30 to 65.81 | 0.9986 | ns |
| 1 | 141.3355 | 52.88599 | 8 | 157.11 | 48.0117836 | 9 | -15.77 | -94.31 to 62.76 | 0.9977 | ns |
| 3 | 136.5271 | 45.75338 | 8 | 137.0364 | 24.78441848 | 9 | -0.5093 | -62.15 to 61.13 | >0.9999 | ns |
| 10 | 132.3671 | 48.22781 | 8 | 144.8666 | 27.26955151 | 9 | -12.5 | -77.67 to 52.67 | 0.9977 | ns |
| 30 | 121.0221 | 60.87722 | 8 | 110.9323 | 24.81165566 | 9 | 10.09 | -71.24 to 91.42 | 0.9999 | ns |
| 100 | 192.0591 | 66.00147 | 8 | 198.9009 | 65.85965711 | 9 | -6.842 | -108.5 to 94.86 | >0.9999 | ns |

| ANOVA table | SS | DF | MS | F (DFn, DFd) | P value |
|---|---|---|---|---|---|
| Intensity x Genotype | 3105 | 7 | 443.6 | F (7, 105) = 0.4790 | P=0.8480 |
| Intensity | 531284 | 7 | 75898 | F (4.364, 65.47) = 81.95 | P<0.0001 |
| Genotype | 937.3 | 1 | 937.3 | F (1, 15) = 0.05944 | P=0.8107 |
| Subject | 236556 | 15 | 15770 | F (15, 105) = 17.03 | P<0.0001 |
| Residual | 97241 | 105 | 926.1 | | |

**Supplemental Table S4.6 related to Figure S4D: amplitude 150 ms post flash (μV)**

| | Cone-Control | | | Cone-HCN1 KO | | | Cone-Control - Cone HCN1 KO | | | |
|---|---|---|---|---|---|---|---|---|---|---|
| $cd.s/m^2$ | mean | st dev | N | mean | st dev | N | mean | 95% CI | adj p | significant |
| 0.003 | 128.095 | 45.07987 | 8 | 136.5962 | 48.91527713 | 9 | -8.501 | -80.72 to 63.71 | >0.9999 | ns |
| 0.01 | 124.5165 | 56.25516 | 8 | 132.0966 | 32.20026246 | 9 | -7.58 | -83.68 to 68.52 | >0.9999 | ns |
| 0.1 | 43.53123 | 26.56132 | 8 | 68.28645 | 46.06745668 | 9 | -24.76 | -83.14 to 33.63 | 0.8187 | ns |
| 1 | 50.05368 | 34.5764 | 8 | 63.32307 | 32.29228626 | 9 | -13.27 | -65.15 to 38.61 | 0.9886 | ns |
| 3 | 51.88838 | 27.67896 | 8 | 52.53023 | 22.2511979 | 9 | -0.6418 | -40.23 to 38.95 | >0.9999 | ns |
| 10 | 44.7359 | 36.95681 | 8 | 63.28604 | 29.7710138 | 9 | -18.55 | -71.44 to 34.34 | 0.9261 | ns |
| 30 | 61.10897 | 50.43681 | 8 | 44.36639 | 21.39502151 | 9 | 16.74 | -50.66 to 84.14 | 0.9844 | ns |
| 100 | 146.3735 | 59.83724 | 8 | 157.5199 | 56.8131143 | 9 | -11.15 | -101.5 to 79.20 | >0.9999 | ns |

| ANOVA table | SS | DF | MS | F (DFn, DFd) | P value |
|---|---|---|---|---|---|
| Intensity x Genotype | 4637 | 7 | 662.4 | F (7, 105) = 0.7345 | P=0.6431 |
| Intensity | 225674 | 7 | 32239 | F (3.364, 50.46) = 35.75 | P<0.0001 |
| Genotype | 2427 | 1 | 2427 | F (1, 15) = 0.3431 | P=0.5668 |
| Subject | 106087 | 15 | 7072 | F (15, 105) = 7.843 | P<0.0001 |
| Residual | 94684 | 105 | 901.8 | | |

| Supplemental Table S4.7 related to Figure S4D: amplitude 200 ms post flash (µV) | | | | | | | | | | |
|---|---|---|---|---|---|---|---|---|---|---|
| | Cone-Control | | | Cone-HCN1 KO | | | Cone-Control - Cone HCN1 KO | | | |
| cd.s/m$^2$ | mean | st dev | N | mean | st dev | N | mean | 95% CI | adj p | significant |
| 0.003 | 19.00799 | 26.65084 | 8 | 20.75132 | 20.0084774 | 9 | -1.743 | -39.24 to 35.75 | >0.9999 | ns |
| 0.01 | 50.26174 | 31.05355 | 8 | 59.58842 | 28.98438188 | 9 | -9.327 | -55.91 to 37.26 | 0.9978 | ns |
| 0.1 | 2.724442 | 7.811548 | 8 | 25.14469 | 36.09846853 | 9 | -22.42 | -66.26 to 21.42 | 0.5822 | ns |
| 1 | 15.4148 | 21.89522 | 8 | 25.02429 | 20.88031065 | 9 | -9.609 | -42.73 to 23.51 | 0.9755 | ns |
| 3 | 26.27001 | 19.35118 | 8 | 31.81418 | 18.80871976 | 9 | -5.544 | -35.04 to 23.95 | 0.9986 | ns |
| 10 | 27.60179 | 30.47292 | 8 | 40.83074 | 19.21162393 | 9 | -13.23 | -54.86 to 28.40 | 0.9502 | ns |
| 30 | 37.20325 | 35.7941 | 8 | 20.53761 | 22.44644712 | 9 | 16.67 | -32.20 to 65.53 | 0.9278 | ns |
| 100 | 108.4484 | 58.40274 | 8 | 113.6394 | 34.48704956 | 9 | -5.191 | -84.41 to 74.03 | >0.9999 | ns |

| ANOVA table | SS | DF | MS | F (DFn, DFd) | P value |
|---|---|---|---|---|---|
| Intensity x Genotype | 3718 | 7 | 531.2 | F (7, 105) = 0.8146 | P=0.5772 |
| Intensity | 118845 | 7 | 16978 | F (2.884, 43.26) = 26.03 | P<0.0001 |
| Genotype | 1345 | 1 | 1345 | F (1, 15) = 0.6272 | P=0.4407 |
| Subject | 32162 | 15 | 2144 | F (15, 105) = 3.288 | P=0.0002 |
| Residual | 68474 | 105 | 652.1 | | |

| Supplemental Table S5 related to Figure S5B: 30Hz flicker amplitude (µV) | | | | | | | | | | |
|---|---|---|---|---|---|---|---|---|---|---|
| | Cone-Control | | | Cone-HCN1 KO | | | Cone-Control - Cone HCN1 KO | | | |
| cd.s/m$^2$ | mean | st dev | N | mean | st dev | N | mean | 95% CI | adj p | significant |
| 3 | 50.16907 | 10.49319 | 7 | 50.17595 | 11.41600902 | 11 | -0.006883 | -16.11 to 16.10 | >0.9999 | ns |
| 10 | 38.43914 | 9.222136 | 7 | 36.29586 | 8.667997901 | 11 | 2.143 | -11.48 to 15.76 | 0.9975 | ns |
| 30 | 33.83314 | 8.499501 | 7 | 30.68332 | 8.031908193 | 11 | 3.15 | -9.418 to 15.72 | 0.9718 | ns |
| 100 | 35.15943 | 4.880929 | 7 | 32.57682 | 7.273965807 | 11 | 2.583 | -6.018 to 11.18 | 0.9437 | ns |
| 300 | 35.15936 | 4.529089 | 7 | 33.05127 | 6.818241318 | 11 | 2.108 | -5.918 to 10.13 | 0.9699 | ns |
| 724 | 26.04307 | 4.825053 | 7 | 24.05682 | 5.54671159 | 11 | 1.986 | -5.557 to 9.530 | 0.9676 | ns |

| ANOVA table | SS | DF | MS | F (DFn, DFd) | P value |
|---|---|---|---|---|---|
| Frequency x Genotype | 24.47 | 5 | 4.895 | F (5, 80) = 0.3975 | P=0.8492 |
| Frequency | 5871 | 5 | 1174 | F (1.980, 31.69) = 95.34 | P<0.0001 |
| Genotype | 102 | 1 | 102 | F (1, 16) = 0.3248 | P=0.5766 |
| Subject | 5026 | 16 | 314.1 | F (16, 80) = 25.51 | P<0.0001 |
| Residual | 985.2 | 80 | 12.31 | | |

**Supplemental Table S6 related to Figure S6: amplitude of fundamental (μV)**

| | Cone-Control | | | Cone-HCN1 KO | | | Cone-Control - Cone HCN1 KO | | | |
|---|---|---|---|---|---|---|---|---|---|---|
| Hz | mean | st dev | N | mean | st dev | N | mean | 95% CI | adj p | significant |
| 0.5 | 11.6717 | 3.11143 | 6 | 15.62098 | 4.306244592 | 6 | -3.949 | -12.03 to 4.127 | 0.6922 | ns |
| 0.75 | 10.47132 | 1.903116 | 6 | 14.51195 | 3.864791995 | 6 | -4.041 | -11.11 to 3.029 | 0.4554 | ns |
| 1 | 9.708163 | 2.176626 | 6 | 10.18622 | 1.790571029 | 6 | -0.4781 | -4.691 to 3.735 | >0.9999 | ns |
| 1.5 | 9.373112 | 1.11526 | 6 | 10.13073 | 1.431333002 | 6 | -0.7576 | -3.487 to 1.971 | 0.9882 | ns |
| 2 | 8.936081 | 1.219023 | 6 | 9.85243 | 2.141034822 | 6 | -0.9163 | -4.834 to 3.001 | 0.9956 | ns |
| 3 | 8.144978 | 1.319111 | 6 | 9.28527 | 1.662469787 | 6 | -1.14 | -4.325 to 2.045 | 0.934 | ns |
| 6 | 7.851962 | 1.142144 | 6 | 8.666592 | 1.096221284 | 6 | -0.8146 | -3.158 to 1.529 | 0.9484 | ns |
| 9 | 9.222809 | 0.956109 | 6 | 10.12186 | 1.274871286 | 6 | -0.8991 | -3.308 to 1.510 | 0.9134 | ns |
| 12 | 7.657157 | 0.552469 | 6 | 8.112055 | 0.909840018 | 6 | -0.4549 | -2.125 to 1.215 | 0.9867 | ns |
| 15 | 5.966479 | 0.785425 | 6 | 6.486527 | 0.83324245 | 6 | -0.52 | -2.216 to 1.176 | 0.9776 | ns |
| 25 | 2.553902 | 0.433825 | 6 | 3.372081 | 0.391419634 | 6 | -0.8182 | -1.685 to 0.04896 | 0.0696 | ns |

| ANOVA table | SS | DF | MS | F (DFn, DFd) | P value |
|---|---|---|---|---|---|
| Frequency x Genotype | 52.8 | 10 | 5.28 | F (10, 100) = 2.497 | P=0.0102 |
| Frequency | 982.3 | 10 | 98.23 | F (1.921, 19.21) = 46.46 | P<0.0001 |
| Genotype | 59.65 | 1 | 59.65 | F (1, 10) = 3.495 | P=0.0911 |
| Subject | 170.7 | 10 | 17.07 | F (10, 100) = 8.071 | P<0.0001 |
| Residual | 211.5 | 100 | 2.115 | | |

**Supplemental Table S7: related to Figure S7: OMR contrast sensitivity (70 cd/m2)**

| | Rod-Control | | | Rod-HCN1 KO | | | Rod-Control - Rod-HCN1 KO | | | |
|---|---|---|---|---|---|---|---|---|---|---|
| Temporal Frequency (Hz) | log mean | st dev | N | log mean | st dev | N | mean | t | adj p | significant |
| 1.5 | 0.35162 | 0.081333 | 5 | 0.324125 | 0.065139715 | 5 | 0.024 | 0.324 | 0.748 | ns |
| 3 | 0.52292 | 0.142427 | 5 | 0.451525 | 0.152631351 | 5 | 0.0585 | 0.79 | 0.437 | ns |
| 6 | 0.35058 | 0.158393 | 5 | 0.3343 | 0.107631873 | 5 | 0.021 | 0.284 | 0.779 | ns |

| ANOVA table | SS | DF | MS | F | P value |
|---|---|---|---|---|---|
| Intensity x frequency | 2 | 0.00217 | 0.00108 | 0.079 | 0.924 |
| Frequency | 2 | 0.158 | 0.0789 | 5.748 | 0.009 |
| Genotype | 1 | 0.00894 | 0.00894 | 0.652 | 0.427 |
| Residual | 24 | 0.329 | 0.0137 | | |
| Total | 29 | 0.498 | 0.0172 | | |
